# Leukocyte Immunoglobulin-Like Receptor B1 and its Interactions with Human Leukocyte Antigens

**DOI:** 10.64898/2026.08.16.745109

**Authors:** Geoff X.Y. Zhang, Jia Q. Truong, Lucy Sullivan, Megan Lake, Timothy Emery, James Roest, Ashley J. Ovens, Muhammad N.H. Khabib, Meiqi Cao, Benjamin R. Turner, Alexander D. Barrow, Jessica K. Holien, Julian P. Vivian, Christopher G. Langendorf

## Abstract

Interactions between Human Leukocyte Antigen (HLA) molecules and their cognate immunoreceptors are essential for regulating innate and adaptive immune cell functions. Leukocyte Immunoglobulin-like Receptors (LILRs) are key regulators of HLA-mediated immune responses, owing to their broad expression across immune cell populations and their ability to modulate both immune activation and tolerance. Among these, LILRB1-HLA interactions are increasingly recognised as important in transplantation, chronic infection and cancer therapies. Unlike other HLA-binding receptors, which recognise epitopes specific to HLA subsets, LILRB1 primarily engages the relatively conserved α3 and β2-microglobulin components of HLA molecules, supporting its role as a broad regulator of pan-HLA class I-mediated functions. Nonetheless, there have been conflicting findings regarding the breadth of LILRB1-HLA-I interactions. While direct affinity studies on a limited subset of HLA-I molecules have revealed no significant differences in LILRB1 binding, broader analyses using single-antigen bead arrays suggest underlying variability. Here, we show through a broad binding assay that, while LILRB1 is a broad HLA-I-binding receptor, it exhibits differential preferences across HLA-I allotypes. Molecular dynamics analyses of the HLA-I-LILRB1 interface suggest that HLA-α3 domain dynamism underlies these binding differences. We further determined the crystal structure of LILRB1 and used it to highlight intrinsic structural flexibility within its domains. Finally, these structural insights were leveraged to refine our understanding of the binding modalities of therapeutic monoclonal antibodies currently described. Together, our findings establish structural and mechanistic bases for differential HLA-I recognition by LILRB1 and provide insights into immunotherapeutic targeting of LILRB1.

**Significance statement:** Leukocyte immunoglobulin-like receptor B1 (LILRB1) is a widely expressed inhibitory immune receptor that functions through interactions with human leukocyte antigen (HLA) class I molecules. By combining broad HLA binding assays, molecular dynamics simulations, and structural analyses, we show how domain flexibility and HLA stability shape LILRB1-HLA interactions and antibody recognition. These insights improve the mechanistic understanding of LILRB1–HLA interactions and inform strategies for therapeutic targeting of LILRB1.

## 1 Introduction

Leukocyte immunoglobulin-like receptor B1 (LILRB1), also known as immunoglobulin-like transcript 2 (ILT2) or CD85j, is an inhibitory immune receptor with an extracellular region composed of four immunoglobulin-like domains (D1-D4) linked by flexible hinge regions (Zhang et al., 2025). LILRB1 is the most broadly expressed member of the LILRB-family, present on both adaptive and innate immune cell populations and has been implicated in multiple facets of immune modulation (Zhang et al., 2025, Deng et al., 2021, Zeller et al., 2023). In adaptive immunity, LILRB1 contributes to the inhibition of T- and B-cell responses (Dietrich et al., 2001, Naji et al., 2014). In innate immune cells, it suppresses macrophage phagocytosis and regulates the inhibition and education of natural killer (NK) cells (Barkal et al., 2018, Leijonhufvud et al., 2021). Given the breadth of its immunoregulatory involvement, the LILRB1 immune checkpoint is emerging as an attractive therapeutic target, exemplified by the development and ongoing clinical evaluation of numerous anti-LILRB1 antibodies (Zhang et al., 2025, Zhao et al., 2019).

The canonical ligands of LILRB1 are the Human Leukocyte Antigen class I (HLA-I) molecules, cell-surface proteins that present intracellular peptides for immune surveillance and tolerance (Zhang et al., 2025, Fukazawa et al., 1994). The structure of HLA-I molecules consists of a heavy chain, comprising α1, α2, and α3 domains, non-covalently associated with the monomorphic β2-microglobulin (β2M) subunit (Zhang et al., 2025, Fukazawa et al., 1994). HLA-I molecules can be categorized into highly polymorphic classical HLA-I molecules, including over 20,000 alleles of HLA-A, HLA-B, and HLA-C, and the less variable non-classical HLA-I molecules, HLA-E, HLA-F, and HLA-G (Robinson et al., 2020). Despite this extensive polymorphism, LILRB1 exhibits broad HLA-I binding capability due to its mode of interaction (Zhang et al., 2025). Whereas sequence variability in the classical HLA-I largely manifests around the α1 and α2 domains near the peptide-binding groove, LILRB1 binds to the relatively conserved α3 domain and β2M subunit (Zhang et al., 2025, Pymm et al., 2024).

Crystallographic structures of LILRB1 in complex with HLA molecules have provided molecular insight into the receptor-ligand interface and the structural basis of HLA recognition (Zhang et al., 2025). The structure of truncated LILRB1 D1-D2 bound to HLA-A*02 elucidated that binding is mediated through 12 residues in D1 interacting with α3 and β2M of HLA-A*02, and 7 residues in D2 interacting with the β2M alone (Willcox et al., 2003). Conversely, binding studies using truncated D3-D4-LILRB1 across a range of HLA-I allotypes, including HLA-B*2702, HLA-Cw*0702, HLA-G1, and HLA-E, observed no measurable interactions (Chapman et al., 1999). Subsequent structures of full-length LILRB1 in complex with HLA-G demonstrated a conserved binding mode, where D1 and D2 form the interaction interface with α3 and β2M of HLA-G, while D3 and D4 act as a rigid structural scaffold that maintains a staggered domain orientation without directly contacting the HLA (Wang et al., 2020). With the accrual of truncated D1-D2-LILRB1 structures in free and ligand-bound states, as well as full-length ligand-bound forms, it has become increasingly evident that the D1 and D2 domains possess substantial interdomain flexibility and conformational plasticity (Zhang et al., 2025, Willcox et al., 2003, Shiroishi et al., 2006, Wang et al., 2020). This property is thought to underlie, at least in part, the ability of LILRB1 to bind a diverse range of HLA molecules and supports the model that interactions between LILRB1 and HLA are entropically driven (Zhang et al., 2025, Shiroishi et al., 2006, Kuroki et al., 2019).

Despite the relatively conserved nature of the HLA α3 domain and β2M and the conformational flexibility of LILRB1 domains D1 and D2, differential binding of LILRB1 to classical HLA-I allotypes has been reported, albeit contentiously. Early surface plasmon resonance (SPR) studies characterising the interactions between LILRB1 and select HLA-I allotypes, including HLA-A*11, HLA-B*27, HLA-B*35, and several HLA-C allotypes, observed little to no difference in binding affinity (Chapman et al., 1999, Shiroishi et al., 2003, Kuroki et al., 2005). However, subsequent attempts to assess the uniformity of HLA-I affinity across a broad panel of cell-derived HLA molecules reported allotype-dependent differences in LILRB1 binding in single-antigen bead experiments. The first such study, by Jones et al., used full-length LILRB1 Fc constructs and identified differences in affinity among HLA-A allotypes that correlated with polymorphisms at residues 193, 194, 207, 246, and 253 within the α3 domain (Jones et al., 2011). Conversely, a later study by Liu et al. using D1-D2-LILRB1 Fc constructs reported that affinity differences among HLA-A allotypes were associated only with polymorphisms at residues 207 and 253 (Liu et al., 2022). Additionally, Liu et al. observed differential binding among HLA-B allotypes linked to polymorphisms at residue 194 of the α3 domain, as well as generally weaker binding to HLA-C allotypes (Liu et al., 2022). Consequently, the extent to which LILRB1 discriminates among classical HLA-I alleles, and the underlying basis for these discrepant observations, remains unresolved.

In this study, we sought to clarify some of the ambiguities surrounding the HLA binding preferences of LILRB1 by conducting a broad HLA single-antigen bead screening assay using four extracellular immunoglobulin domain LILRB1 tetramers (D1-D4). This approach indicated that LILRB1 binds HLA allotypes differentially, both across the full panel and within the HLA-A, -B, and -C groups. However, when categorised by polymorphisms in α3 contact residues, binding differences were only observed for a subset of HLA-B allotypes.

Molecular dynamics simulations comparing representative strong and weak binders suggest that these affinity preferences may be attributed to differences in HLA domain dynamism. We further determined the 3.0 Å crystal structure of unbound four extracellular domains LILRB1 and used it as a reference to analyse domain hinge angles across available LILRB1 structures. These analyses revealed flexibility in the D1-D2 hinge region that is further exaggerated upon ligand binding, implying the fundamental role of dynamism in the interactions between LILRB1 and HLA. Additionally, structural modelling of currently available anti-LILRB1 antibodies showed that their epitopes localise not only to the D1-D2 hinge but also to the D3-D4 hinge, implying possible mechanisms for modulating LILRB1 signalling beyond simply mimicking or blocking HLA engagement. Collectively, these findings advance our understanding of the structural and dynamic mechanisms underlying LILRB1-HLA interactions, which will advance a framework for the therapeutic targeting of this innate and adaptive immune checkpoint.

## 2 Results

### 2.1 Broad Screening of HLA-I allotypes Reveals Variation in LILRB1 Binding

We assessed LILRB1 binding to classical HLA-I molecules using LILRB1 tetramers and a panel of 97 common HLA-A, HLA-B, and HLA-C allotypes immobilised on single-antigen beads (see Figure 1A for full list). The mean fluorescence intensity (MFI) of LILRB1 tetramers, normalised to the MFI of the pan-HLA monoclonal antibody W6/32, was used as a proxy for LILRB1 binding (see Supplementary Figure 1 for unnormalised data).

**Figure 1:**
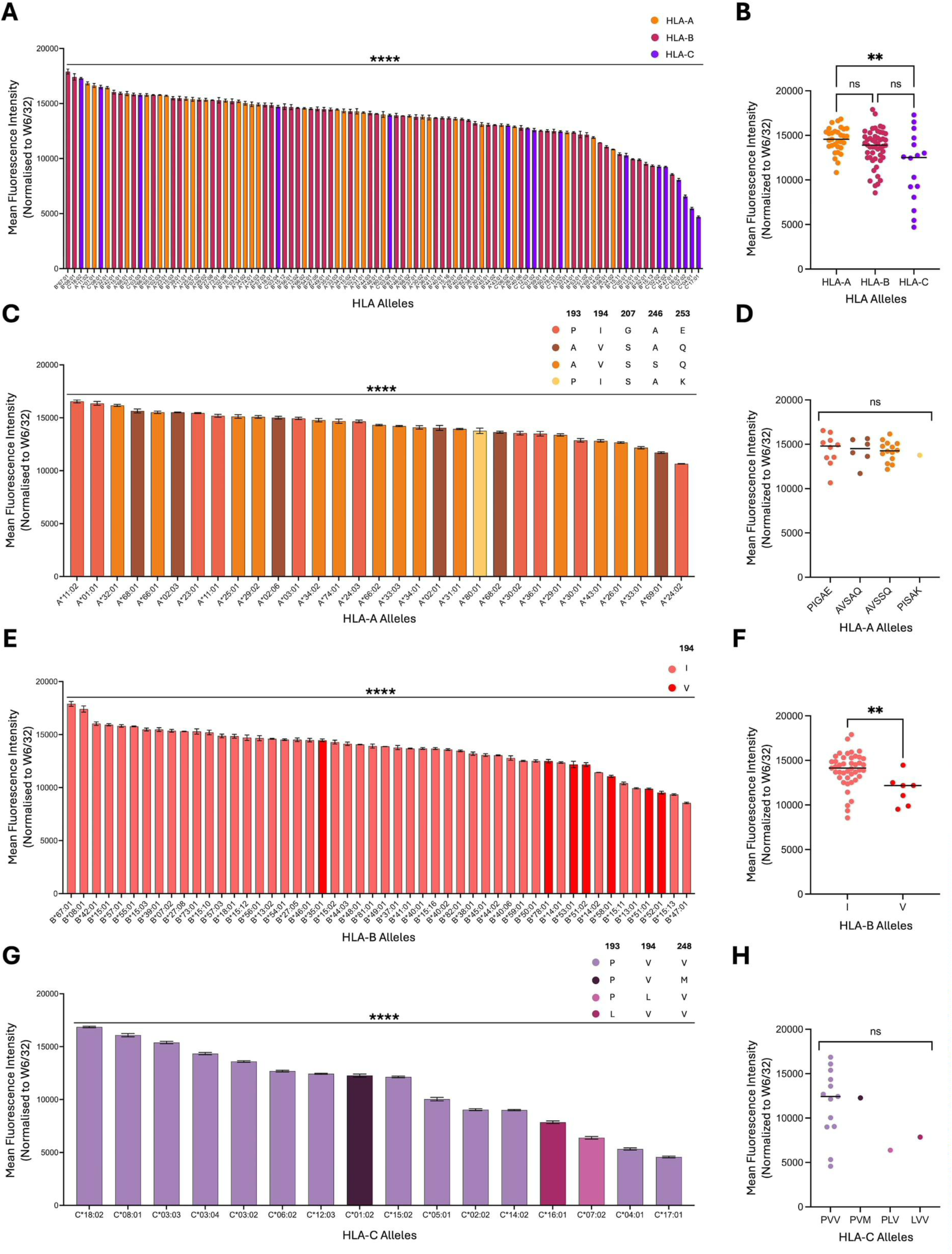
Interactions between LILRB1 and class I HLAs. All data are presented as mean fluorescence intensity (MFI) of LILRB1 tetramer binding normalised to the pan-HLA monoclonal antibody W6/32. **(A)** Comparison of LILRB1 tetramer binding to 97 HLA allotypes immobilised on single-antigen beads. **(B)** LILRB1 binding across 31 HLA-A allotypes (orange), 50 HLA-B allotypes (magenta), and 16 HLA-C allotypes (purple). **(C)** Comparison of binding of LILRB1 tetramers across the 31 HLA-A allotypes; bars are coloured according to specific sequence motifs at positions 193, 194, 207, 246, and 253: vermillion for PIGAE, orange for AVSAQ, brown for AVSSQ, and yellow for PISAK. **(D)** LILRB1 binding to HLA-A allotypes grouped by these motif categories. **(E)** Comparison of binding of LILRB1 tetramers to 50 common HLA-B allotypes; bars are coloured by the residue at position 194: pink for I and red for V. **(F)** LILRB1 binding to HLA-B allotypes grouped according to residue 194. **(G)** Binding of LILRB1 tetramers to 16 common HLA-C allotypes; bars are coloured according to residues at positions 193, 194, and 248: lilac for PVV, dark purple for PVM, red violet for PLV, and light magenta for LVV. **(H)** LILRB1 binding to HLA-C allotypes grouped by these motif categories. Bars represent mean values from three replicates ± SEM. Statistical analyses were performed using the Mann-Whitney U test or Kruskal-Wallis test; *p < 0.05, **p < 0.01, ***p < 0.001, ****p < 0.0001.

When comparing the normalised data across the HLA allotypes, we observed significant differences in LILRB1 binding (Figure 1A). When categorised into HLA-A, -B, or -C, we observed significantly lower mean binding to HLA-C compared to HLA-A, while no significant differences were detected between HLA-A and HLA-B or between HLA-B and HLA-C (Figure 1A and 1B).

Concentrating on HLA-A, we observed statistically significant binding differences between the 31 allotypes (Figure 1C). We next examined whether differences in LILRB1 binding could be attributed to specific residue polymorphisms within the HLA-A family. LILRB1 interacts with the β2M and α3 domain of HLAs. As the β2M is conserved, we focused on polymorphisms occurring in the α3 domain. HLA-A was segregated into allotype groups based on residues at positions 193 and 194, which represent the only polymorphic residues within the LILRB1 binding interface on the α3 domain (Supplementary Figure 2). Further segregation was based on non-contact residues 207, 246 and 253, which have also been previously reported to influence the affinity of LILRB1 for HLA-A (Jones et al., 2011).

Based on the combinations of residues at these positions, HLA-A allotypes were classified into four sequence-motif groups: PIGAE, AVSAQ, AVSSQ, and PISAK (Figure 1C & D). No significant differences in mean LILRB1 binding were observed between these HLA-A groups (Figure 1D), indicating that differences between HLA-A allotype binding to LILRB1 is not due to these previously reported sequence differences.

A similar strategy was employed for HLA-B, where we also observed binding differences between the 50 allotypes (Figure 1E & 1F). Since no α3 domain residue polymorphisms have been reported to influence LILRB1 binding, we stratified HLA-B allotypes based on residues within the α3 domain that contact LILRB1, which include residues 193, 194, 195, 196, 198, and 248 (Supplementary Figure 3). Among these, only position 194 varied within the tested HLA-B alleles, with either isoleucine (Ile194) or the conservative polymorphism, valine (Val194) present (Figure 1E). Although the number of alleles differs between polymorphisms (n = 7 for Val194 and n = 44 for Ile194), mean LILRB1 binding was significantly reduced in allotypes with Val194 compared to those with Ile194 (p < 0.01) (Figure 1F). Notably, this same polymorphism in HLA-A allotypes was not correlated with any differences in LILRB1 binding (Supplementary Figure 2). This finding suggests that this single polymorphism may influence the binding between LILRB1 and HLA-B, likely through factors beyond direct binding affinity differences.

For HLA-C, we observed no binding differences between the 16 allotypes when categorised according to polymorphisms that form the LILRB1-interacting interface (Figure 1G and 1H). We applied a similar stratification strategy to HLA-C, for which no α3 domain residues have previously been implicated in modulating LILRB1 binding. Within our panel, polymorphisms were present among α3 residues that contact LILRB1 at positions 193, 194, and 248 (Supplementary Figure 4). Grouping HLA-C allotypes based on combinations at these positions yielded four motifs: PVV, PVM, PLV, and LVV (Figure 1G). No significant differences in LILRB1 binding were observed among these groups (Figure 1H), although this observation is caveated by the small number of representative HLA-C molecules for three of the polymorphism combinations. It should be noted that the single representative HLA-C polymorphisms containing PLV and LVV were in the bottom 4 of all HLA-C binders and well below the mean MFI of PVV containing polymorphisms. A larger panel of HLA-C allotypes, containing a higher number of the PLV, LVV and PVM polymorphisms is required to determine if this region affects LILRB1 binding.

Taken together, these results indicate that LILRB1 acts as an effective pan-HLA binder, binding to all 97 tested HLA allotypes. However, the strength of this interaction varies across the full panel and among HLA-A, HLA-B, and HLA-C alleles. Barring a valine or isoleucine polymorphism at residue 194 in HLA-B allotypes, no other binding differences were associated with polymorphisms in α3 domain residues known to contact LILRB1. Notably, the same substitution at residue 194 in HLA-A allotypes was not linked to binding variation. This suggests that sequence variation within the α3 domain alone is generally insufficient to account for differences in LILRB1 binding, implying that additional structural or contextual factors may contribute.

### 2.2 Comparative molecular dynamics simulations of LILRB1 bound to HLA-B*35:01 with Val194 or Ile194

We utilised molecular dynamics to further investigate the role of HLA polymorphisms on LILRB1 binding. The only identifiable α3-domain residue that correlated with a significant difference in LILRB1 binding was the conservative valine to isoleucine polymorphism, found at residue 194 in HLA-B allotypes (Figure 2; stereo images in Supplementary Figure 5). We used molecular dynamics program GROMACS 2025.3 (Abraham et al., 2015) to perform 100-ns simulations to model LILRB1 binding to HLA-B*35:01 containing either Val194 (Figure 2A) or a point-substituted Ile194 (Figure 2B).

**Figure 2.**
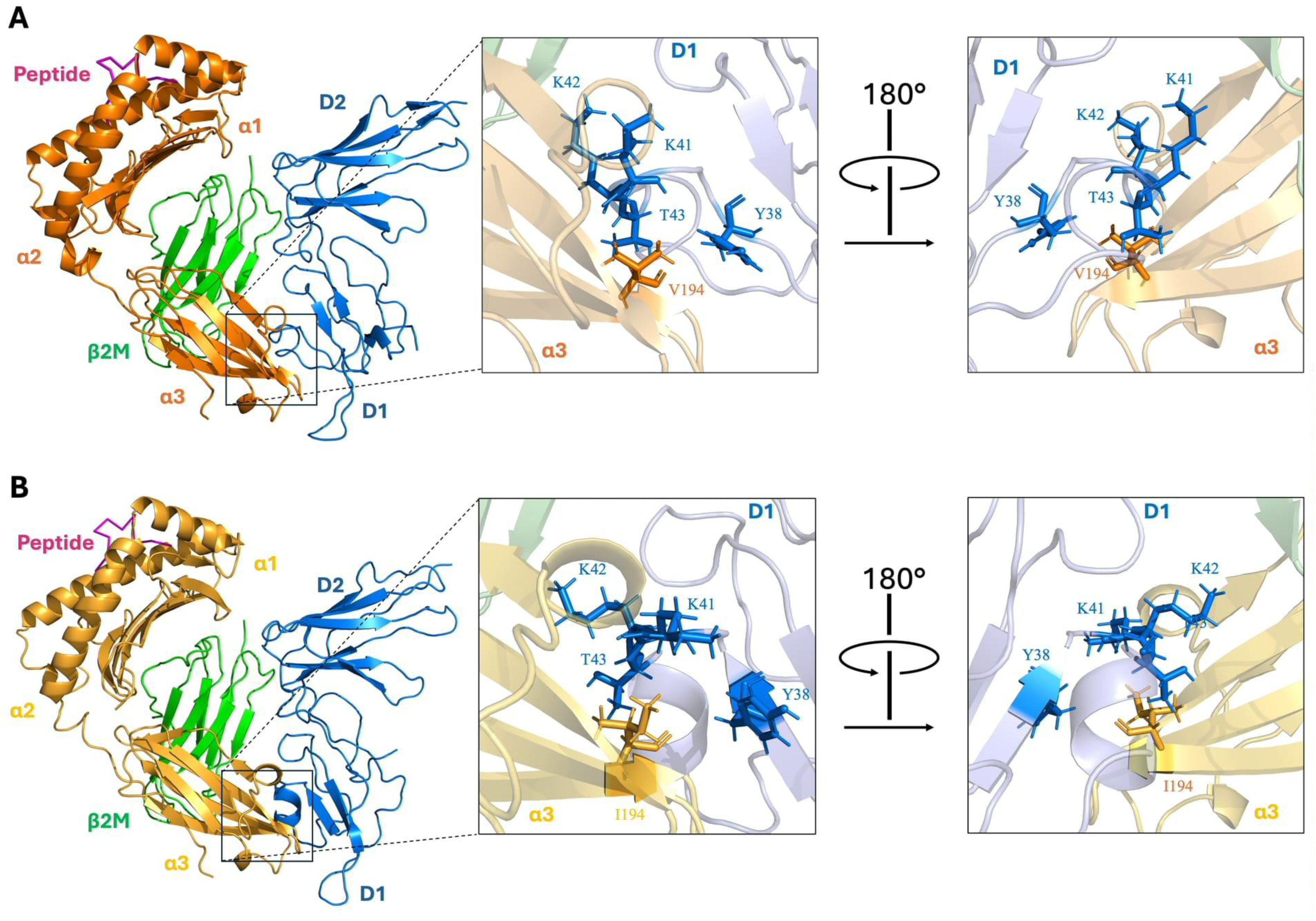
Close-up views of equilibrated structures of LILRB1 interacting with HLA-B*35 at the start of the simulations. Shown are interactions between LILRB1 and HLA-B*35 containing **(A)** valine (orange) or **(B)** isoleucine (yellow) at residue 194. For both, proteins are shown in cartoon representation, with residue 194 of the HLA heavy chain and possible interacting residues on LILRB1 depicted as sticks. The HLA heavy chain is coloured orange, β2M green, the HLA peptide magenta, and LILRB1 blue.

In total, we performed six independent molecular dynamics simulations for each system to compare their conformational behaviour. Analysis of the standard deviations of RMSD values (Figure 3; RMSD across the trajectories is shown in Supplementary Figure 6), reflective of conformational dynamics and regional flexibility, showed higher values for the β2M residues that contact LILRB1 in the HLA-B complex with Ile194 (Figure 3G). This change was not accompanied by differences in the overall protein domains or in the other interacting interfaces (Figure 3), indicating that the Ile194 substitution locally alters the LILRB1-β2M contact interface, despite being located approximately 25 Å from β2M.

**Figure 3.**
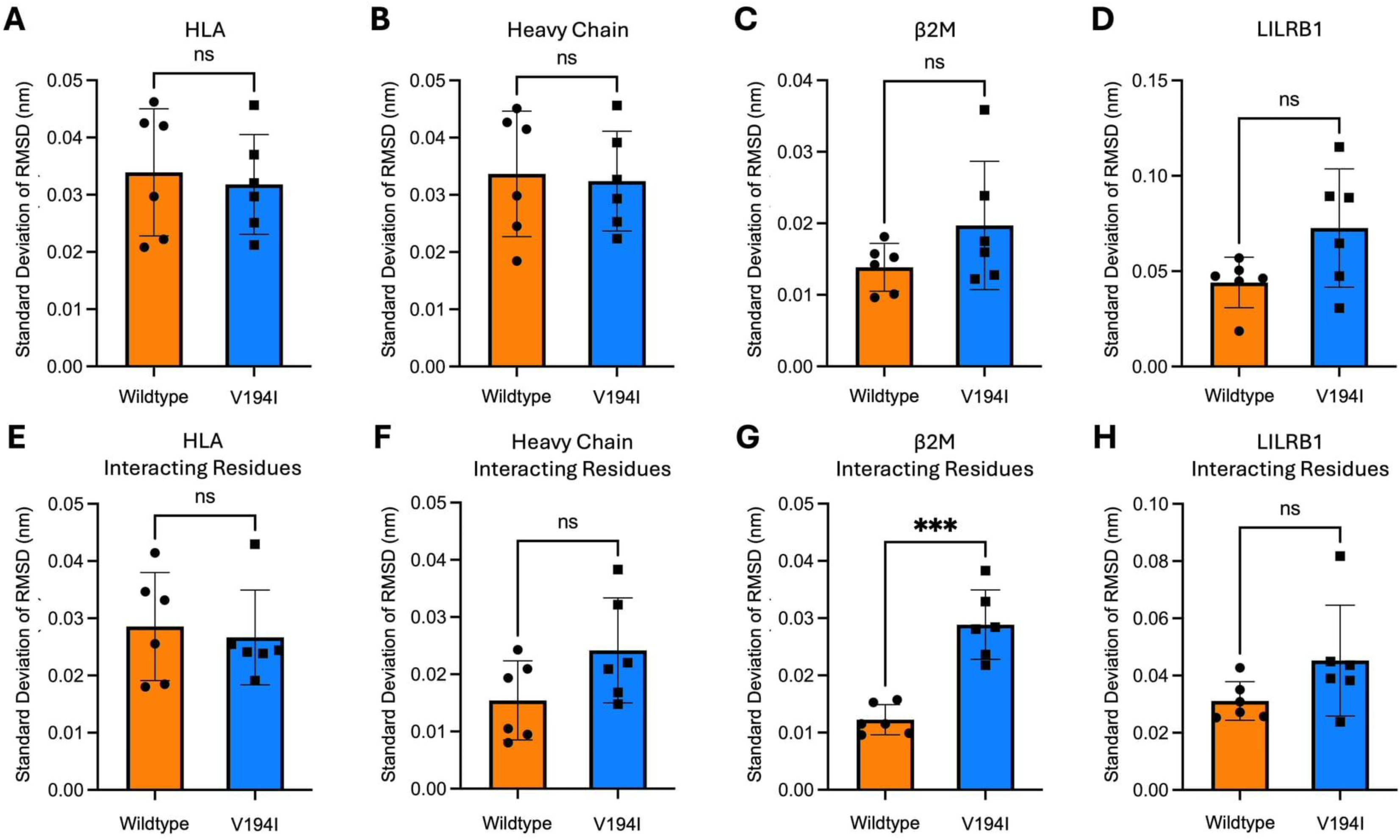
Standard deviations of root mean square deviation (RMSD) values from simulations of LILRB1 in complex with wild-type HLA-B*35:01 (orange) or HLA-B*35:01 containing a valine-to-isoleucine point mutation at residue 194 of the heavy chain (blue). Comparisons are shown for **(A)** the whole HLA complex, **(B)** the heavy chain, **(C)** β2M, **(D)** LILRB1, and for the interacting residues of **(E)** HLA, **(F)** the heavy chain, **(G)** β2M, and **(H)** LILRB1. Interacting residues comprise heavy-chain residues 193-196, 198, and 248; β2M residues 1-4, 86-89, 91-93, and 96; and LILRB1 residues 18, 36, 38, 39, 41-43, 67, 68, 76, 97-100, 125-127, 184, and 187 (Willcox et al., 2003). Data are presented as mean ± SEM from si× 100-ns independent simulations of each system. Statistical analyses were performed using Welch’s t-test; *p<0.05, **p<0.01, ***p<0.001.

Analysis of overall mean RMSD values revealed higher values for the overall HLA molecule with Ile194 (Supplementary Figure 7A), which was primarily attributable to β2M rather than the heavy chain (Supplementary Figure 7C). Comparison of residues within the LILRB1-HLA interface showed higher RMSD values across all residues involved in the interacting interface between LILRB1 and HLA (Supplementary Figure 7E-H). Collectively, these results indicate that, despite their structural similarity, the presence of Ile194 versus Val194 is associated with conformational differences in β2M and the LILRB-HLA contact residues, as well as altered dynamics of the LILRB1-interacting residues on β2M.

These conformational differences were also observed in trajectory snapshots from the 100-ns simulations, aligned using the HLA heavy chain as a reference (Figure 4). Notably, LILRB1 interacting with HLA-B with Val194 had a narrower range of interdomain angles, from 86° to 92° (89.6° ± 2.3°), than the HLA with Ile194, from 78° to 91° (84.6° ± 4.9°) (Figure 4).

**Figure 4.**
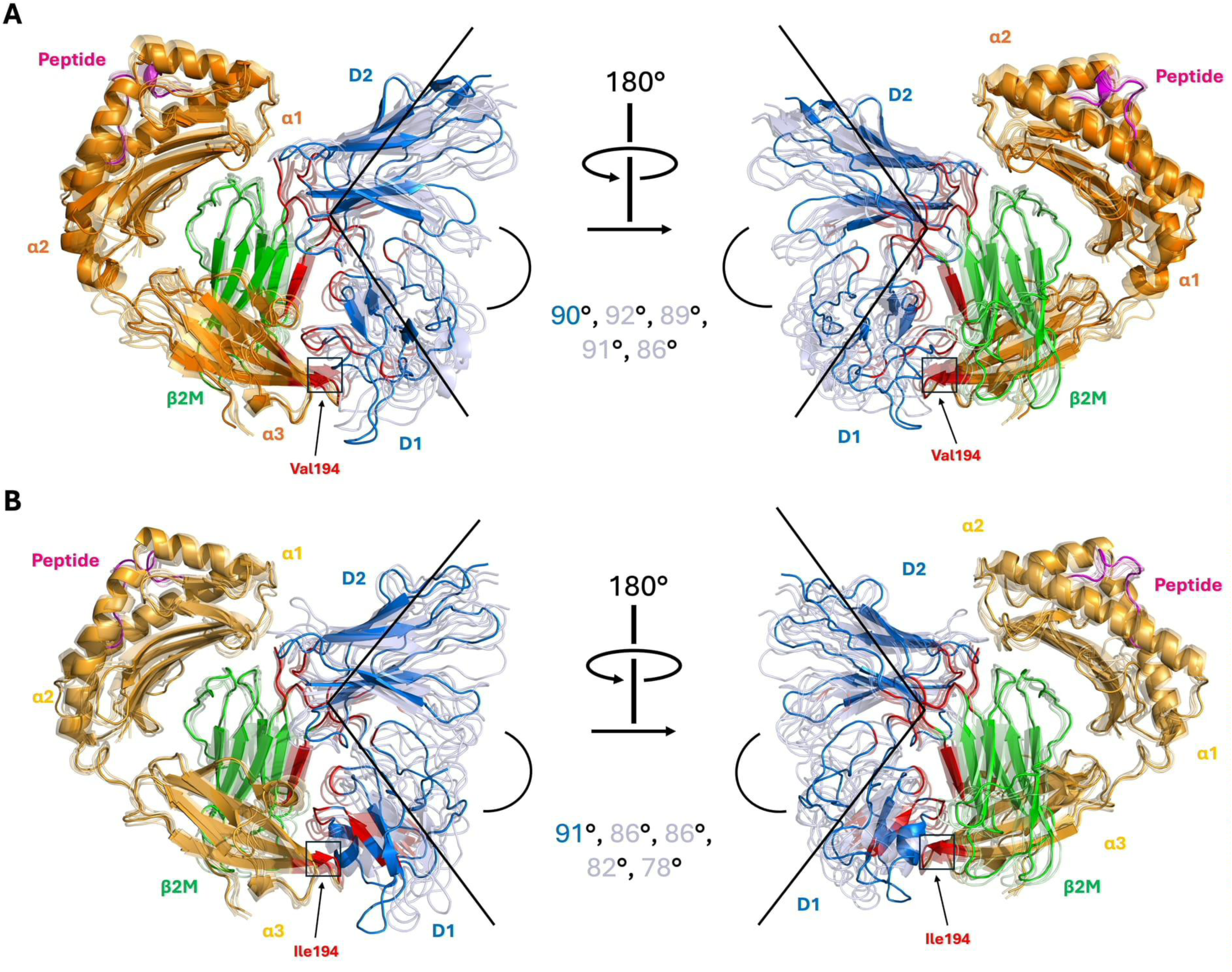
Representative snapshots from 100 ns molecular dynamics simulations of LILRB1 in complex with HLA-B*35. Cartoon representations of two orientations of **(A)** HLA-B*35 containing valine at position 194 (orange) or **(B)** HLA-B*35 containing isoleucine at position 194 (yellow). The HLA heavy chain is coloured in orange, β2M in green, peptide in magenta, and LILRB1 in blue, with residues previously reported to participate in the interaction highlighted in red. For comparison, the heavy chain was used as the alignment anchor across and within each system. The initial structure at 5 ns is shown as the opaque model, while superimposed translucent models represent snapshots taken at 25, 50, 75, and 95 ns of the simulation. Angles measured in the initial structure are depicted with the black lines.

These data suggest that although no differences are evident when LILRB1 conformations are averaged across simulations, as reflected by comparable RMSD values, the identity of residue 194 may nonetheless influence the extent of interdomain bending in LILRB1 during HLA-B binding.

We next assessed whether the presence of hydrophobic residues Ile194 or Val194 alters the non-covalent interaction landscape between LILRB1 and HLA by quantifying the occupancies of van der Waals, hydrophobic, hydrogen bond, and ionic interactions throughout the simulations. Analysis of summed occupancies across the full HLA-LILRB1 interface revealed no significant differences between valine- and isoleucine-containing HLAs for any interaction type (Figure 5A1-A4). Similarly, no differences were observed when occupancies of the overall interactions between the HLA heavy chain and LILRB1 were examined in isolation (Figure 5B1-B4). However, despite comparable overall occupancies, the extent to which individual residues contributed to this total varied depending on the presence of Val194 or Ile194 (Supplementary Tables 2, 4, 6, 8). This aligns with earlier observations of conformational differences among heavy chain residues involved in LILRB1 binding, suggesting that this polymorphism may influence α3 domain dynamism or its preferred orientation during the LILRB1-HLA interaction.

**Figure 5.**
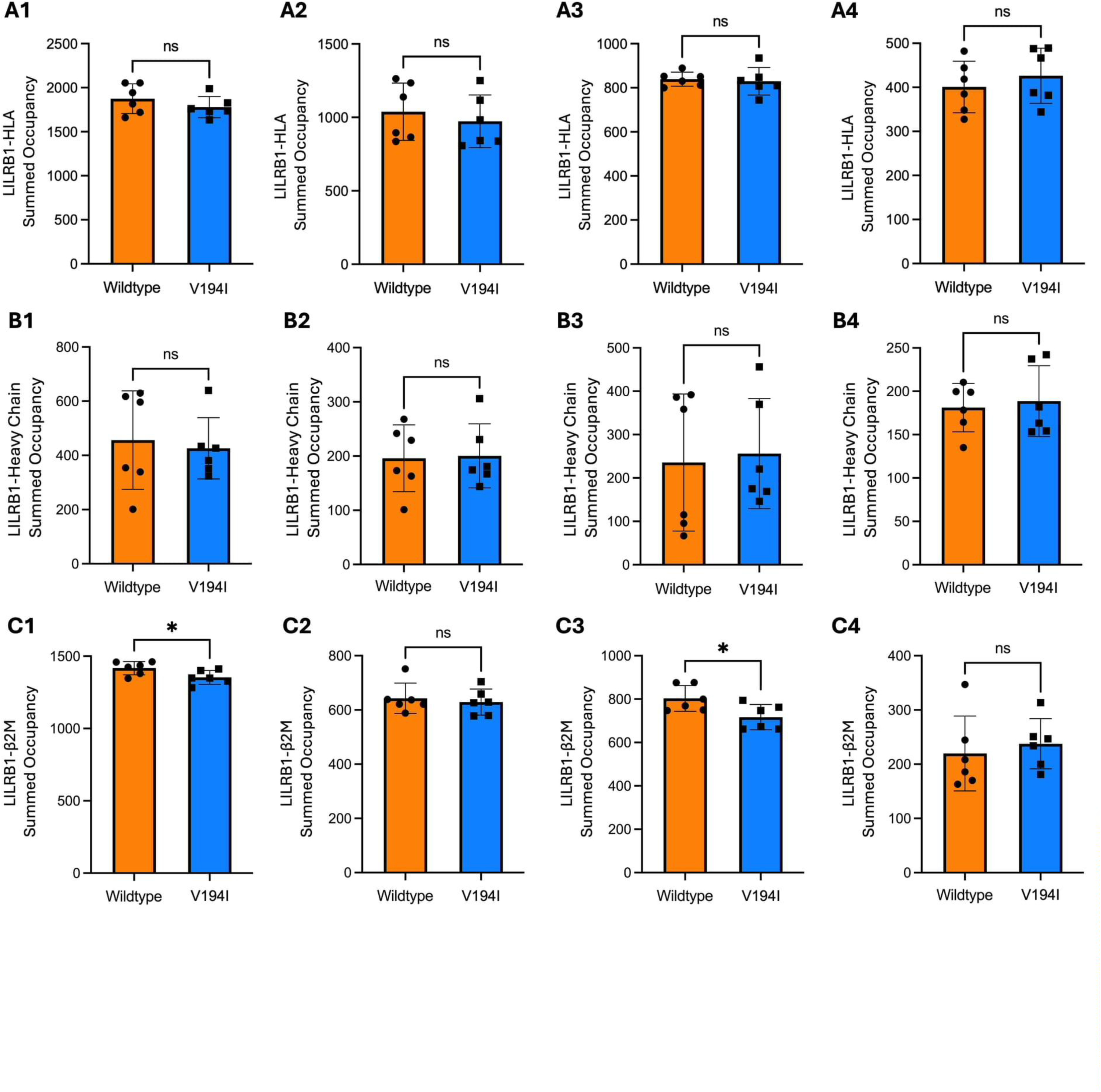
Comparison of summed occupancies of non-covalent interactions between LILRB1 and wild-type HLA-B*35:01 (orange) or HLA-B*35:01 containing a valine-to-isoleucine point mutation at residue 194 of the heavy chain (blue). Interactions between LILRB1 and the overall HLA complex **(A)**, the heavy chain **(B)**, and β2M **(C)** were quantified for (1) van der Waals, (2) hydrogen bond, (3) hydrophobic, and (4) ionic interactions. Data are presented as mean ± SEM from six independent 100-ns molecular simulations. Statistical analyses were performed using Welch’s t-test; *p<0.05.

Analysis of the interactions between the HLA-B β2M and LILRB1 revealed reduced summed occupancies for both van der Waals and hydrophobic interactions for Ile194 compared to Val194 (Figure 5C1 and 5C3). This is despite earlier broad screening panels indicating that LILRB1 binds Ile194 containing HLA-B molecules with a higher capacity than Val194. The Ile194 variant formed a greater number of transient van der Waals contacts, indicative of weaker and more spatially dispersed interactions, whereas the Val194-containing system exhibited fewer but more persistent contacts, resulting in higher overall occupancy (Supplementary Table 3). This difference may be driven by the greater intrinsic rotameric flexibility of the sec-butyl side chain of isoleucine compared to the more rigid isopropyl side chain of valine. Reduced hydrophobic interactions in the Ile194 variant were particularly evident within the β2M loop encompassing residues 91-94 (Supplementary Table 7).

Although no differences in summed hydrogen bond or ionic interaction occupancies were observed, differences in residue-level interaction preferences were observed (Supplementary Table 5, 9). These observations are consistent with the previously observed conformational differences and increased flexibility of β2M residues involved in LILRB1 binding.

Taken together, these findings suggest that the higher binding affinity of LILRB1 to HLA-B with Ile194 is unlikely to arise from an increased number of non-covalent interactions.

Instead, the data suggest that the presence of Ile194 is associated with greater conformational dynamism and increased malleability at the β2M-LILRB1 interface, which favours LILRB1 binding.

### 2.3 Mapping Functional and Mutational Sites onto the Full-Length Unbound Structure of LILRB1

To gain further insight into the structural mechanisms underlying LILRB1 function, we determined the structure of the four extracellular domains of LILRB1, D1-D4 (Figure 6A). The ectodomain of LILRB1 consists of four immunoglobulin-like domains, including domain 1 (D1; residues 35-123), domain 2 (D2; residues 124-229), domain 3 (D3; residues 230-320), and domain 4 (D4; residues 321-424). These domains are oriented at acute interdomain angles, with D1 and D2 angled at 73°, D2 and D3 at 67°, and D3 and D4 at 64°, resulting in a molecule with an overall bent and non-linear conformation (Figure 6A).

**Figure 6:**
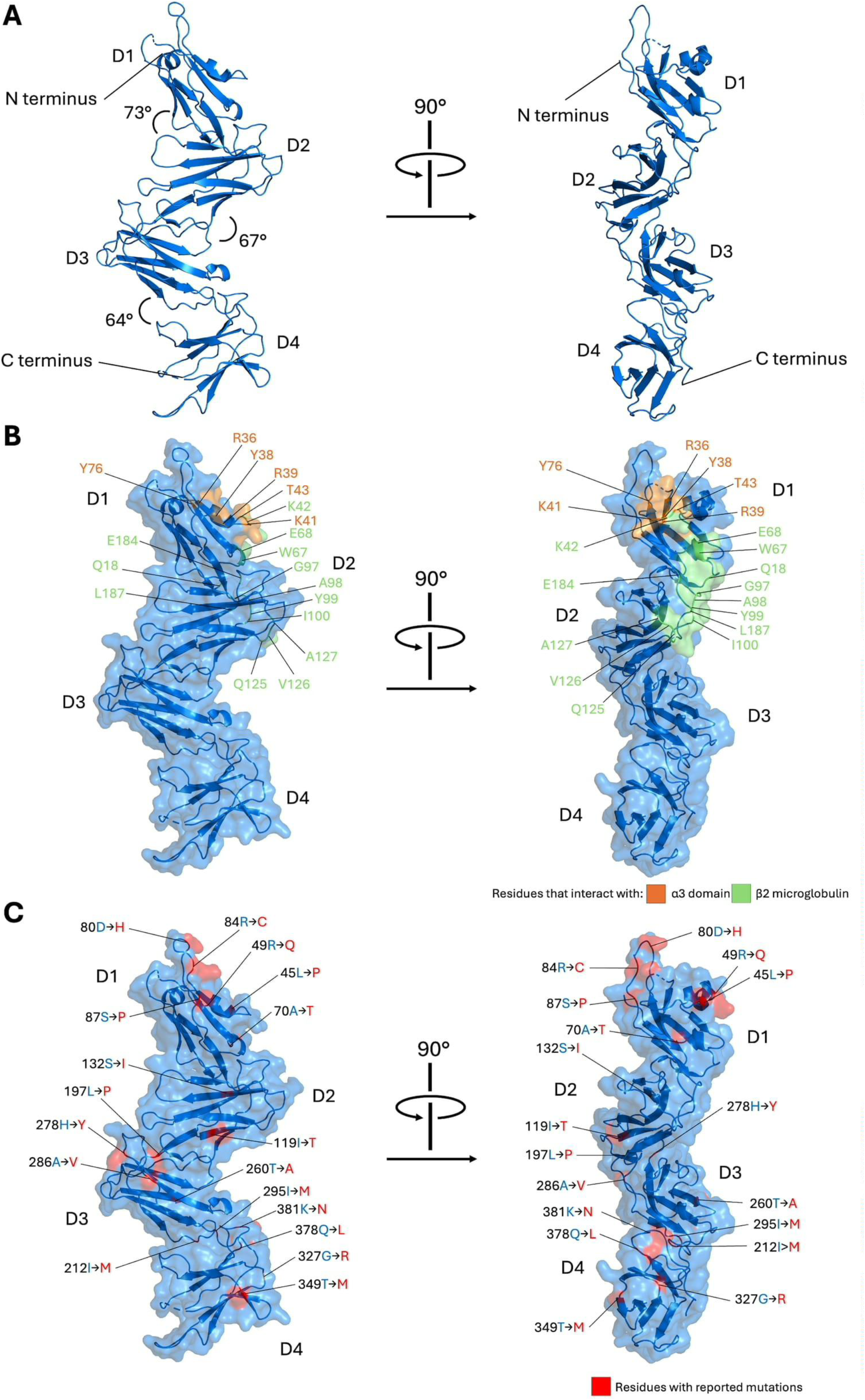
Structure of full-length unbound LILRB1. **(A)** Cartoon representations of two orthogonal views of LILRB1, with domains D1-D4 and the N- and C-termini labelled. Interdomain angles were calculated as follows: D1-D2 = 73°, D2-D3 = 67°, and D3-D4 = 64°. **(B)** Two orthogonal views of surface representations of LILRB1 with previously reported HLA-binding sites highlighted. Residues known to contact the HLA α3 domain are coloured magenta, and residues that interact with β2M are coloured orange. **(C)** Surface representations of LILRB1 from two orthogonal views with population variants, highlighted in red, mapped onto the structure.

To contextualise the interaction between LILRB1 and HLA, we mapped previously reported HLA-binding residues onto the LILRB1 structure (Figure 6B) (Willcox et al., 2003). All six residues reported to contact the HLA α3 domain, highlighted in magenta, are located within D1. In contrast, the 13 residues reported to interact with β2M, highlighted in orange, are distributed across D1 and D2, with four in D1 and nine in D2.

We also mapped naturally occurring polymorphisms of LILRB1 observed in the human population onto the structure of LILRB1 (Figure 6C) (Liu et al., 2022, Davidson et al., 2010). Of the mapped variant sites, six are within D1, three within D2, four within D3, and five within D4. Although many involve non-conservative amino acid substitutions, none overlap with known HLA-binding residues, suggesting that they are unlikely to impact HLA recognition directly.

### 2.4 Domain Dynamism in Existing LILRB1 and HLA Structures

Since we observed that domain dynamism is associated with differences in LILRB1-HLA interactions, we sought to further investigate their structural flexibilities. To do so, we compared the interdomain angles and overall conformations of our full-length unbound LILRB1 structure with previously solved LILRB1 structures in the Protein Data Bank (PDB), including partial (D1-D2 or D3-D4) and full-length forms, in both unbound and ligand-bound states. We also compiled available HLA structures from the PDB and compared the relative positioning of the α3 and β2M domains after aligning on their α1 and α2 domains.

For truncated LILRB1 D1-D2 structures, our analysis included four unbound structures (PDB: 1G0X (Chapman et al., 2000), 1UFU (Shiroishi et al., 2006), 1UGN (Brondijk et al., 2010), 1VDG (Kuroki et al., 2005)) (Figure 7A), five LILRB1-HLA-A2 complex structures (PDB: 1P7Q (Willcox et al., 2003), 4NO0 (Mohammed et al., 2017), 6EWA (Mohammed et al., 2019), 6EWC (Mohammed et al., 2019), and 6EWO (Mohammed et al., 2019)), one structure bound to HLA-F (PDB: 5KNM (Dulberger et al., 2017)), and one structure and one structure in complex with viral immune evasion molecule and HLA mimetic, UL18 (PDB: 3D2U (Yang and Bjorkman, 2008)) (Figure 7B). Among the unbound structures, D1-D2 interdomain angles ranged from 76° to 80° (Table 1). Conversely, the D1-D2 angles of the ligand-bound structures were wider, spanning 76° to 88° (Table 1). When superimposed onto our unbound full-length structure, the ligand-bound D1-D2 structures possessed lower RMSD values compared to the unbound D1-D2 structures (Table 1), indicating conformational differences between these states. These observations suggest that unbound LILRB1 adopts a more closed, compact D1-D2 conformation, which becomes more dynamic and open upon ligand binding. Such flexibility likely enables LILRB1 to accommodate residue variation across different HLA allotypes.

**Figure 7.**
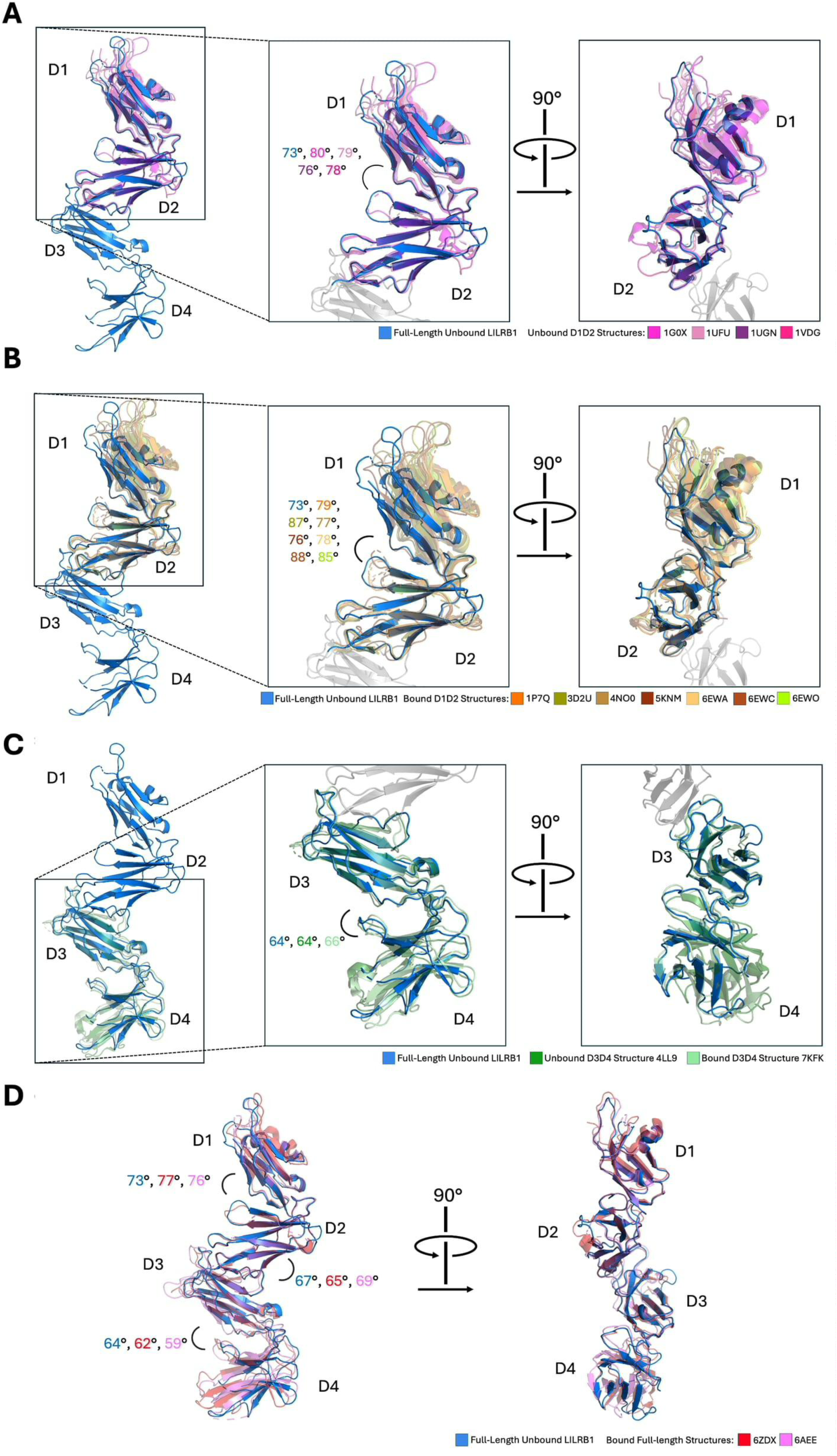
Comparison between LILRB1 structures shows domain flexibility. **(A)** Cartoon representation of full-length LILRB1 and two close-up views of the D1D2 region of full-length unbound LILRB1 overlaid with existing structures of unbound LILRB1 D1D2, aligned on D2. Accession codes of existing structures include were 1G0X (magenta), 1UFU (pink), 1UGN (purple), and 1VDG (hot pink). Interdomain D1-D2 angles calculated were 73° for this study, 80° for 1G0X, 79° for 1UFU, 76° for 1UGN, and 78° for 1VDG. **(B)** Cartoon representation of full-length LILRB1 and two close-up views of the D1D2 region of full-length unbound LILRB1 overlaid with existing structures of bound LILRB1 D1D2, aligned on D2. Accession codes of existing structures included were 1P7Q (orange), 3D2U (olive), 4NO0 (light brown), 5KNM (chocolate), 6EWA (pale yellow), 6EWC (pale brown), and 6EWO (light green). Interdomain D1-D2 angles calculated were 73° for this study, 87° for 3D2U, 77° for 4NO0, 76° for 5KNM, 78° for 6EWA, 88° for 6EWC, and 85° for 6EWO. **(C)** Cartoon representation of full-length LILRB1 and two close-up views of the D3D4 region overlaid with an unbound D3D4 structure, aligned on D3. Existing structures include an unbound D3D4 structure with accession code 4LL9 (green) and a bound D3D4 structure with accession code 7KFK (light teal). Interdomain D3-D4 angles calculated were 64° for this study, 64° for 4LL9, and 66° for 7KFK. **(D)** Cartoon representation of two orthogonal views of full-length unbound LILRB1 overlaid with full-length bound structures with accession codes 6ZDX (red) and 6AEE (pink), aligned on D2. Interdomain angles for D1-D2, D2-D3, and D3-D4 were 73°, 67°, and 64° for our structure, 73°, 65°, and 62° for 6ZDX, and 76°, 69°, 59° for 6AEE.

We also compared our D1-D4 unbound LILRB1 structure to D3-D4-only structures. Alignment on D3 revealed structural similarity with both an unbound D3-D4 structure (PDB: 4LL9 (Nam et al., 2013)) and a D3-D4 structure bound by a malarial Repetitive Interspersed Family (RIFIN) protein (PDB: 7KFK (Chen et al., 2021)) (Figure 7C). The interdomain angle between D3 and D4 was highly consistent across structures, ranging from 64° to 66°, with RMSD values further supporting their structural similarity (Table 1). These findings suggest that the D3-D4 region of LILRB1 remains relatively consistent and may function primarily as a structural scaffold.

Furthermore, we compared full-length bound structures aligned on D2, including LILRB1 bound by RIFIN (PDB: 6ZDX (Harrison et al., 2020)) and in complex with HLA-G (PDB: 6AEE (Wang et al., 2020)) (Figure 7D). These structures aligned closely with our unbound structure, with RMSD values of 1.5 Å and 1.2 Å, respectively (Table 1). This similarity is further reflected in the narrow range of interdomain angles, with D1-D2 ranging from 73° to 77°, D2-D3 ranging from 65° to 69°, and D3-D4 ranging from 59° to 64° (Table 1). Notably, the angular differences observed between the full-length bound and unbound LILRB1 structures are less pronounced than those observed when comparing only the D1-D2 LILRB1 structures (Figures 7A and 7B), while the D3-D4 interdomain angles remain comparably consistent across both full-length and truncated structures (Figure 7C). However, given the limited number of available full-length LILRB1 structures, and the absence of structures of full-length LILRB1 in complex with classical HLA-I, these comparisons are unlikely to capture the full extent of domain dynamism in D1-D4 LILRB1.

Finally, we compiled available structures of HLA-A, HLA-B, and HLA-C and aligned them on the α1 and α2 domains to visualise movements in the α3 and β2m domains. For clarity, only the extreme conformations, defined by the greatest centre-of-mass separation between α3 or β2m domains, are shown. Between the α3 extremes (PDB: 1EFX (Boyington et al., 2000) and 7UC5 (Nguyen et al., 2022)), there was an angular difference of 18°, corresponding to a centre-of-mass separation of 8.16 Å (Figure 8A). When LILRB1-bound classical HLA structures, which included five HLA-A2 structures, were overlaid, their α3 domains were distributed across this range (Figure 8A). Similarly, analysis of β2m extremes (PDB: 3MV7 (Gras et al., 2010) and 7STF (Wright et al., 2023)) revealed a maximum centre-of-mass separation of 4.52 Å (Figure 8B). Imposing the LILRB1-bound HLA-A2 structures showed that β2m positions likewise span this conformational range (Figure 8B). Together, these observations indicate substantial dynamism in both the α3 and β2m domains of HLA. This flexibility, in conjunction with the dynamism of the LILRB1 D1-D2 domain, suggests potential complementary roles for domain dynamics during LILRB1-HLA interactions.

**Figure 8.**
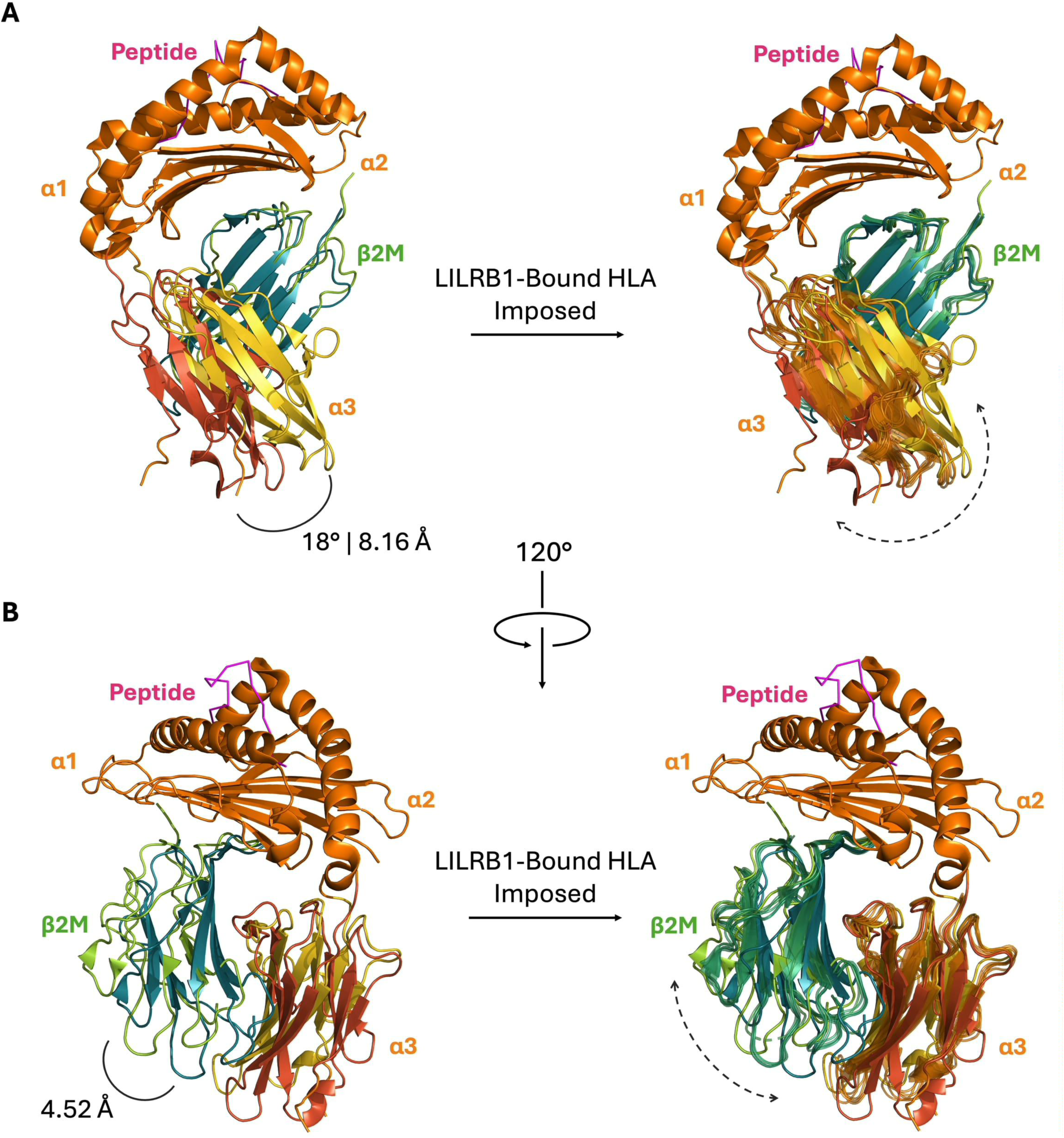
Domain dynamism in the α3 and β2M domains of human leukocyte antigens (HLA). **(A)** Structural representation of HLA illustrating the movement range of the α3 domain. Extreme conformations were derived from previously reported classical HLA structures, aligned on the α1 and α2 domains (1EFX in red and 7UC5 in yellow), based on distances between the centres of mass of the α3 domains and their inter-domain orientations. **(B)** Structural representation of HLA rotated 120° anticlockwise to highlight β2M dynamics. β2M extremes (3MV7 in dark green and 7STF in light green) were derived with a similar approach. These extremes are overlaid with structures of HLA in complex with previously reported LILRB1-HLA complex structures (1P7Q, 4NO0, 6EWA, 6EWC, and 6EWO).

### 2.5 Mapping LILRB1 Antibody Binding Sites onto the LILRB1 Structure

LILRB1 has emerged as a promising target for checkpoint immunotherapeutic intervention, as evidenced by a growing number of monoclonal antibodies currently spanning various preclinical and clinical developmental stages (Hu et al., 2024). To gain insight into their potential modes of action, and to assess whether LILRB1 dynamism is relevant to its therapeutic targeting, we modelled LILRB1 binding to a selection of therapeutic antibodies using AlphaFold 3 and mapped these predicted interactions onto the LILRB1 structure for visualisation (Figure 9). A total of ten antibodies (Sequences in Supplementary Figure 8), selected based on predicted pTM and iPTM scores exceeding 0.6 during modelling, were included in this analysis. Antibody 6 was derived from patent WO2021222544A1 (Duey, 2021), while antibodies 1 to 5 and 7 to 10 were extrapolated from patent WO2022026360A2 (An, 2021). The interactions between these antibodies and LILRB1 were modelled using AlphaFold 3 and categorised according to their reported function, with antibodies 1 to 6 classified as antagonists and antibodies 7 to 10 as agonists (Full list of contacts in Supplementary Tables 10-19).

**Figure 9:**
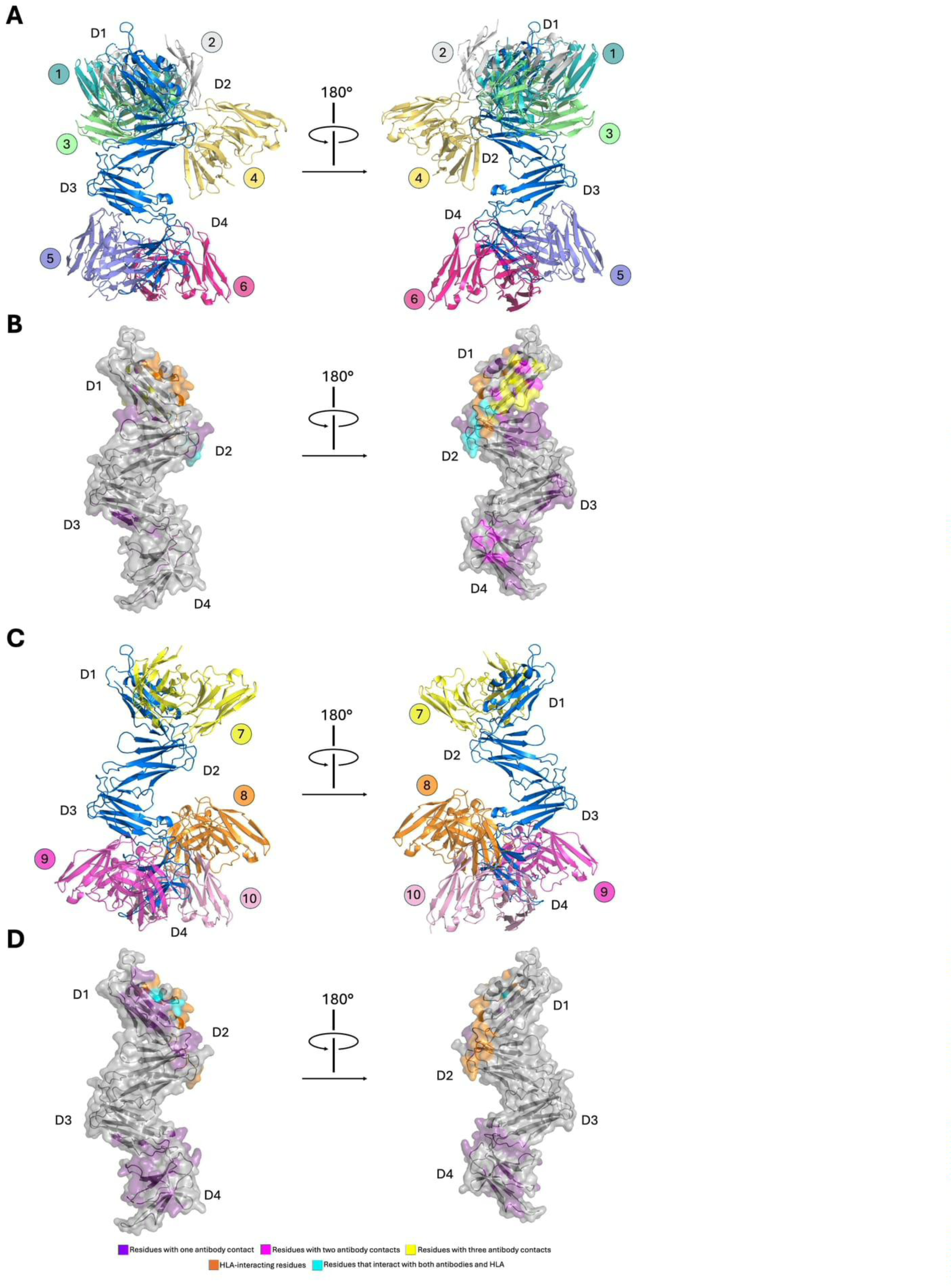
Predicted interactions between LILRB1 and existing LILRB1-binding antibodies. Structural models were generated using AlphaFold 3 to visualize the interactions between LILRB1 and 12 published monoclonal antibodies. **(A)** Cartoon representations of two orthogonal views showing the variable heavy and light chain domains of six antagonistic LILRB1-binding antibodies in complex with full-length unbound LILRB1. Antibodies are labelled 1-6 and coloured as follows: antibody 1 in turquoise, antibody 2 in grey, antibody 3 in green, antibody 4 in yellow, antibody 5 in purple, and antibody 6 in magenta. **(B)** Two orthogonal surface views of LILRB1 showing antibody contact regions. Contacts were defined as residues within 4.0 Å of any antibody heavy or light chain. Interaction sites on LILRB1 are coloured according to frequency: purple for one antibody contact, magenta for two antibody contacts, yellow for three antibody contacts, and brown for four antibody contacts. Regions known to interact with HLA are shown in pink while residues that interact with both HLA and at least one antibody are shown in orange. **(C)** Cartoon representations of two orthogonal views showing the variable heavy and light chain domains of four agonistic LILRB1-binding antibodies in complex with full-length unbound LILRB1. Antibodies are labelled 7-12 and coloured as follows: antibody 7 in yellow, antibody 8 in orange, antibody 9 in magenta, antibody 10 in pink. **(D)** Two orthogonal surface views of LILRB1 showing mapped antibody contact regions on D3D4, with contacts defined and colour coded as in **(B)**.

Among the six modelled antagonistic antibodies, four interact with regions in D1-D2 (Ab 1-4), while two interact with regions in D3-D4 (Ab 5 and 6; Figure 9A and 9B). Of the four D1-D2-interacting antibodies, their epitopes localised around the interdomain hinge region, particularly on the orthogonal face where much of the surface of β2M-interacting residues of LILRB1 is exposed. This trend highlights the importance of this structural plane of LILRB1, suggesting that blocking of this region likely inhibits LILRB1-HLA interactions through steric hinderance. However, the frequent targeting of sites outside the HLA-binding interface suggests that additional mechanisms may contribute to inhibition, such as compromising the D1-D2 conformational dynamics required for HLA binding. The presence of two antagonistic antibodies targeting the D3-D4 region indicates that LILRB1 inhibition can occur without directly engaging the HLA-binding interface. These effects may be mediated through steric hindrance, disruption of receptor clustering, or allosteric effects that alter D1-D2 conformational dynamics and impair HLA engagement.

Among the four modelled agonistic antibodies, one interacts with D1-D2 (Ab 7), while three interact with D3-D4 (Ab 8-10; Figure 9C and 9D). The D1-D2-interacting antibody (Ab 7) binds to residues located near the HLA-binding interface of LILRB1, including several residues directly involved in HLA recognition. Additionally, D3-D4-interacting agonistic antibodies (Ab 8-10) indicates that binding this region may also promote LILRB1 signalling. Mechanisms for agonism include receptor clustering or allosteric effects that stabilise the receptor.

## 3 Discussion

LILRB1 is a critical component of effective immune surveillance and tolerance and displays one of the broadest HLA binding profiles of any immune receptor. In this study, we aimed to elucidate the structural mechanisms underlying broad LILRB1-HLA binding and to define the drivers of subtle affinity differences that contribute to HLA allotype-specific preferences. Overall, LILRB1 bound all tested HLA allotypes, likely enabled in part by conformational adaptability and flexbility, as reflected in the D1-D2 interdomain angle variability and RMSD differences between ligand-bound and unbound states. However, binding differences between allotypes was observed, implying that this adaptability does not allow for fully uniform HLA affinity. Results from our broad screening panel and molecular dynamics analyses imply that HLA stability and domain dynamism may contribute to these differences. Finally, the epitopes of modelled antibody-LILRB1 interactions localised to the D1-D2 and D3-D4 hinge regions, both within and beyond the HLA-binding interface, indicating the functional importance of interdomain flexibility and the therapeutic potential of these regions.

Comparisons between our single-antigen bead binding results and previous studies on LILRB1-HLA interactions reveal notable differences. While previous SPR studies on LILRB1-HLA interactions, which included HLA-A*11, HLA-B*27, HLA-B*35, and several HLA-C allotypes, reported no binding differences (Chapman et al., 1999, Shiroishi et al., 2003, Kuroki et al., 2005), our broad screening panel indicates that LILRB1 can discriminate between these HLA allotypes. These discrepancies may stem from differences in peptide repertoires, as the cell-derived HLA molecules on the single-antigen beads possess heterogeneous repertoires (Ellis, 2013), whereas the HLA in SPR measurements are typically refolded with a single high-affinity peptide, resulting in more stable molecules (Chapman et al., 1999, Shiroishi et al., 2003, Kuroki et al., 2005). Consequently, SPR measurements may overlook nuances, such as stability differences, observed with a cellular repertoire.

Differences are also evident among studies using HLA single-antigen beads. Jones et al. (Jones et al., 2011) reported that LILRB1 discriminates only among HLA-A allotypes, with no differences across HLA-B or HLA-C allotypes. Conversely, Liu et al. (Liu et al., 2022) previously observed similar discrimination among HLA-B allotypes and the weaker binding to HLA-C allotypes, but also reported differences in HLA-A allotype binding different from those described by Jones et al. (Jones et al., 2011). These inconsistencies likely arise from methodological differences, where Jones et al. (Jones et al., 2011) used Fc constructs of D1-D4 LILRB1, Liu et al. (Liu et al., 2022) used Fc constructs of D1-D2 LILRB1, while we used D1-D4 LILRB1 tetramers. Together, these discrepancies highlight the complexity of LILRB1-HLA interactions, indicating that LILRB1 is not simply a pan-HLA binder but exhibits context-dependent binding behaviour.

Our molecular dynamics analysis of polymorphisms associated with differences in LILRB1 binding indicates that domain dynamism may influence LILRB1-HLA interactions. These analyses focused on interactions between LILRB1 and HLA-B*35 containing the WT Val194 or Ile194 point-mutant, the only contact residue polymorphism that affected LILRB1 binding in our broad screen. Despite its highly conservative nature, this substitution induced long-range allosteric effects on β2M, where Val194 was associated with a less dynamic β2M that formed more persistent interactions with LILRB1, while Ile194 introduced to a more dynamic β2M that formed less consistent interactions with LILRB1. This observation appears to contradict the findings from the broad screen, wherein LILRB1 exhibited weaker binding to HLA-B allotypes with Val194 compared to those with Ile194. However, prior structural and kinetic evidence has indicated that the D1-D2 flexibility of LILRB1 constitutes a core feature of its interactions with HLA molecules (Shiroishi et al., 2006). Moreover, our comparison of existing LILRB1 and HLA structures indicated that the range of movement of the D1-D2 region of LILRB1 is larger when associated with ligand binding, while the α3 and β2M domains of HLA possesses substantial intrinsic flexibility that persists during LILRB1 complexation. Therefore, in the context of LILRB1-HLA interactions, domain dynamism may actively facilitate robust complex formation, suggesting a degree of induced fit is required.

Beyond HLA α3 residue polymorphisms and their potential effects on domain dynamism, another factor likely influencing LILRB1-HLA interactions in our screening panel is the stability of the HLA molecules themselves. Since the HLA molecules on single-antigen beads are cell-derived and present heterogeneous peptide repertoires, there is often bead-to-bead variation between HLA allotypes, correlating with differences in the extent of protein denaturation, misfolding, and conformational heterogeneity (Ellis, 2013, Dédier et al., 2000, Jappe et al., 2020). The influence of these factors is illustrated by the similar relative binding patterns across HLA allotypes observed for LILRB1 and the pan-HLA monoclonal antibody W6/32, despite their non-overlapping contact regions on HLA (Pymm et al., 2024, Willcox et al., 2003). In addition to bead-specific effects, intrinsic peptide-dependent stability differences between different HLA allotypes may have also affected LILRB1 binding. This notion is supported by prior work reporting a descending order of thermostability for cell-derived HLA-B*07:02, HLA-A*02:01, and HLA-C*04:01 (Jappe et al., 2020), which parallels the relative binding of LILRB1 and W6/32 to these allotypes observed in our bead assay. However, binding mode and affinity differences between LILRB1 and W6/32 likely confer distinct sensitivities to HLA stability (Pymm et al., 2024). Indeed, the binding discrepancy between the strongest and weakest allotypes is substantially larger for LILRB1 than for W6/32. Such sensitivity differences may reflect avidity differences, with W6/32 being dimeric and out LILRB1 being tetrameric. Furthermore, sensitivity differences may explain the weaker binding of LILRB1 to HLA-C allotypes, even after normalisation to W6/32 binding, as HLA-C molecules exhibit lower cell surface expression and stability than HLA-A and HLA-B (McCutcheon et al., 1995, Zemmour and Parham, 1992, Jappe et al., 2020). Overall, alongside domain dynamism differences associated with α3 residue polymorphisms, intrinsic stability differences between HLA allotypes in complex with their cell-derived peptide repertoires may also influence LILRB1 binding.

The epitopes in our modelled antibody-LILRB1 interactions cluster around the D1-D2 and D3-D4 hinges, highlighting the functional importance of these regions. Given that the D1-D2 region mediates HLA binding (Willcox et al., 2003), it is unsurprising that it is targeted by both agonistic and antagonistic antibodies. Agonism at this site may occur via HLA mimicry, as exemplified by Plasmodium RIFINs, which exploit this strategy for immune evasion (Harrison et al., 2020). Conversely, antagonism may occur through preventing HLA binding, and at least one previously reported antagonistic antibody has been mapped to the D1-D2 region (Chen et al., 2020). Several modelled agonistic and antagonistic antibodies also bound to the D3-D4 region, despite its lack of involvement in HLA binding (Wang et al., 2020).

Mechanistically, these antibodies may modulate receptor dynamism or receptor clustering. The plausibility of agonism via the D3-D4 region is supported by the observation that RIFINs bound to D3 to induce LILRB1 signalling (Chen et al., 2021). Similarly, antagonism is supported by the identification of an antibody that inhibits LILRB1 signalling by targeting around the D3-D4 region (Wicher et al., 2023). Notably, the lack of modelled antibodies targeting the D2-D3 hinge aligns with the current understanding that this region functions primarily as a structural scaffold (Wang et al., 2020). Together, these findings identify the D1-D2 and D3-D4 regions as key functional sites of LILRB1 and highlight their potential as therapeutic targets for modulating receptor activity, either by directly blocking HLA binding or indirectly by altering receptor clustering or conformational dynamics.

Ultimately, this study demonstrates that LILRB1 acts as a broad pan-HLA receptor, although the strength and nature of LILRB1-HLA interactions are modulated by α3 domain dynamics driven by residue polymorphisms and by intrinsic stability differences between HLA allotypes. This hierarchy of binding may have important implications for interpreting and stratifying HLA-associated disease susceptibility. Furthermore, these findings suggest that therapeutic targeting of the D1-2 versus D3-D4 regions of LILRB1 may produce distinct functional outcomes depending on the HLA allotype involved.

## 4 Materials and Methods

### 4.1 Protein Expression and Purification

The LILRB1 ectodomain (residues 24-423) was cloned into the pHLsec mammalian expression vector (AgeI/KpnI) with a C-terminal 6xHis purification tag (AddGene plasmid #99845). The plasmid for secreted BirA Ligase with a C-terminal Flag purification tag was purchased (AddGene plasmid #64395). LILRB1 and BirA Ligase were co-expressed in Expi293F GnTI-cells. Secreted LILRB1 was harvested and purified from culture media seven days after transfection. For purification, the media was diluted four-fold into 10 mM imidazole, 10 mM Tris (pH 8.0), and 300 mM NaCl, before being passed through a 5 mL HisTrap FF column (Cytiva). Bound LILRB1 was eluted with 10 mM Tris pH 8.0, 300 mM NaCl, and 50 mM EDTA. The protein underwent further purification via size-exclusion chromatography using a HiLoad 16/600 Superdex 200 pg column. Protein identity and quality were analysed by SDS-PAGE, and biotinylation was verified using a streptavidin pulldown assay.

### 4.2 HLA Binding Assay

PE-conjugated LILRB1 tetramers were made by incubating biotinylated LILRB1 with eBioscience Streptavidin-PE Conjugate. The binding assay utilized a panel of single antigen beads (One Lambda Thermo Fisher Scientific, Cat. #LS1A04, lot 015), each coated with one of 97 classical HLA molecules, including 31 HLA-A, 50 HLA-B, and 16 HLA-C alleles. PE-conjugated LILRB1 tetramers (5 μg per test) were incubated with HLA-coated beads for 30 minutes at room temperature in the dark in PBS containing 5 mM EDTA supplemented with 5% fetal calf serum. Beads were washed three times in LABScreen wash buffer, and fluorescence intensity was measured using the Luminex platform (FLEXMAP 3D; One Lambda). Samples were run in triplicate and the binding of pan-HLA monoclonal antibody W6/32 was used as a positive control (supplier provided for lot 015). LILRB1 mean fluorescence intensity (MFI) values were normalised to W6/32 using the following formula:

*(Maximum W6/32 MFI ÷ W6/32 MFI for the allele) × LILRB1 MFI for the allele*.

### 4.3 Crystallisation and Data Collection

LILRB1 was concentrated to 5.0 mg/mL in 10 mM Tris pH 8.0 and 300 mM NaCl, deglycosylated with endoglycosidase H (New England Biolabs), and crystallized using the hanging-drop vapor diffusion method at 298 K. The crystallization condition included a mother liquor of 0.8 M NaH_2_PO_4_/K_2_HPO_4_ (pH 5.2) with 0.02% sodium azide. Protein drops were prepared by mixing protein solution and mother liquor in a 1:1 volume ratio equilibrated over 1000 μL reservoir. For freezing, crystals were cryoprotected using the mother liquor supplemented with 10% glycerol. X-ray diffraction data were collected at 100 K on the MX2 beamline at the Australian Synchrotron.

### 4.4 Structural Determination, Refinement, and Analysis

LILRB1 crystals diffracted to a resolution of 3.0 Å and belonged to the space group *P*2_1_, with unit cell dimensions of a = 60.03 Å, b = 35.82 Å, c = 105.85 Å, and β = 98.30°. Diffraction data were processed using XDS and SCALA (Kabsch, 2010, Evans, 2006). Model building and refinement were performed iteratively using Coot and PHENIX (Emsley and Cowtan, 2004, Adams et al., 2010). Final model validation was conducted with MolProbity (Davis et al., 2007). Refinement statistics are summarised in (Supplementary Table 1). Coordinates and structure factors have been deposited in the Protein Data Bank under accession code (27IW). Structural analyses, including identification of contacting residues, structural alignment, interdomain angle measurements, and calculation of root-mean-square deviation (RMSD) values, were performed using PyMOL (DeLano, 2002).

### 4.5 Molecular Dynamics Simulations

Molecular dynamics (MD) simulations were performed using GROMACS 2025.3 (Abraham et al., 2015). Initial simulation models were generated by merging existing structures of LILRB1 in complex with HLA-A*02 (PDB ID: 6EWO) and HLA-B*35 (PDB ID: 1XH3) (Mohammed et al., 2019, Probst-Kepper et al., 2004). Missing residues were manually built and chemically validated using Coot and PyMOL and point mutations at residue 194 were generated using PyMOL (Emsley and Cowtan, 2004, DeLano, 2002). Simulation parameters and protein solvation were performed using CHARMM-GUI (Jo et al., 2008). Systems were solvated with model-TIP3P waters in rectangular water boxes with an edge distance of 10 Å and 0.15 M concentration of NaCl. Grid information for PME FFT was automatically generated. The force field CHARMM36m was used throughout (Huang et al., 2017). Simulations were performed at a temperature of 303.15 K and pressure at 1 bar. The systems were minimised and equilibrated for 125 ps sequentially under NVT and NPT ensemble. Production MD simulations were run for 100 ns in six replicates for each condition. Trajectory analyses were performed using GROMACS after correction for periodic boundary conditions. Root mean square deviation (RMSD) was calculated for the domains of each protein binding partner and for residues reported to participate in the LILRB1-HLA interface. An overall RMSD value for each component in each independent simulation was obtained by averaging RMSD values over the 100 ns trajectory. Root mean square fluctuation (RMSF) was computed for residues in both proteins. Noncovalent interactions between LILRB1 and HLA were analysed using ProLIF (Bouysset and Fiorucci, 2021). Interaction occupancies for which the average across three independent simulations was below 10% in both the wild-type HLA-B35 and the HLA-B35 variant containing a valine-to-isoleucine substitution at residue 194 of the heavy chain were excluded from further analysis.

### 4.6 Modelling Antibody Interactions

Complementarity-determining regions (CDRs) and variable heavy and light chain sequences of anti-LILRB1 antibodies, along with their reported functions, were obtained from published patents WO2021222544A1 (Duey, 2021) and WO2022026360A2 (An, 2021). Antibody-LILRB1 interactions were modelled using AlphaFold3 with the sequence of our LILRB1 construct (residues 24-423) (Jumper et al., 2021). Only models with both a predicted template modelling score (pTM) and interface predicted template modelling score (ipTM) greater than 0.6 were collated for antibody mapping. This process was accompanied by manual inspection to assess plausibility. Residues on LILRB1 and the antibodies were considered contact residues if they were within 4 Å.

### 4.7 Statistical Analysis

Data were analysed using GraphPad Prism 10 (Swift, 1997). Statistical analyses were conducted using the Kruskal-Wallis or Mann-Whitney tests, or Welch’s t-test, with p ≤ 0.05 considered statistically significant.

## Supplementary Material

Supplementary Figures S1-S4 present raw, unnormalised mean fluorescence intensity (MFI) values for LILRB1 tetramers and the pan-HLA monoclonal antibody W6/32 measured using Luminex. Supplementary Figures S2-S4 additionally compare sequence variation within the α3 domain of the analysed HLA allotypes. Supplementary Figure S5 shows stereo views of the point mutation interrogated in the molecular dynamics simulations. Supplementary Figures S6 and S7 present root-mean-square deviation (RMSD) metrics from the molecular dynamics simulation runs. Supplementary Figure S8 displays sequence alignments of the antibodies used for modelling.

Table S1 provides the data collection and refinement statistics for the LILRB1 crystal structure. Supplementary Tables S2-S9 summarise occupancy of the residues forming non-covalent interactions during the molecular dynamics simulations. Supplementary Tables S10-S19 list the contacts formed between LILRB1 and the modelled antibodies.

## Data availability

All data are contained within the manuscript. All data are available upon request

## Supporting information

This article contains supporting information.

## Acknowledgements

This project was supported by St Vincent’s Institute of Medical Research (Australia) and in part by the Victorian Government’s Operational Infrastructure Support Program. This work was conducted, in part, at the MX2 beamline of the Australian Synchrotron, part of ANSTO.

## Author contributions

GXYZ, JQT., LS, JR, AJO, MNKK, MC, BRT, JPV and CGL provided reagents, performed the experiments and analysis. ADB and JKH provided conceptual input. CGL and JPV conceived the study. GXYZ, JPV and CGL wrote the initial draft and all authors contributed to the final manuscript.

## Supplementary Figures

**Figure S1.**
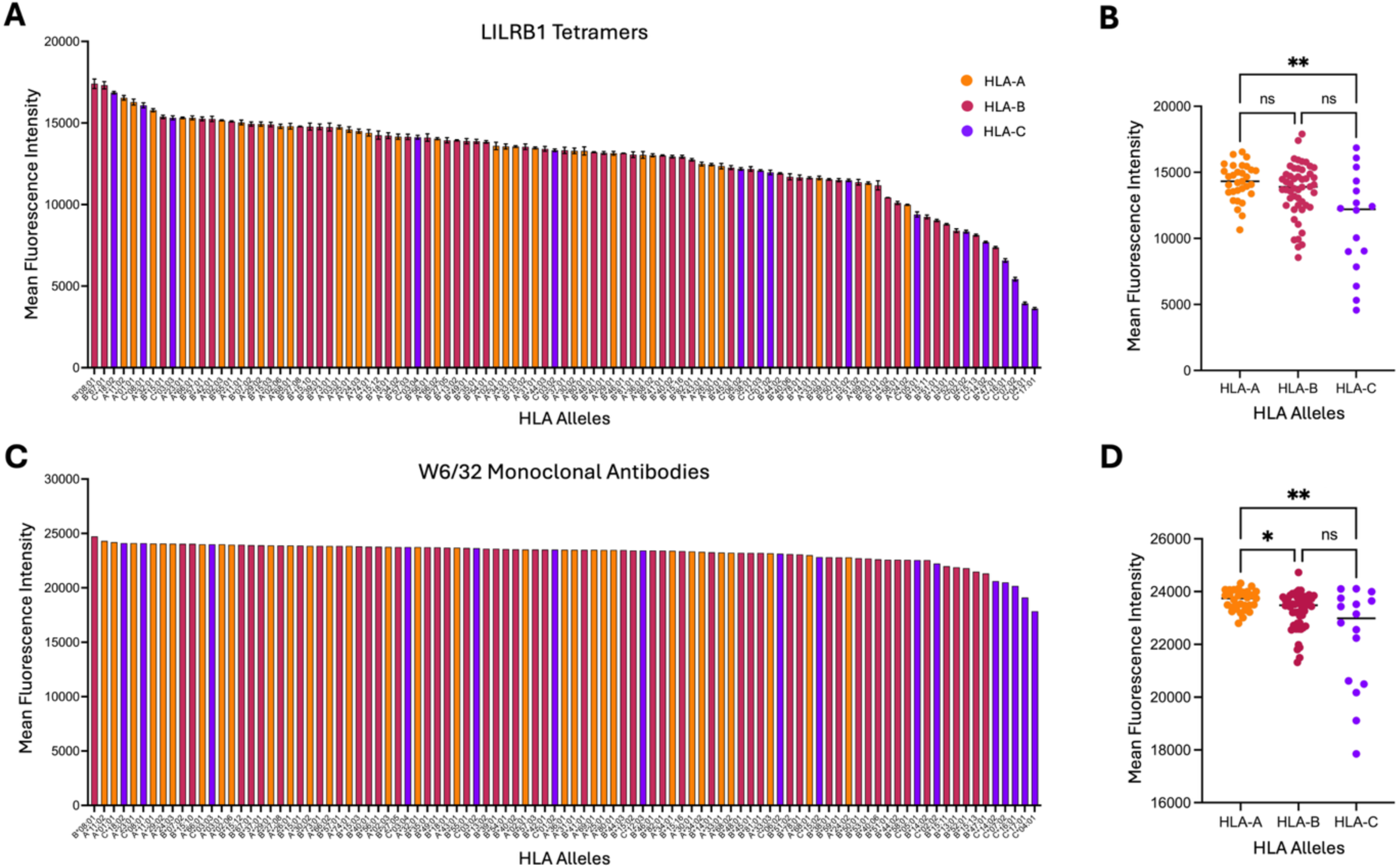
Interactions between LILRB1 tetramers or W6/32 and HLA molecules, shown as raw mean fluorescence intensity (MFI) values. (A) Binding of LILRB1 tetramers to 97 HLA allotypes on single-antigen beads, comprising 31 HLA-A (orange), 50 HLA-B (magenta), and 16 HLA-C (purple) allotypes. (B) Comparison of LILRB1 binding across HLA-A, HLA-B, and HLA-C allotypes. (C) Binding of W6/32 to the same 97 HLA allotypes. (D) Comparison of W6/32 binding across HLA-A, HLA-B, and HLA-C allotypes. Bars for LILRB1 represent mean values from three independent replicates ± SEM, whereas W6/32 data are from a single replicate. Statistical analyses were performed using the Mann–Whitney U test or Kruskal-Wallis test; *p < 0.05.

**Figure S2:**
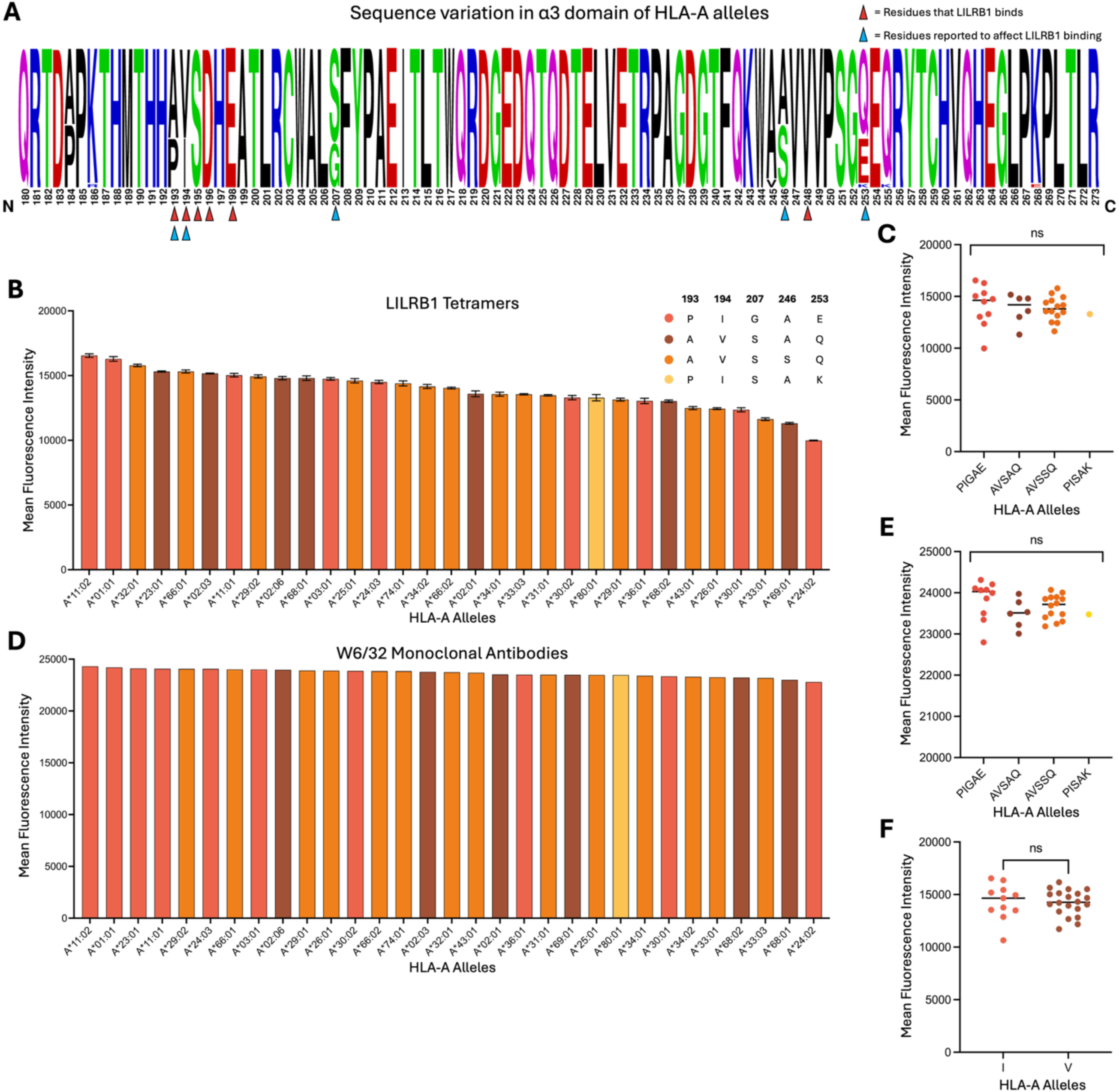
Interactions between LILRB1 tetramers or W6/32 and HLA-A allotypes, shown as raw mean fluorescence intensity (MFI) values. **(A)** Sequence variation within the α3 domain across the included HLA-A allotypes, with residues reported to form the LILRB1 interaction interface and residues previously correlated with differences in LILRB1 binding marked with red and blue arrows, respectively. **(B)** Binding of LILRB1 tetramers to 31 HLA-A allotypes, with bars coloured according to specific sequence motifs at positions 193, 194, 207, 246, and 253: vermillion for PIGAE, orange for AVSAQ, brown for AVSSQ, and yellow for PISAK. **(C)** Comparison of LILRB1 binding across HLA-A allotypes grouped by motif category. **(D)** Binding of W6/32 to the same 31 HLA-A allotypes grouped by motif category. **(E)** Comparison of W6/32 binding across HLA-A allotype groups. Statistical analyses were performed using the Mann-Whitney U test; *p < 0.05.

**Figure S3:**
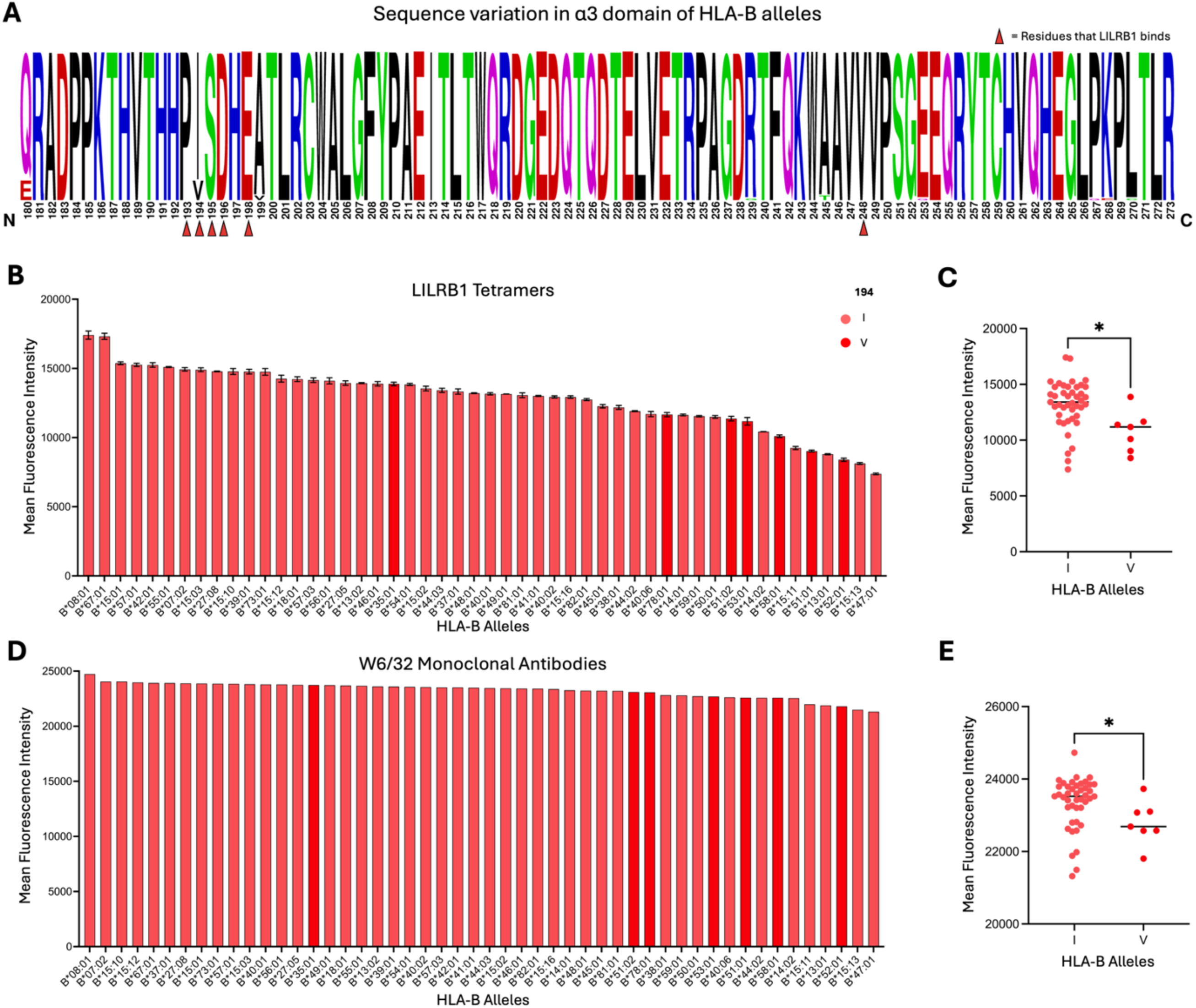
Interactions between LILRB1 tetramers or W6/32 and HLA-B allotypes, shown as raw mean fluorescence intensity (MFI) values. **(A)** Sequence variation within the α3 domain across the included HLA-B allotypes, with residues reported to form the LILRB1 interaction interface marked with red arrows. **(B)** Binding of LILRB1 tetramers to 50 HLA-B allotypes, with bars coloured by the residue at position 194: pink for I and red for V. **(C)** Comparison of LILRB1 binding across HLA-B allotypes grouped by motif category. **(D)** Binding of W6/32 to the same 50 HLA-B allotypes grouped by motif category. **(E)** Comparison of W6/32 binding across HLA-B allotype groups. Statistical analyses were performed using the Kruskal-Wallis test; *p < 0.05.

**Figure S4:**
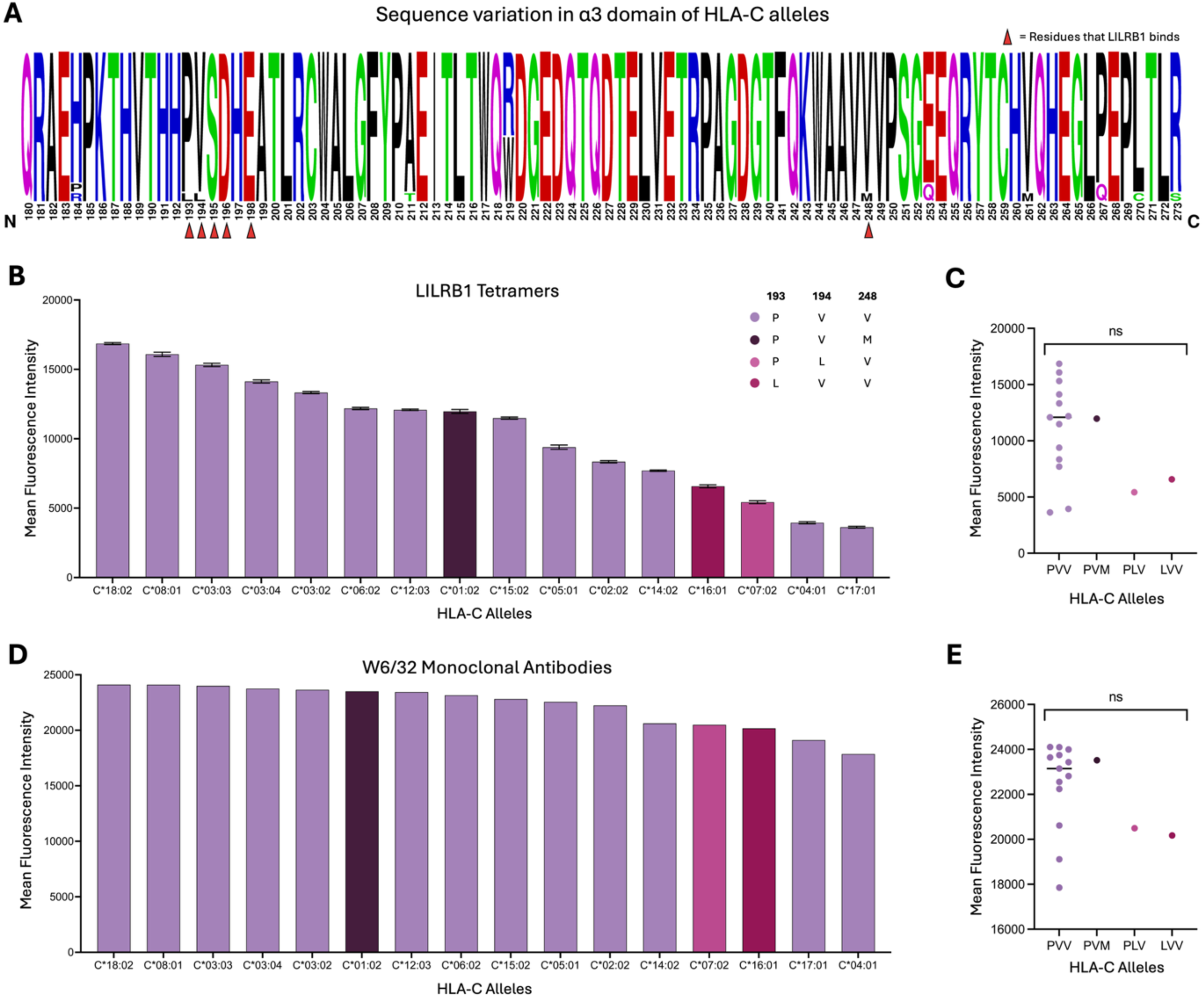
Interactions between LILRB1 tetramers or W6/32 and HLA-C allotypes, shown as raw mean fluorescence intensity (MFI) values. **(A)** Sequence variation within the α3 domain across the included HLA-C allotypes, with residues reported to form the LILRB1 interaction interface marked with red arrows**. (B)** Binding of LILRB1 tetramers to 16 common HLA-C allotypes, with bars coloured according to residues at positions 193, 194, and 248: lilac for PVV, dark purple for PVM, red violet for PLV, and light magenta for LVV. **(C)** Comparison of LILRB1 binding across HLA-C allotypes grouped by motif category. **(D)** Binding of W6/32 to the same 16 HLA-C allotypes grouped by motif category. **(E)** Comparison of W6/32 binding across HLA-C allotype groups. Statistical analyses were performed using the Mann-Whitney U test; *p < 0.05.

**Figure S5.**
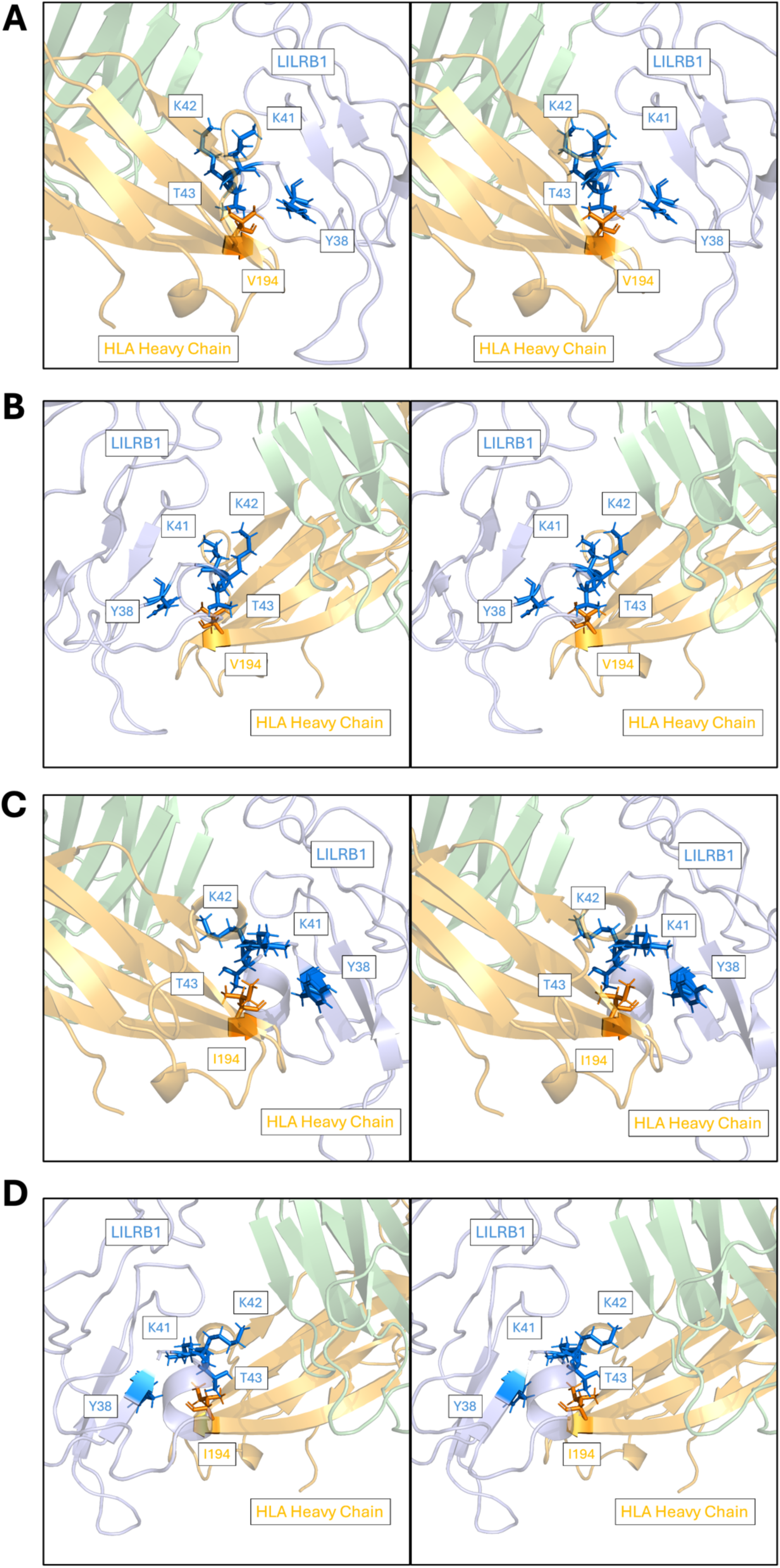
Stereo views of equilibrated structures of LILRB1 in complex with HLA-B35 following 5 ns of molecular dynamics simulation. Interactions between LILRB1 and HLA-B35 are shown for **(A)** Val194 and **(B)** its mirrored counterpart, and **(C)** Ile194 and **(D)** its mirrored counterpart. In all panels, proteins are rendered in cartoon representation, with residue 194 of the HLA heavy chain and putative interacting residues on LILRB1 shown as sticks. The HLA heavy chain is coloured orange, β2M green, the bound peptide magenta, and LILRB1 blue.

**Figure S6.**
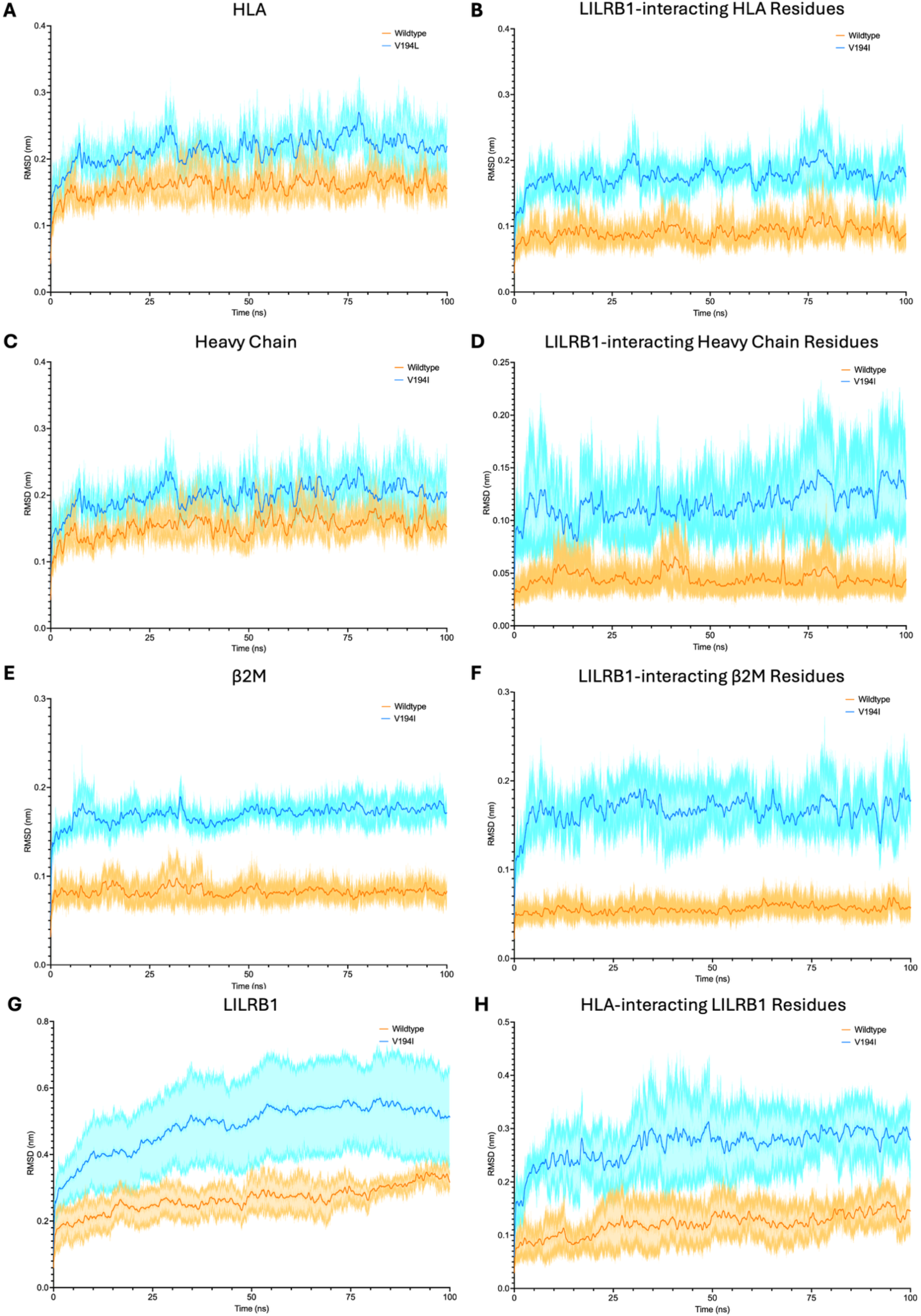
Root mean square deviation (RMSD) trajectories of LILRB1 in complex with wild-type HLA-B*35:01 (orange) or HLA-B*35:01 containing a valine-to-isoleucine point mutation at residue 194 of the heavy chain (blue). RMSD comparisons were performed for RMSD was calculated for **(A)** the entire HLA complex, **(B)** HLA residues contacting LILRB1, **(C)** the HLA heavy chain, **(D)** heavy-chain residues contacting LILRB1, **(E)** β2M, **(F)** β2M residues contacting LILRB1, **(G)** LILRB1, and **(H)** LILRB1 residues contacting HLA. Data are presented as mean ± SEM from six independent 100-ns molecular dynamics simulations for each system, with second-order smoothing applied over 500 neighbouring frames for clarity.

**Figure S7.**
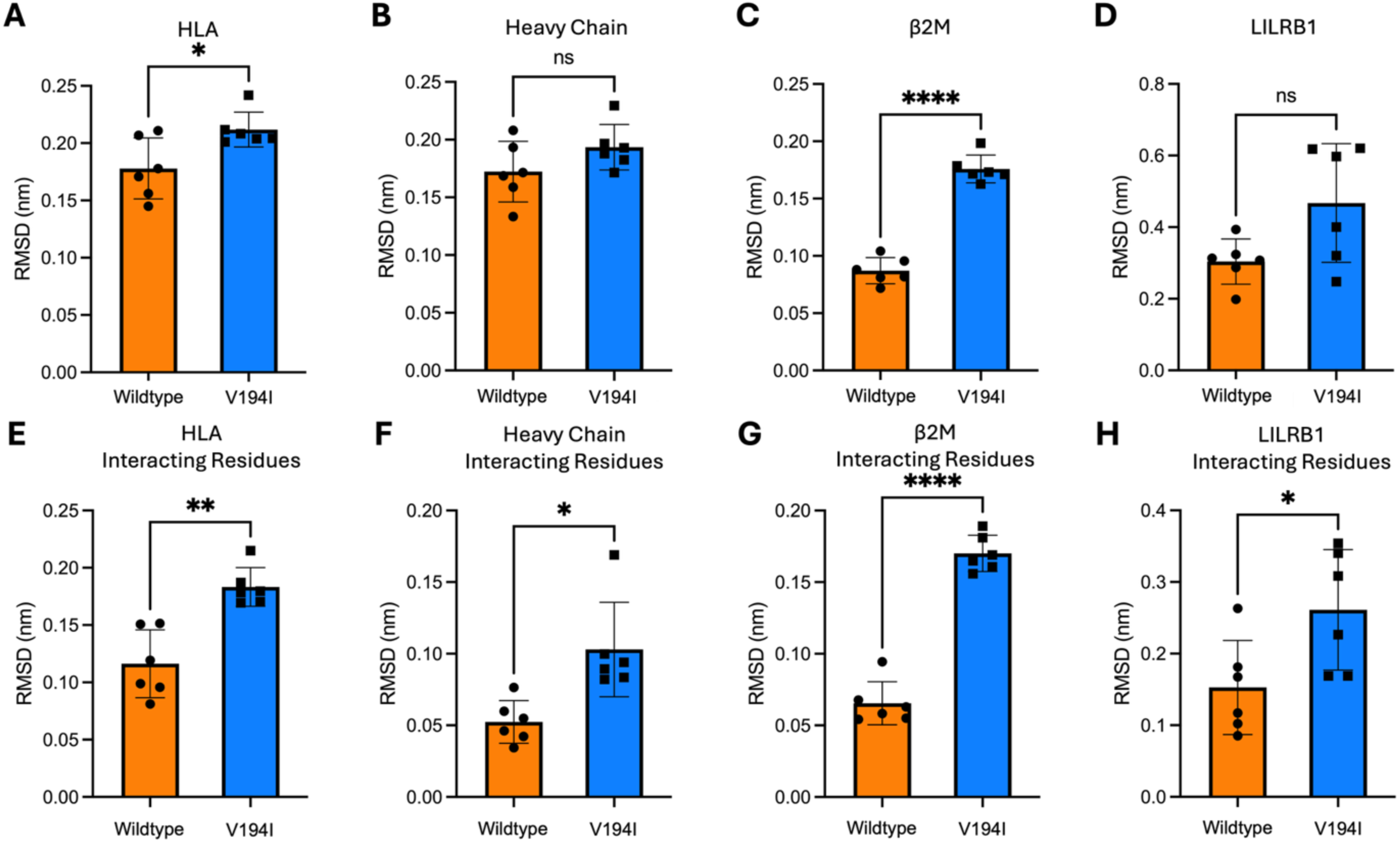
Averages of root mean square deviation (RMSD) values from simulations of LILRB1 in complex with wild-type HLA-B*35:01 (orange) or HLA-B*35:01 containing a valine-to-isoleucine point mutation at residue 194 of the heavy chain (blue). Comparisons are shown for **(A)** the whole HLA complex, **(B)** the heavy chain, **(C)** β2M, **(D)** LILRB1, and for the interacting residues of **(E)** HLA, **(F)** the heavy chain, **(G)** β2M, and **(H)** LILRB1. Interacting residues comprise heavy-chain residues 193-196, 198, and 248; β2M residues 1-4, 86-89, 91-93, and 96; and LILRB1 residues 18, 36, 38, 39, 41-43, 67, 68, 76, 97-100, 125-127, 184, and 187. Data are presented as mean ± SEM from si× 100-ns independent simulations of each system. Statistical analyses were performed using Welch’s t-test; *p<0.05, **p<0.01, ***p<0.001.

**Figure S8.**
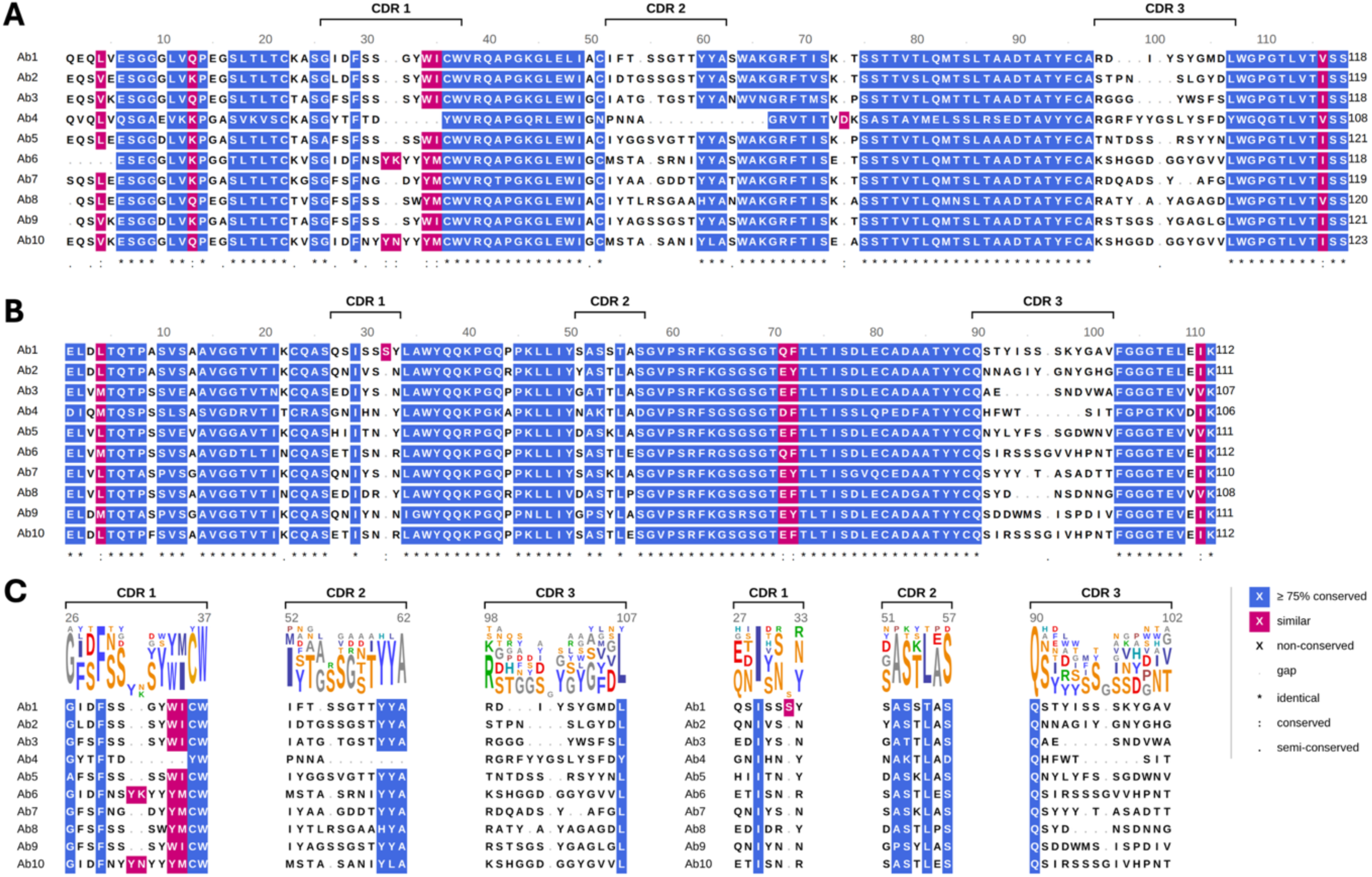
Sequence alignment of patent-derived antibodies modelled for interactions with LILRB1. **(A)** Aligned sequences of the antibody variable heavy chains, with complementarity-determining regions (CDRs) 1, 2, and 3 annotated. **(B)** Aligned sequences of the antibody variable light chains, with CDRs 1, 2, and 3 annotated. **(C)** Sequence logos of the heavy chain CDRs. **(D)** Sequence logos of the light chain CDRs. Dots represent semi-conservative substitutions, colons denote conservative substitutions, and asterisks indicate fully conserved residues. Residues highlighted in magenta are chemically similar while residues in blue are conserved in more than 75% of the 12 antibodies.

## Supplementary Tables

**Table S1.**
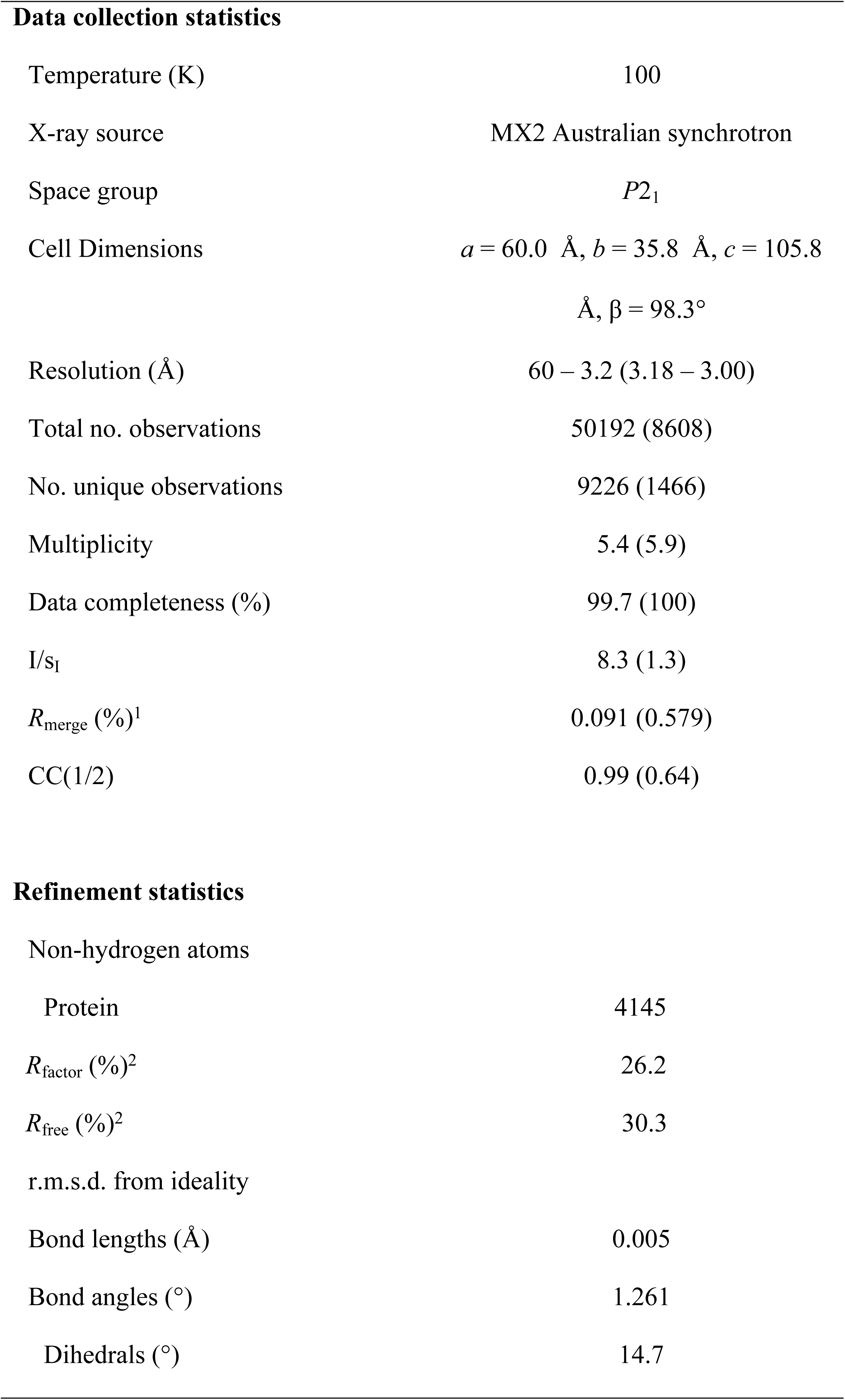
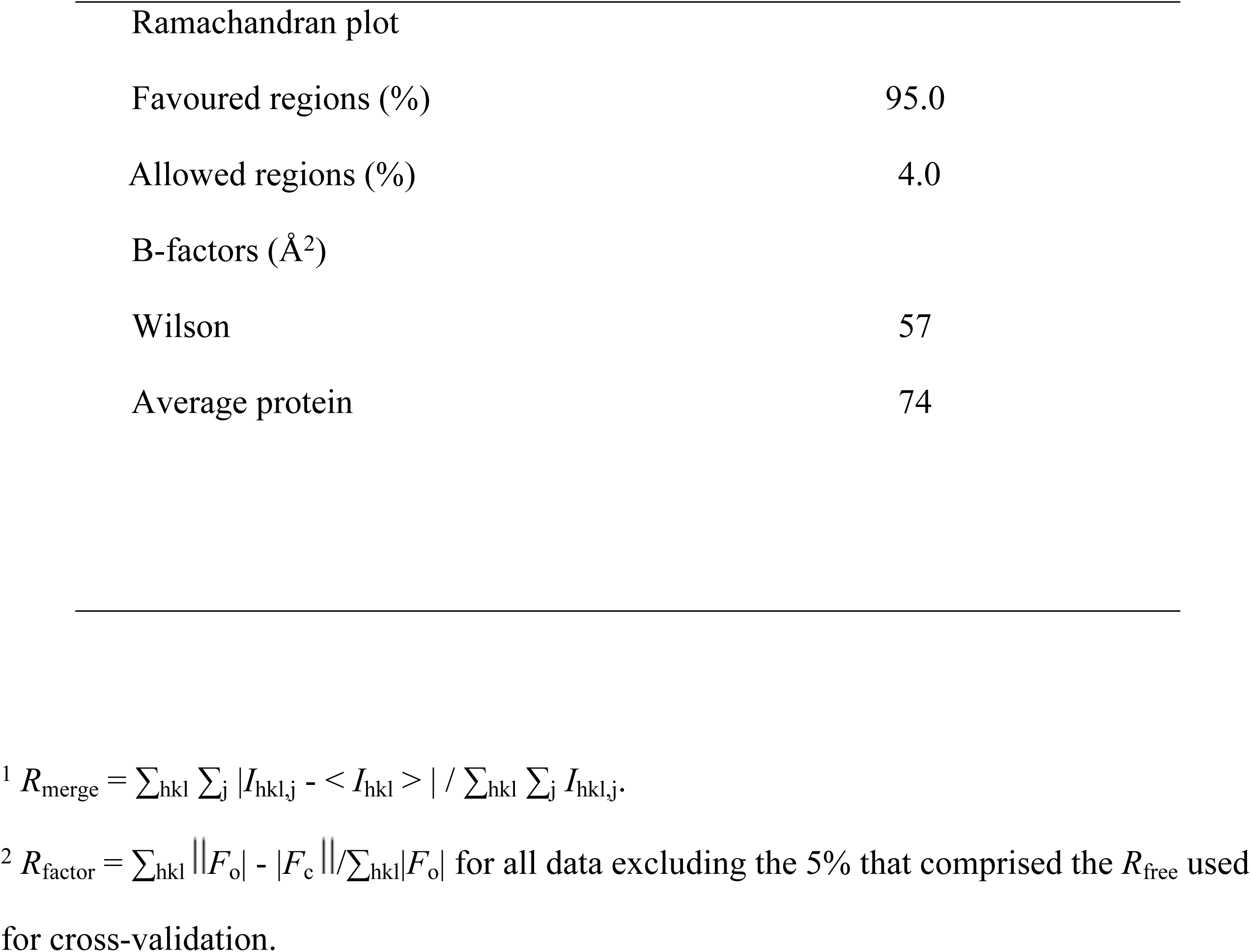
Data collection and refinement statistics for LILRB1.

| Data collection statistics |  |
| --- | --- |
| Temperature (K) | 100 |
| X-ray source | MX2 Australian synchrotron |
| Space group | P2<sub>1</sub> |
| Cell Dimensions | a = 60.0 Å, b = 35.8 Å, c = 105.8 Å, β = 98.3° |
| Resolution (Å) | 60 – 3.2 (3.18 – 3.00) |
| Total no. observations | 50192 (8608) |
| No. unique observations | 9226 (1466) |
| Multiplicity | 5.4 (5.9) |
| Data completeness (%) | 99.7 (100) |
| I/s<sub>I</sub> | 8.3 (1.3) |
| R<sub>merge</sub> (%)<sup>1</sup> | 0.091 (0.579) |
| CC(1/2) | 0.99 (0.64) |
| Refinement statistics |  |
| Non-hydrogen atoms |  |
| Protein | 4145 |
| R<sub>factor</sub> (%)<sup>2</sup> | 26.2 |
| R<sub>free</sub> (%)<sup>2</sup> | 30.3 |
| r.m.s.d. from ideality |  |
| Bond lengths (Å) | 0.005 |
| Bond angles (°) | 1.261 |
| Dihedrals (°) | 14.7 |
| Ramachandran plot |  |
| Favoured regions (%) | 95.0 |
| Allowed regions (%) | 4.0 |
| B-factors (Å<sup>2</sup>) |  |
| Wilson | 57 |
| Average protein | 74 |

<sup>1</sup> $R_{\text{merge}} = \sum_{\text{hkl}} \sum_j |I_{\text{hkl},j} - \langle I_{\text{hkl}} \rangle| / \sum_{\text{hkl}} \sum_j I_{\text{hkl},j}$ .
<sup>2</sup> $R_{\text{factor}} = \sum_{\text{hkl}} ||F_o| - |F_c|| / \sum_{\text{hkl}} |F_o|$ for all data excluding the 5% that comprised the $R_{\text{free}}$ used for cross-validation.

**Table S2:**
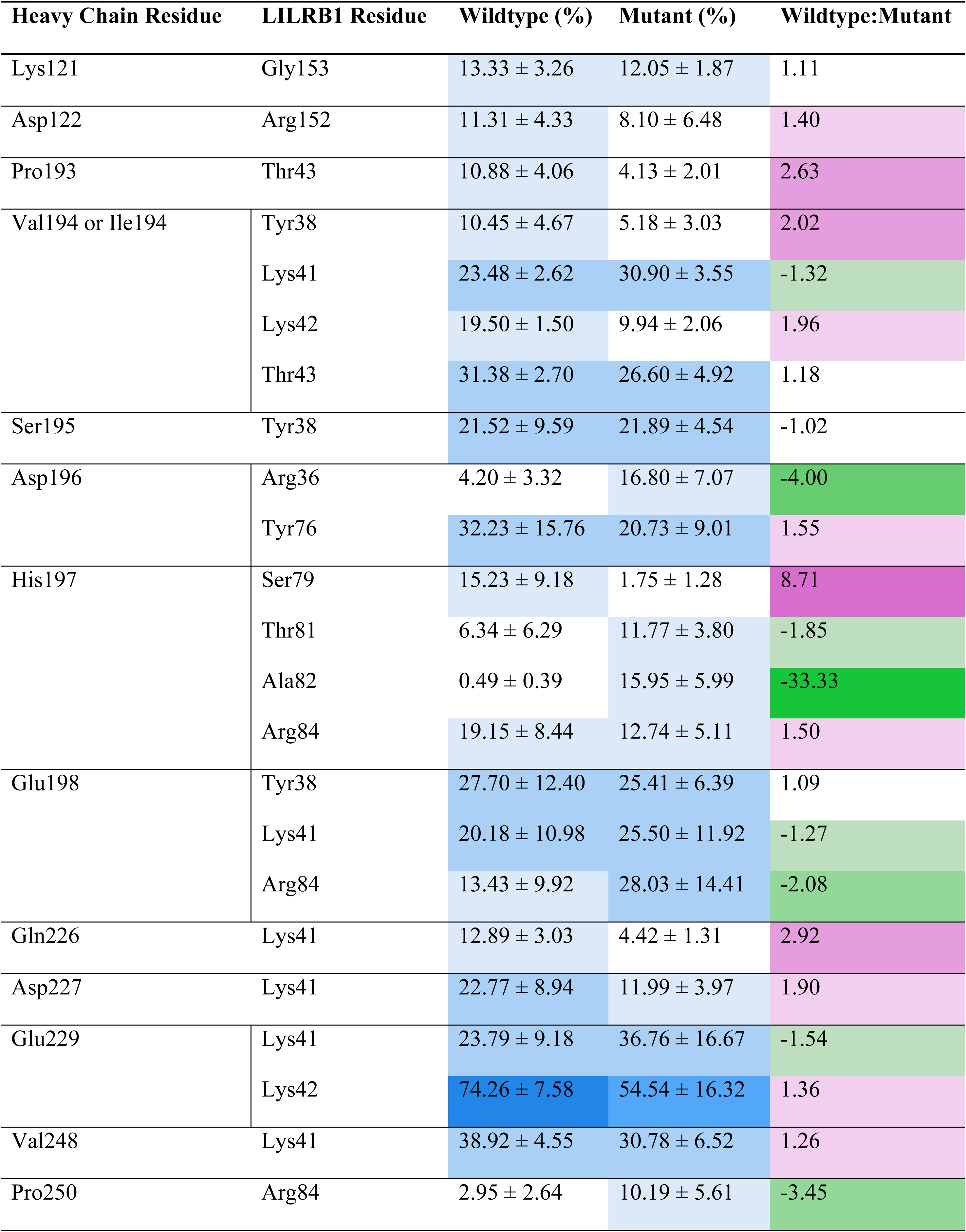
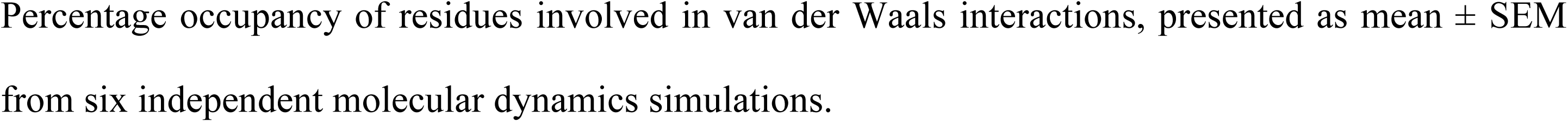
van der Waals interaction occupancy between wildtype and mutant HLA-B*35 heavy chain and LILRB1.

| Heavy Chain Residue | LILRB1 Residue | Wildtype (%) | Mutant (%) | Wildtype:Mutant |
| --- | --- | --- | --- | --- |
| Lys121 | Gly153 | 13.33 ± 3.26 | 12.05 ± 1.87 | 1.11 |
| Asp122 | Arg152 | 11.31 ± 4.33 | 8.10 ± 6.48 | 1.40 |
| Pro193 | Thr43 | 10.88 ± 4.06 | 4.13 ± 2.01 | 2.63 |
| Val194 or Ile194 | Tyr38 | 10.45 ± 4.67 | 5.18 ± 3.03 | 2.02 |
|  | Lys41 | 23.48 ± 2.62 | 30.90 ± 3.55 | -1.32 |
|  | Lys42 | 19.50 ± 1.50 | 9.94 ± 2.06 | 1.96 |
|  | Thr43 | 31.38 ± 2.70 | 26.60 ± 4.92 | 1.18 |
| Ser195 | Tyr38 | 21.52 ± 9.59 | 21.89 ± 4.54 | -1.02 |
| Asp196 | Arg36 | 4.20 ± 3.32 | 16.80 ± 7.07 | -4.00 |
|  | Tyr76 | 32.23 ± 15.76 | 20.73 ± 9.01 | 1.55 |
| His197 | Ser79 | 15.23 ± 9.18 | 1.75 ± 1.28 | 8.71 |
|  | Thr81 | 6.34 ± 6.29 | 11.77 ± 3.80 | -1.85 |
|  | Ala82 | 0.49 ± 0.39 | 15.95 ± 5.99 | -33.33 |
|  | Arg84 | 19.15 ± 8.44 | 12.74 ± 5.11 | 1.50 |
| Glu198 | Tyr38 | 27.70 ± 12.40 | 25.41 ± 6.39 | 1.09 |
|  | Lys41 | 20.18 ± 10.98 | 25.50 ± 11.92 | -1.27 |
|  | Arg84 | 13.43 ± 9.92 | 28.03 ± 14.41 | -2.08 |
| Gln226 | Lys41 | 12.89 ± 3.03 | 4.42 ± 1.31 | 2.92 |
| Asp227 | Lys41 | 22.77 ± 8.94 | 11.99 ± 3.97 | 1.90 |
| Glu229 | Lys41 | 23.79 ± 9.18 | 36.76 ± 16.67 | -1.54 |
|  | Lys42 | 74.26 ± 7.58 | 54.54 ± 16.32 | 1.36 |
| Val248 | Lys41 | 38.92 ± 4.55 | 30.78 ± 6.52 | 1.26 |
| Pro250 | Arg84 | 2.95 ± 2.64 | 10.19 ± 5.61 | -3.45 |

Percentage occupancy of residues involved in van der Waals interactions, presented as mean ± SEM from six independent molecular dynamics simulations.

**Table S3:** van der Waals interaction occupancy between wildtype and mutant HLA-B*35 β2M and LILRB1.

| β2M Residue | LILRB1 Residue | Wildtype (%) | Mutant (%) | Wildtype:Mutant |
| --- | --- | --- | --- | --- |
| Ile1 | Gln125 | 56.06 ± 2.53 | 56.47 ± 2.79 | -1.01 |
|  | Ala127 | 31.42 ± 3.54 | 32.96 ± 0.94 | -1.05 |
|  | Gly153 | 19.87 ± 2.26 | 8.42 ± 2.12 | 2.36 |
|  | Ser154 | 2.48 ± 0.90 | 10.27 ± 2.88 | -4.17 |
|  | Ser155 | 19.52 ± 1.98 | 12.22 ± 1.13 | 1.60 |
| Gln2 | Gln125 | 97.33 ± 0.83 | 97.33 ± 1.54 | 1.00 |
|  | Val126 | 75.26 ± 3.42 | 74.20 ± 1.63 | 1.01 |
|  | Ala127 | 99.57 ± 0.16 | 99.48 ± 0.15 | 1.00 |
| Arg3 | Ala127 | 32.26 ± 2.04 | 29.21 ± 2.15 | 1.10 |
|  | Asp178 | 0.16 ± 0.13 | 10.11 ± 6.47 | -50.00 |
| Thr4 | Tyr99 | 46.77 ± 1.39 | 47.22 ± 1.56 | -1.01 |
|  | Asp178 | 18.11 ± 2.12 | 18.96 ± 3.63 | -1.04 |
| Lys6 | Asp178 | 15.66 ± 15.13 | 6.94 ± 6.80 | 2.26 |
|  | Asn180 | 33.72 ± 2.52 | 29.37 ± 2.80 | 1.15 |
| Thr86 | Tyr99 | 67.97 ± 3.02 | 65.14 ± 0.98 | 1.04 |
|  | Ile100 | 91.37 ± 1.30 | 92.71 ± 1.75 | -1.01 |
|  | Val126 | 54.00 ± 1.41 | 56.47 ± 2.09 | -1.04 |
| Leu87 | Ala98 | 40.94 ± 1.20 | 42.84 ± 2.53 | -1.04 |
|  | Tyr99 | 31.67 ± 1.21 | 35.25 ± 2.53 | -1.11 |
| Ser88 | Gly97 | 61.08 ± 2.23 | 58.59 ± 1.27 | 1.04 |
|  | Ala98 | 88.73 ± 1.14 | 92.18 ± 0.82 | -1.04 |
|  | Tyr99 | 14.85 ± 1.53 | 15.36 ± 1.05 | -1.03 |
|  | Leu187 | 27.79 ± 1.49 | 33.16 ± 0.69 | -1.19 |
| Gln89 | Val95 | 0.08 ± 0.04 | 13.16 ± 6.34 | -100.00 |
|  | Thr96 | 0.08 ± 0.05 | 12.16 ± 5.82 | -100.00 |
|  | Gly97 | 47.80 ± 7.30 | 15.74 ± 7.30 | 3.04 |
|  | Ala98 | 18.01 ± 4.44 | 10.37 ± 4.95 | 1.74 |
|  | Leu187 | 0.02 ± 0.01 | 12.37 ± 6.06 | -Inf |
| Lys91 | Trp67 | 47.16 ± 9.00 | 26.50 ± 8.33 | 1.78 |
|  | Asn180 | 0.23 ± 0.20 | 49.89 ± 22.30 | -Inf |
|  | Glu184 | 98.28 ± 0.61 | 56.74 ± 19.37 | 1.73 |
| Ile92 | Trp67 | 58.92 ± 6.73 | 18.63 ± 4.81 | 3.16 |
| Val93 | Trp67 | 32.80 ± 8.75 | 8.41 ± 4.34 | 3.90 |
| Lys94 | Pro45 | 0.07 ± 0.06 | 34.86 ± 16.05 | -Inf |
|  | Trp67 | 4.15 ± 3.82 | 45.12 ± 12.98 | -11.11 |
|  | Glu68 | 55.69 ± 3.97 | 0.05 ± 0.05 | 1109.84 |
| Asp96 | Arg39 | 0.00 ± 0.00 | 13.59 ± 12.85 | -Inf |
| Met99 | Lys42 | 26.98 ± 6.72 | 10.92 ± 7.07 | 2.47 |

Percentage occupancy of residues involved in van der Waals interactions, presented as mean ± SEM from six independent molecular dynamics simulations.

**Table S4:** Hydrogen bond interaction occupancy between wildtype and mutant HLA-B*35 heavy chain and LILRB1.

| Heavy Chain Residue | LILRB1 Residue | Wildtype (%) | Mutant (%) | Wildtype:Mutant |
| --- | --- | --- | --- | --- |
| Asp196 | Arg36 (Donor) | 4.02 ± 3.20 | 15.61 ± 6.57 | -3.85 |
|  | Tyr76 (Donor) | 29.16 ± 14.64 | 16.47 ± 9.41 | 1.77 |
| Glu198 | Tyr38 (Donor) | 27.24 ± 12.21 | 25.08 ± 6.33 | 1.09 |
|  | Lys41 (Donor) | 19.84 ± 10.84 | 25.18 ± 11.80 | -1.27 |
|  | Arg84 (Donor) | 13.28 ± 9.84 | 24.28 ± 12.40 | -1.82 |
| Asp227 | Lys41 (Donor) | 15.76 ± 9.77 | 9.05 ± 4.02 | 1.74 |
| Glu229 | Lys41 (Donor) | 13.04 ± 5.22 | 30.89 ± 15.68 | -2.38 |
|  | Lys42 (Donor) | 73.71 ± 7.51 | 53.90 ± 16.14 | 1.37 |

Percentage occupancy of residues involved in hydrogen bond interactions, presented as mean ± SEM from six independent molecular dynamics simulations.

**Table S5:** Hydrogen bond interaction occupancy between wildtype and mutant HLA-B*35 β2M and LILRB1.

| β2M Residue | LILRB1 Residue | Wildtype (%) | Mutant (%) | Wildtype:Mutant |
| --- | --- | --- | --- | --- |
| Gln2 | Gln125 (Acceptor) | 93.87 ± 1.34 | 94.32 ± 2.07 | 1.00 |
|  | Ala127 (Donor) | 99.01 ± 0.13 | 98.60 ± 0.29 | 1.00 |
| Lys6 | Asp178 (Acceptor) | 15.13 ± 14.64 | 6.84 ± 6.75 | 2.21 |
| Thr86 | Ile100 (Donor) | 88.17 ± 1.64 | 89.90 ± 2.17 | -1.02 |
| Ser88 | Gly97 (Acceptor) | 12.45 ± 1.66 | 17.52 ± 3.87 | -1.41 |
|  | Ala98 (Acceptor) | 87.05 ± 1.31 | 90.41 ± 0.86 | -1.04 |
| Gln89 | Gly97 (Acceptor) | 42.58 ± 9.43 | 21.74 ± 3.60 | 1.96 |
|  | Ala98 (Acceptor) | 7.37 ± 5.44 | 13.93 ± 3.24 | -1.89 |
| Lys91 | Glu184 (Acceptor) | 96.88 ± 0.74 | 98.94 ± 0.26 | -1.02 |
| Ile92 | Trp67 (Donor) | 26.94 ± 7.02 | 5.61 ± 2.23 | 4.80 |
| Lys94 | Glu68 (Acceptor) | 53.78 ± 3.95 | 65.65 ± 7.73 | -1.22 |
| Asp96 | Lys42 (Donor) | 6.14 ± 4.45 | 25.54 ± 12.94 | -4.17 |
| Met99 | Lys42 (Donor) | 14.63 ± 9.03 | 0.00 ± 0.00 | 41699.56 |

Percentage occupancy of residues involved in hydrogen bond interactions, presented as mean ± SEM from six independent molecular dynamics simulations.

**Table S6:**
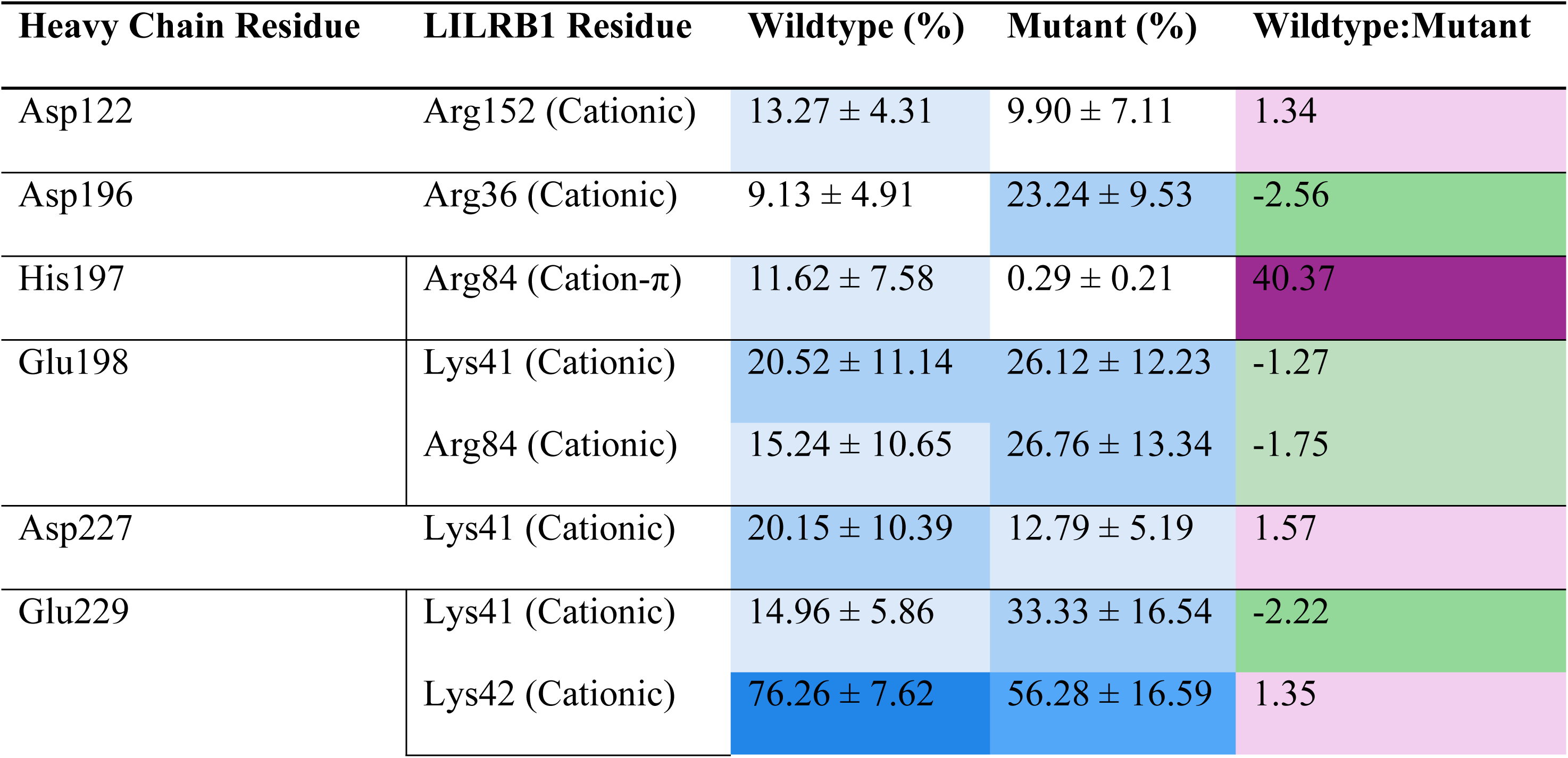
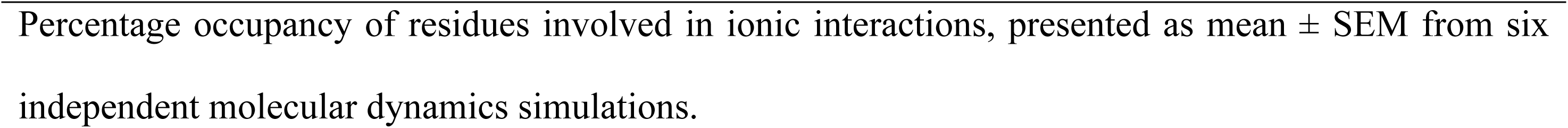
Ionic interaction occupancy between wildtype and mutant HLA-B*35 heavy chain and LILRB1.

| Heavy Chain Residue | LILRB1 Residue | Wildtype (%) | Mutant (%) | Wildtype:Mutant |
| --- | --- | --- | --- | --- |
| Asp122 | Arg152 (Cationic) | 13.27 ± 4.31 | 9.90 ± 7.11 | 1.34 |
| Asp196 | Arg36 (Cationic) | 9.13 ± 4.91 | 23.24 ± 9.53 | -2.56 |
| His197 | Arg84 (Cation-π) | 11.62 ± 7.58 | 0.29 ± 0.21 | 40.37 |
| Glu198 | Lys41 (Cationic) | 20.52 ± 11.14 | 26.12 ± 12.23 | -1.27 |
|  | Arg84 (Cationic) | 15.24 ± 10.65 | 26.76 ± 13.34 | -1.75 |
| Asp227 | Lys41 (Cationic) | 20.15 ± 10.39 | 12.79 ± 5.19 | 1.57 |
| Glu229 | Lys41 (Cationic) | 14.96 ± 5.86 | 33.33 ± 16.54 | -2.22 |
|  | Lys42 (Cationic) | 76.26 ± 7.62 | 56.28 ± 16.59 | 1.35 |

Percentage occupancy of residues involved in ionic interactions, presented as mean ± SEM from six independent molecular dynamics simulations.

**Table S7:** Hydrophobic interaction occupancy between wildtype and mutant HLA-B*35 heavy chain and LILRB1.

| Heavy Chain Residue | LILRB1 Residue | Wildtype (%) | Mutant (%) | Wildtype:Mutant |
| --- | --- | --- | --- | --- |
| Val194 or Ile194 | Tyr38 | 35.35 ± 15.90 | 19.45 ± 11.89 | 1.82 |
| | Lys41 | 3.49 ± 2.71 | 13.95 ± 4.18 | -4.00 |
| | Thr43 | 12.23 ± 4.20 | 23.09 ± 6.74 | -1.89 |
| Asp196 | Tyr76 | 23.49 ± 11.39 | 19.59 ± 8.81 | 1.20 |
| His197 | Tyr76 | 22.98 ± 10.52 | 6.93 ± 2.51 | 3.32 |
| | Ala82 | 0.78 ± 0.52 | 31.97 ± 12.86 | -50.00 |
| | Arg84 | 21.67 ± 9.99 | 17.43 ± 8.73 | 1.24 |
| Glu198 | Arg84 | 1.52 ± 1.52 | 21.92 ± 13.13 | -14.29 |
| Glu229 | Lys41 | 34.61 ± 15.09 | 18.64 ± 8.94 | 1.86 |
| Val248 | Lys41 | 76.09 ± 8.45 | 68.30 ± 11.30 | 1.11 |
| Pro250 | Arg84 | 4.08 ± 4.07 | 14.90 ± 9.21 | -3.70 |
| Percentage occupancy of residues involved in hydrophobic interactions, presented as mean ± SEM from six independent molecular dynamics simulations. |  |  |  |  |

**Table S8:** Hydrophobic interaction occupancy between wildtype and mutant HLA-B*35 β2M and LILRB1.

| β2M Residue | LILRB1 Residue | Wildtype (%) | Mutant (%) | Wildtype:Mutant |
| --- | --- | --- | --- | --- |
| Ile1 | Ala127 | 90.50 ± 1.49 | 89.19 ± 1.76 | 1.01 |
| Gln2 | Gln125 | 24.19 ± 5.45 | 31.89 ± 4.42 | -1.32 |
|  | Val126 | 69.03 ± 5.22 | 60.02 ± 1.44 | 1.15 |
| Arg3 | Ala127 | 69.17 ± 10.98 | 70.39 ± 4.75 | -1.02 |
| Thr4 | Tyr99 | 96.35 ± 0.41 | 96.51 ± 0.41 | 1.00 |
|  | Asp178 | 25.00 ± 7.40 | 18.98 ± 6.16 | 1.32 |
| Lys6 | Asn180 | 46.37 ± 5.50 | 43.31 ± 5.29 | 1.07 |
| Thr86 | Val126 | 99.84 ± 0.02 | 99.80 ± 0.02 | 1.00 |
| Leu87 | Tyr99 | 91.54 ± 2.28 | 88.67 ± 2.89 | 1.03 |
| Lys91 | Trp67 | 57.56 ± 13.40 | 22.11 ± 6.38 | 2.60 |
| Ile92 | Trp67 | 16.07 ± 14.41 | 0.17 ± 0.04 | 96.60 |
| Val93 | Trp67 | 65.18 ± 15.39 | 40.73 ± 9.55 | 1.60 |
| Lys94 | Trp67 | 11.07 ± 10.07 | 0.39 ± 0.20 | 28.16 |
|  | Glu68 | 5.21 ± 2.57 | 19.24 ± 5.15 | -3.70 |
| Met99 | Lys42 | 36.07 ± 15.94 | 35.57 ± 14.26 | 1.01 |

Percentage occupancy of residues involved in hydrophobic interactions, presented as mean ± SEM from six independent molecular dynamics simulations.

**Table S9:**
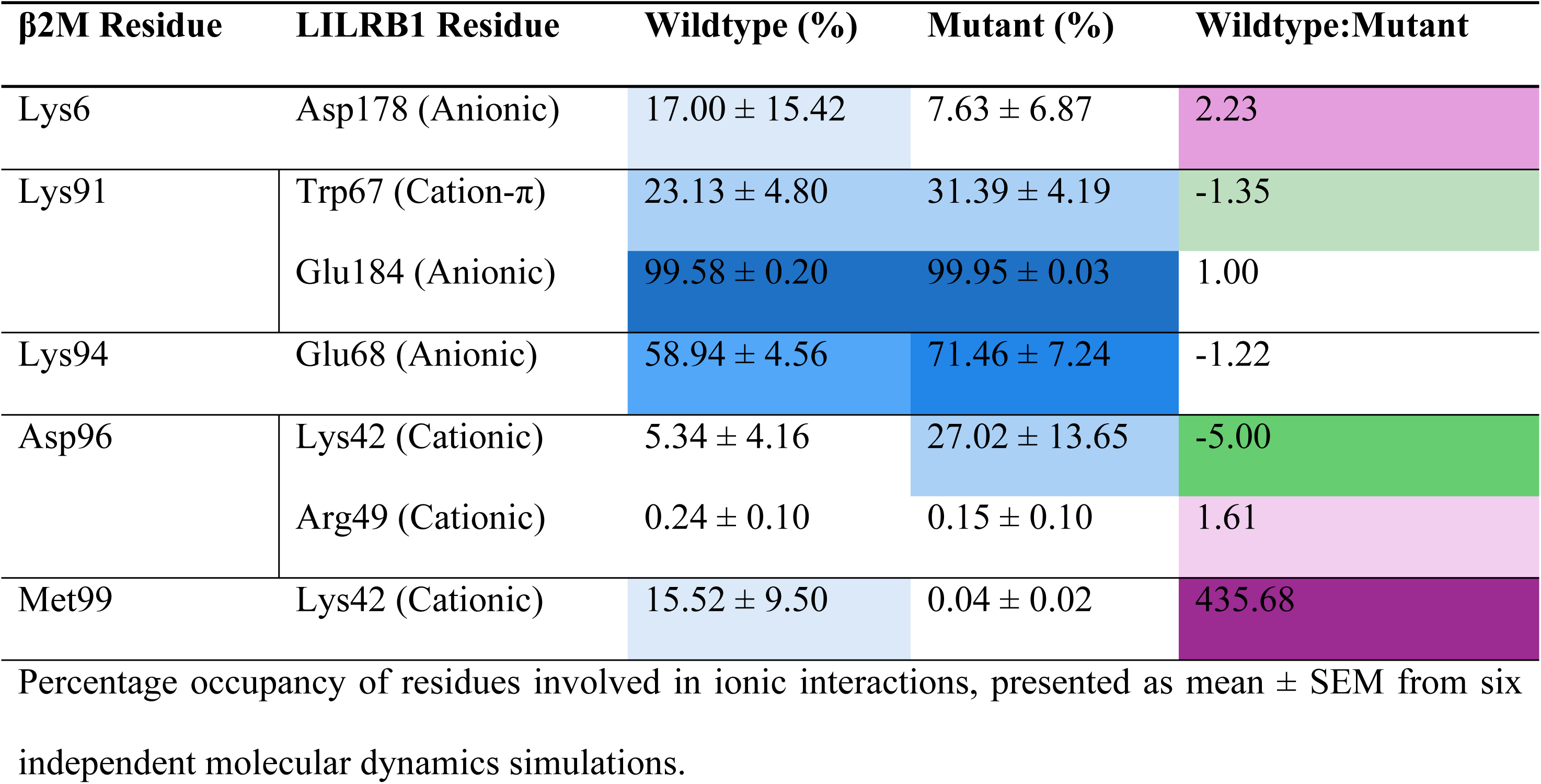
Ionic interaction occupancy between wildtype and mutant HLA-B*35 β2M and LILRB1.

| β2M Residue | LILRB1 Residue | Wildtype (%) | Mutant (%) | Wildtype:Mutant |
| --- | --- | --- | --- | --- |
| Lys6 | Asp178 (Anionic) | 17.00 ± 15.42 | 7.63 ± 6.87 | 2.23 |
| Lys91 | Trp67 (Cation-π) | 23.13 ± 4.80 | 31.39 ± 4.19 | -1.35 |
|  | Glu184 (Anionic) | 99.58 ± 0.20 | 99.95 ± 0.03 | 1.00 |
| Lys94 | Glu68 (Anionic) | 58.94 ± 4.56 | 71.46 ± 7.24 | -1.22 |
| Asp96 | Lys42 (Cationic) | 5.34 ± 4.16 | 27.02 ± 13.65 | -5.00 |
|  | Arg49 (Cationic) | 0.24 ± 0.10 | 0.15 ± 0.10 | 1.61 |
| Met99 | Lys42 (Cationic) | 15.52 ± 9.50 | 0.04 ± 0.02 | 435.68 |

Percentage occupancy of residues involved in ionic interactions, presented as mean ± SEM from six independent molecular dynamics simulations.

**Table S10:**
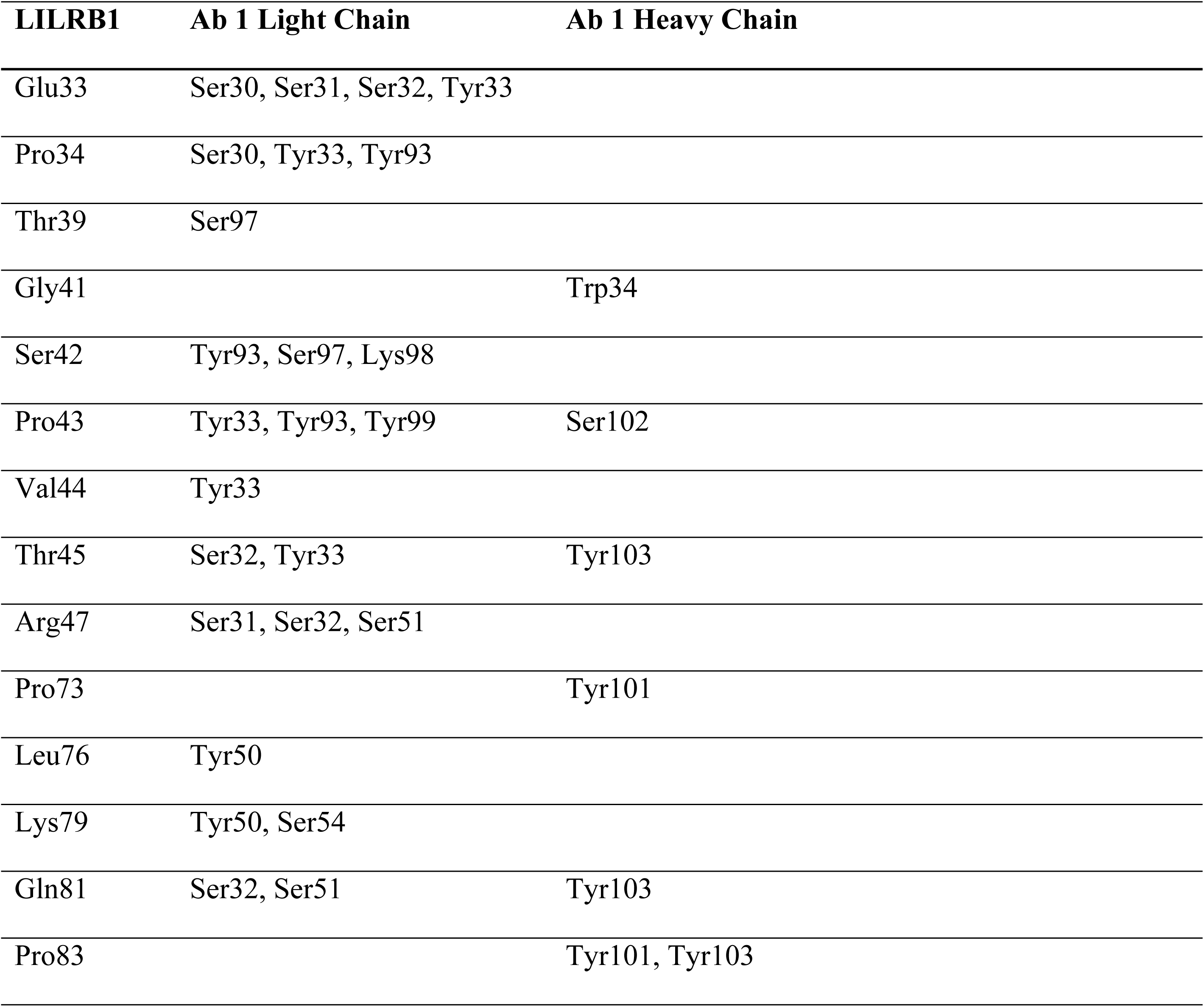
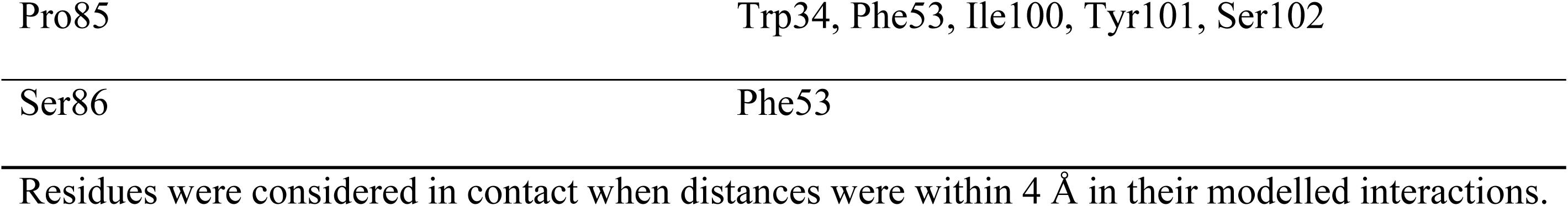
Modelled residue contacts between LILRB1 and antibody 1.

**Table S11:**
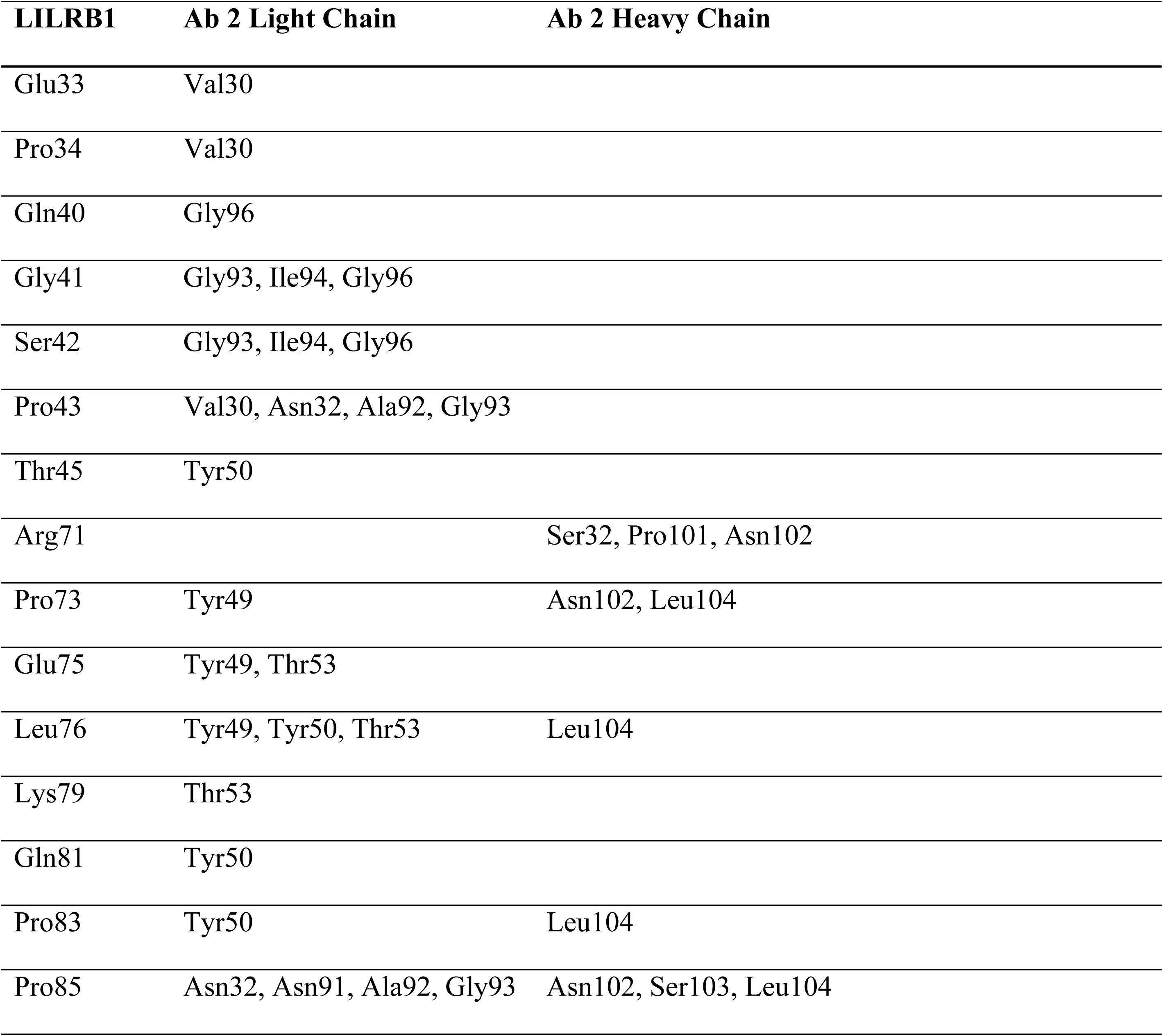
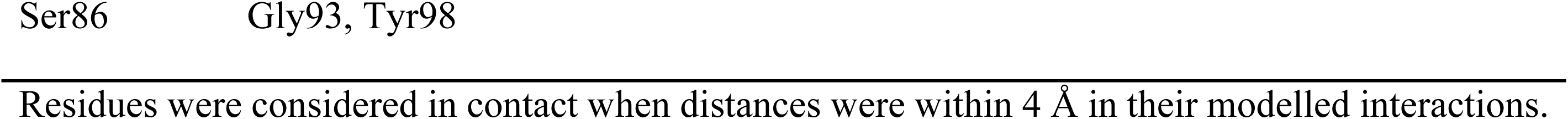
Modelled residue contacts between LILRB1 and antibody 2.

**Table S12:**
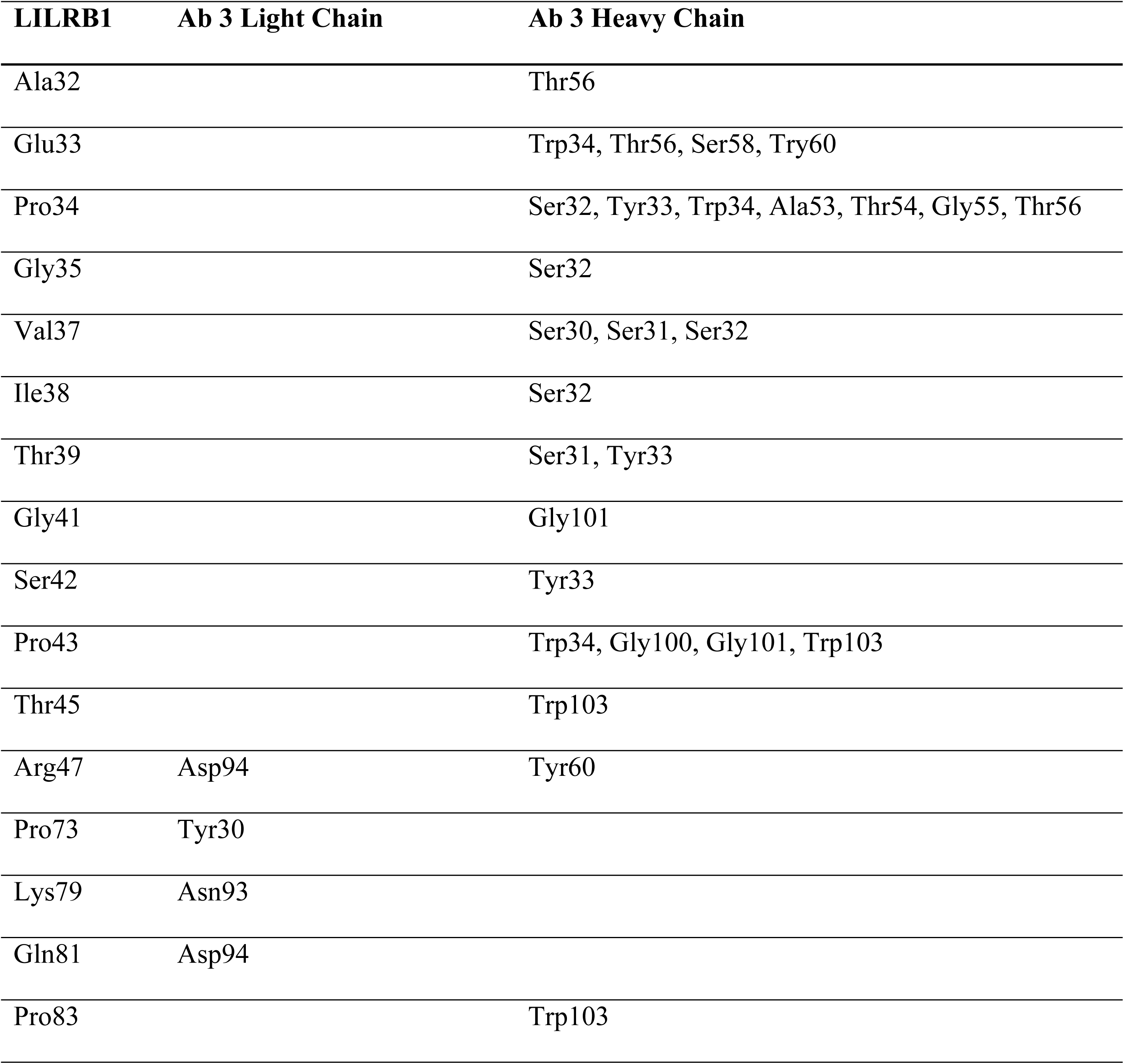
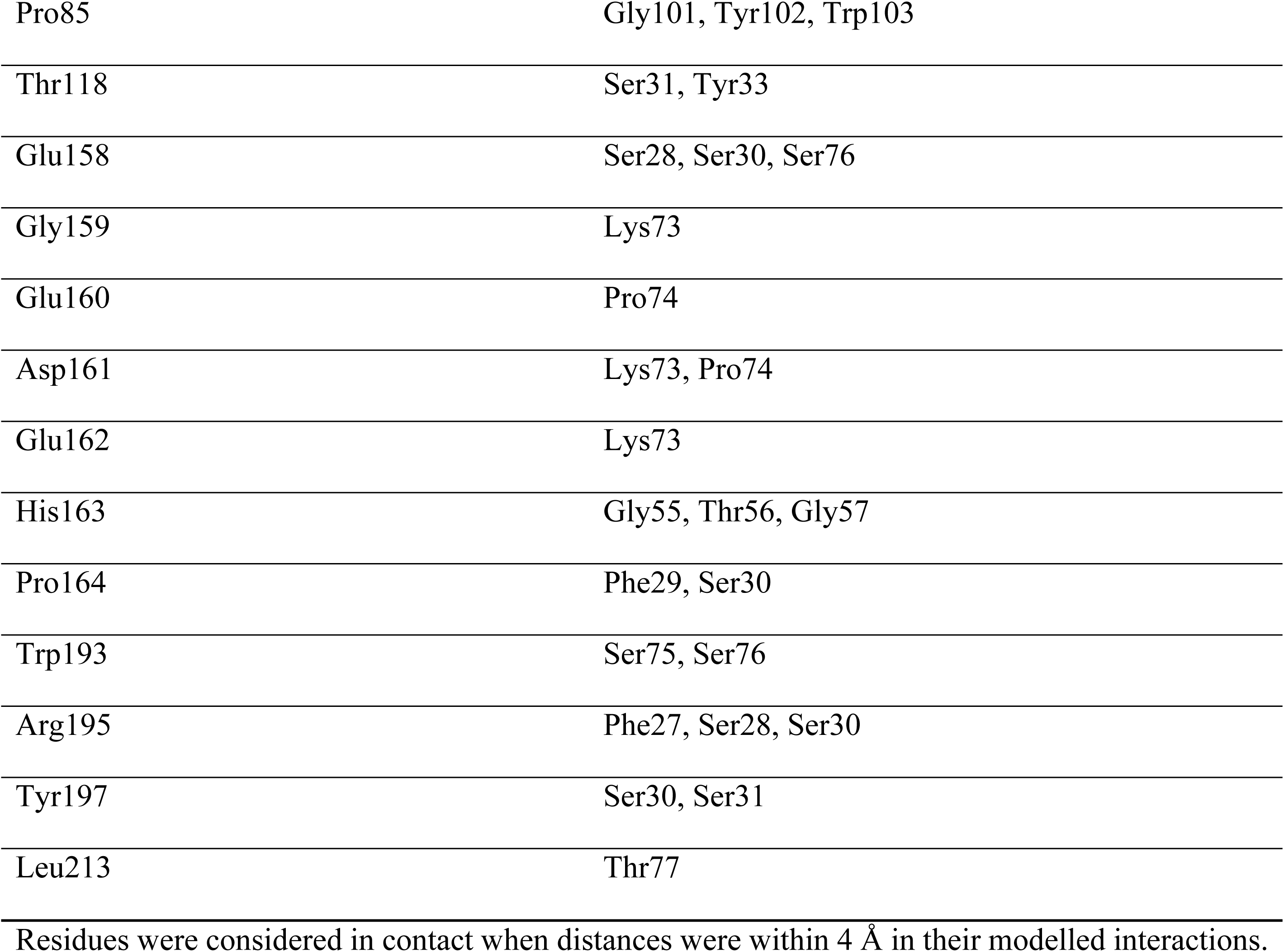
Modelled residue contacts between LILRB1 and antibody 3.

**Table S13:** Modelled residue contacts between LILRB1 and antibody 4.

| LILRB1 | Ab 4 Light Chain | Ab 4 Heavy Chain |
| --- | --- | --- |
| Ala120 | Pro48 | Asp1 |
| Tyr121 | Pro48 | Thr93 |
| Ile122 | Trp44, Gly46, Pro48 |  |
| Gln147 | Tyr32, Trp44, Leu92 |  |
| Val148 |  | Phe91, Thr93, Ser94, Ile95 |
| Ala149 | Ser91 | Phe91, Trp92, Thr93 |
| Phe150 |  | Trp92, Thr93 |
| Asp151 |  | Trp92 |
| Arg174 |  | Tyr32 |
| Gly175 |  | Tyr32, Trp92 |
| Ser177 | Gly90, Ser91 |  |
| Asp200 |  | Trp92, Thr93 |

Residues were considered in contact when distances were within 4 Å in their modelled interactions.

**Table S14:**
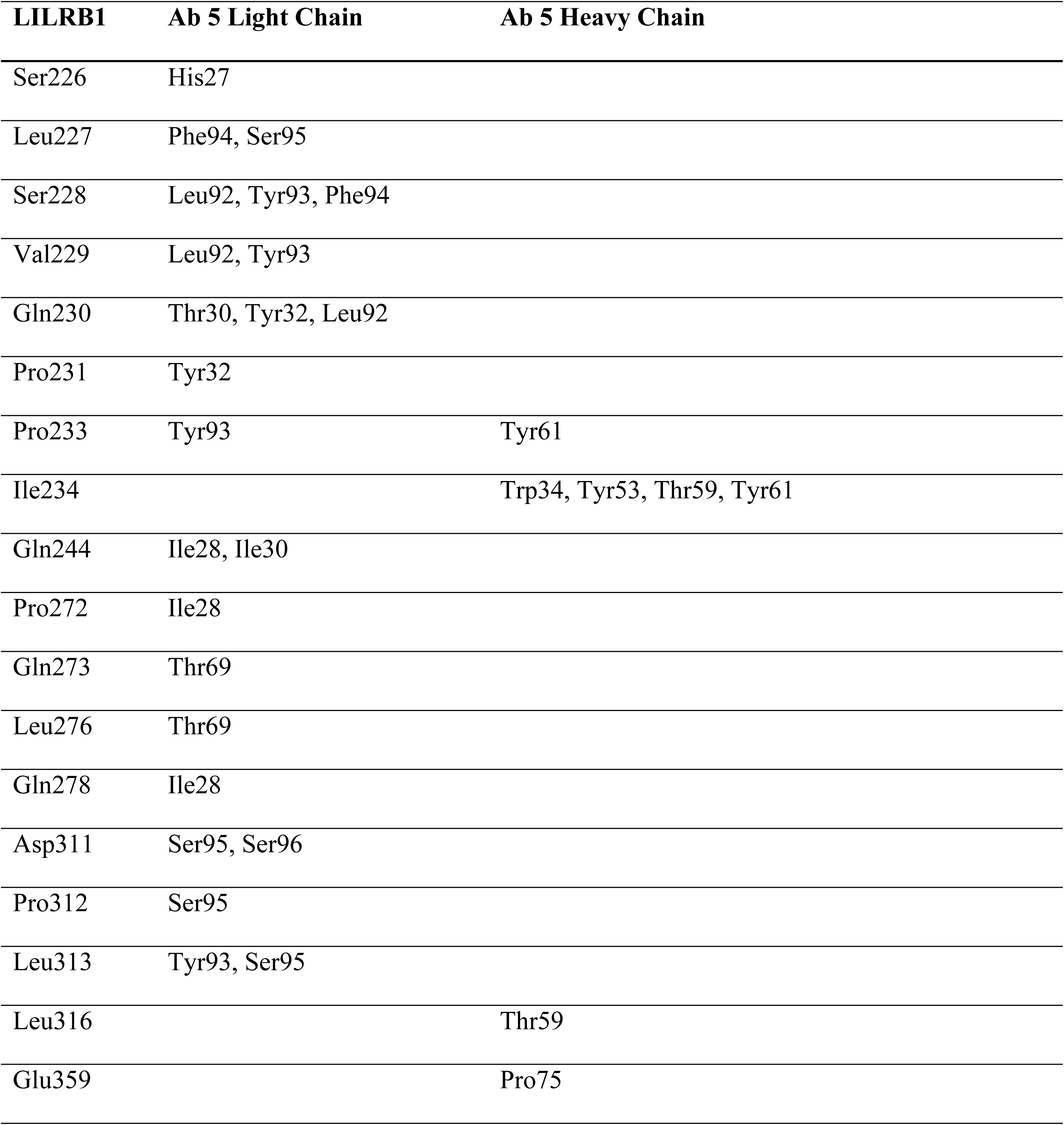
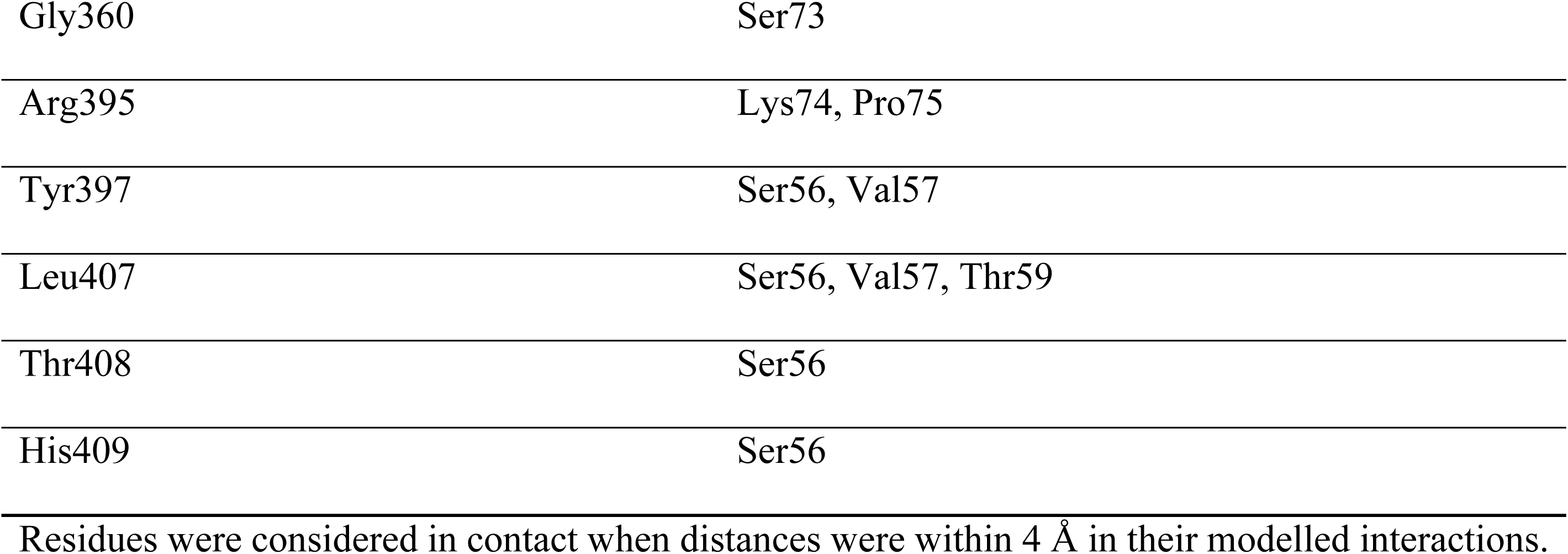
Modelled residue contacts between LILRB1 and antibody 5.

**Table S15:**
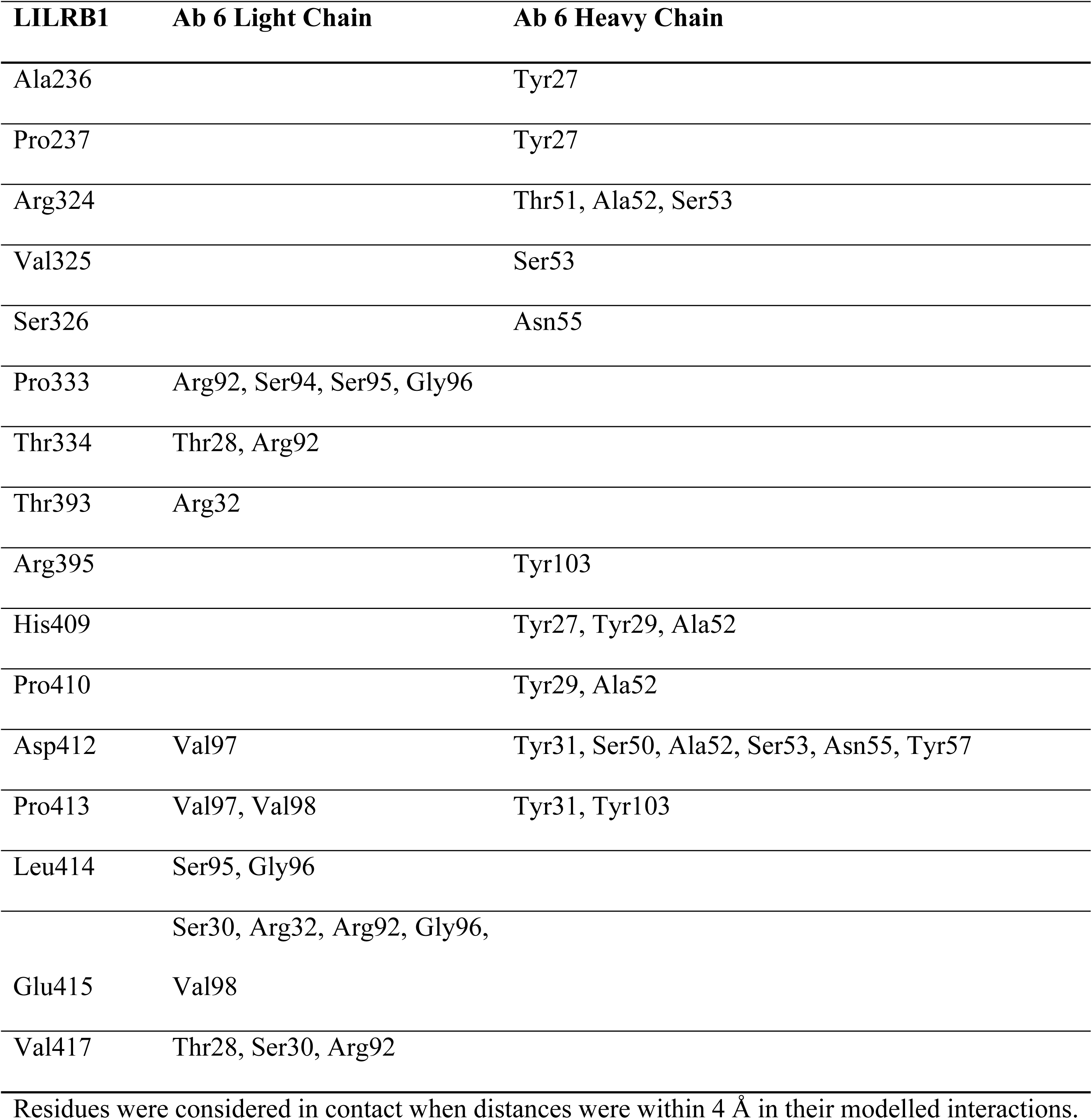
Modelled residue contacts between LILRB1 and antibody 6.

**Table S16:**
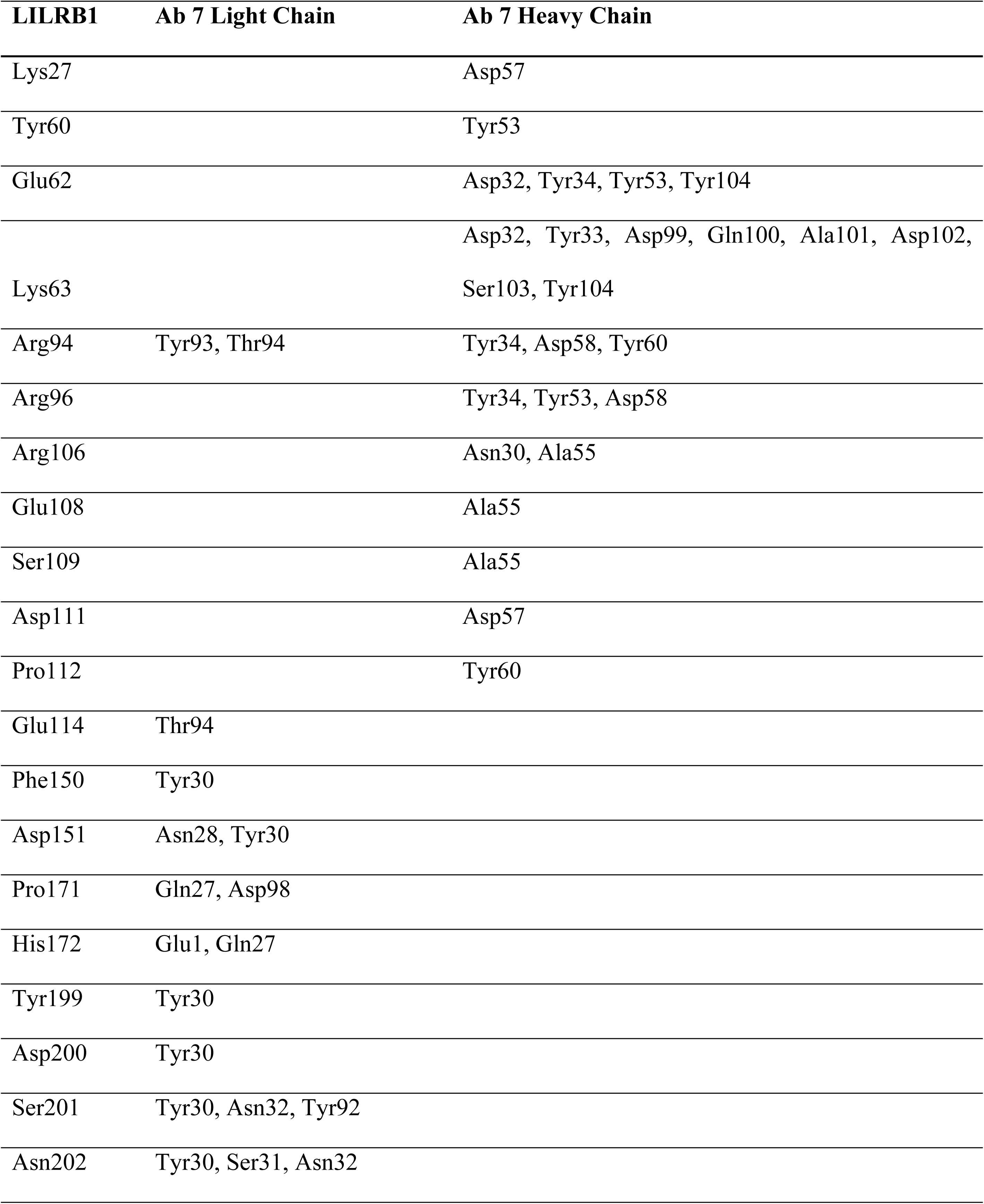
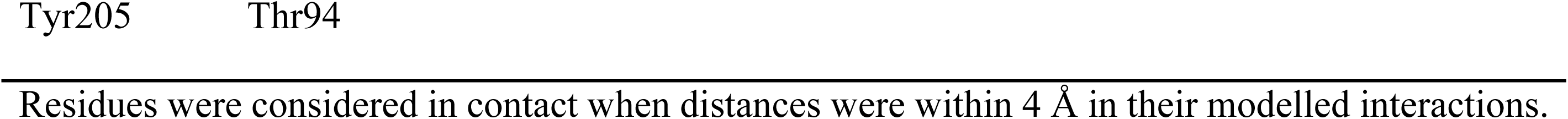
Modelled residue contacts between LILRB1 and antibody 7.

**Table S17:** Modelled residue contacts between LILRB1 and antibody 8.

| LILRB1 | Ab 8 Light Chain | Ab 8 Heavy Chain |
| --- | --- | --- |
| Pro231 |  | Arg55 |
| Ile234 |  | Arg55 |
| Val235 |  | Arg55 |
| Ala236 |  | Tyr33, Tyr52 |
| Pro237 |  | Tyr33 |
| Glu238 |  | Ala58, His60, Tyr103 |
| Glu239 |  | Tyr52, Arg55, Ala58 |
| Arg288 | Tyr32, Asp92 |  |
| Gly319 | Tyr101 |  |
| Gln320 | Tyr101 |  |
| Phe321 | Arg31 | Thr100, Tyr101 |
| Tyr322 | Val49, Pro55 | Thr100, Tyr101 |
| Arg324 | Ser31, Trp32 |  |
| Arg395 |  | Ser29, Leu54 |
| Lys403 | Arg31, Tyr32 |  |
| His409 |  | Ser31, Thr100 |
| Pro410 |  | Ser31, Leu54 |
| Asp412 |  | Ser30 |
| Pro413 |  | Ser30 |

Residues were considered in contact when distances were within 4 Å in their modelled interactions.

**Table S18:**
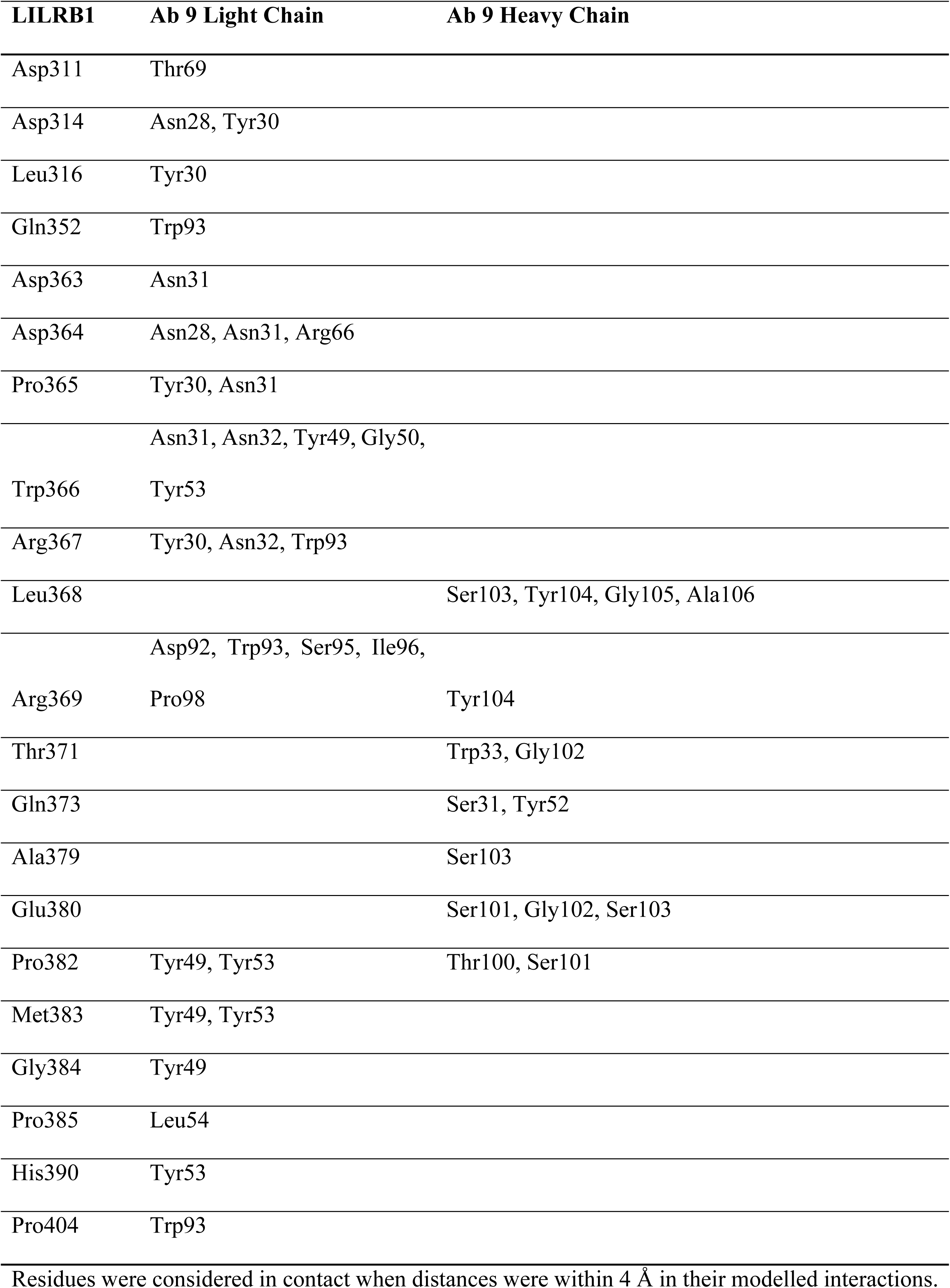
Modelled residue contacts between LILRB1 and antibody 9.

**Table S19:**
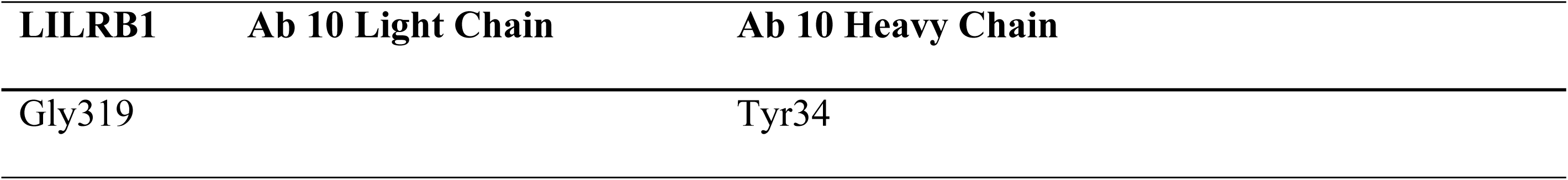
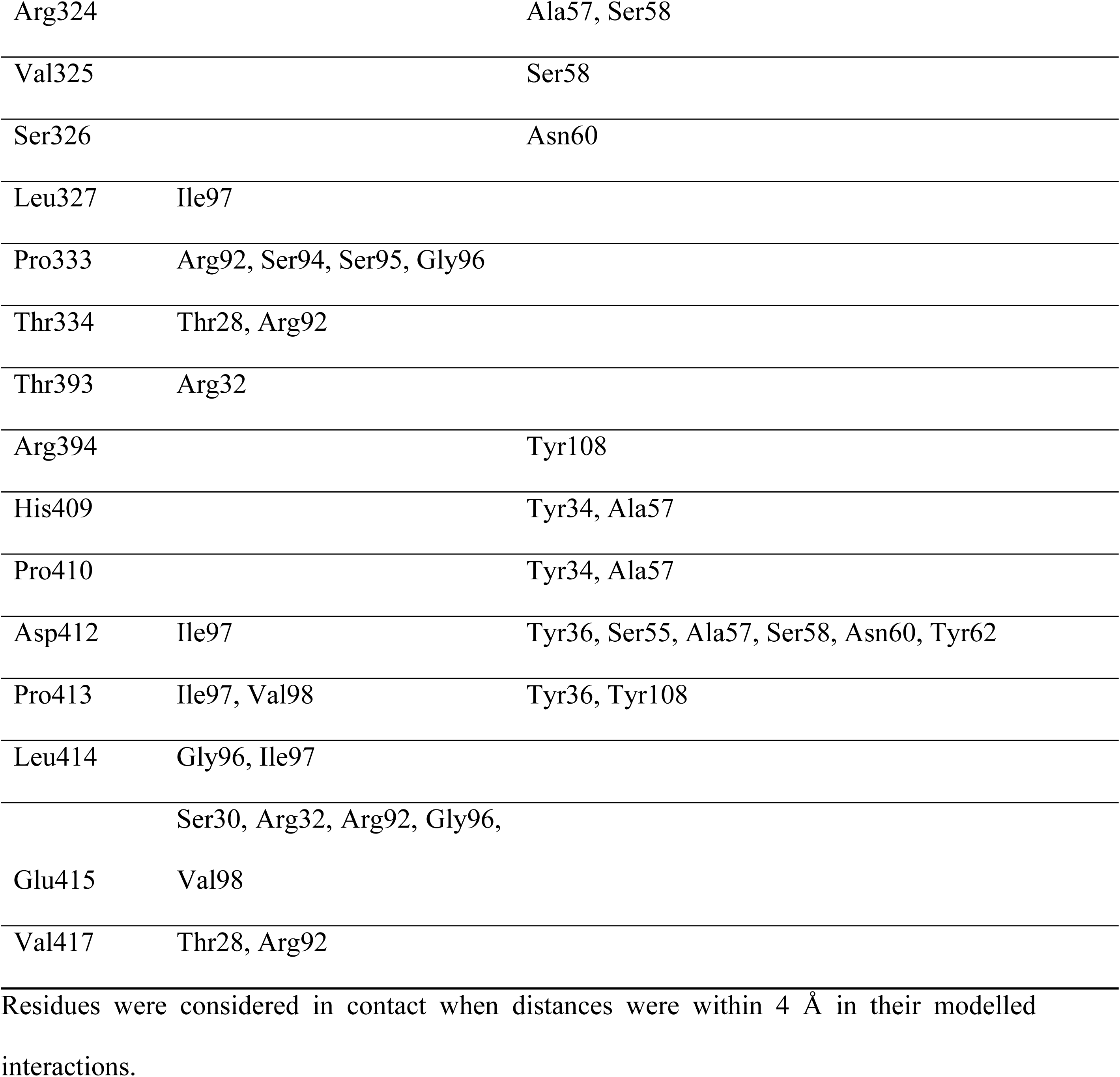
Modelled residue contacts between LILRB1 and antibody 10.

## References

Abraham, M. J., Murtola, T., Schulz, R., Páll, S., Smith, J. C., Hess, B. C Lindahl, E. 2015. GROMACS: High performance molecular simulations through multi-level parallelism from laptops to supercomputers. Softwarex, 1-2, 19–25.

Adams, P. D., Afonine, P. V., Bunkoczi, G., Chen, V. B., Davis, I. W., Echols, N., Headd, J. J., Hung, L.-W., Kapral, G. J., Grosse-Kunstleve, R. W., Mccoy, A. J., Moriarty, N. W., Oeffner, R., Read, R. J., Richardson, D. C., Richardson, J. S., Terwilliger, T. C. C Zwart, P. H. 2010. PHENIX: a comprehensive Python-based system for macromolecular structure solution. Acta Crystallographica Section D, 66, 213–221.

An, Z. Z., Chengcheng; Zhang, Ningyan; Chen, Yuanzhi; Chen, Heyu. 2021. Monoclonal antibodies against lilrb1 for diagnostic and therapeutic use. WO patent application.

Barkal, A. A., Weiskopf, K., Kao, K. S., Gordon, S. R., Rosental, B., Yiu, Y. Y., George, B. M., Markovic, M., Ring, N. G., Tsai, J. M., Mckenna, K. M., Ho, P. Y., Cheng, R. Z., Chen, J. Y., Barkal, L. J., Ring, A. M., Weissman, I. L. C Maute, R. L. 2018. Engagement of MHC class I by the inhibitory receptor LILRB1 suppresses macrophages and is a target of cancer immunotherapy. Nat Immunol, 19, 76–84.

Bouysset, C. C Fiorucci, S. 2021. ProLIF: a library to encode molecular interactions as fingerprints. Journal of cheminformatics, 13, 72.

Boyington, J. C., Motyka, S. A., Schuck, P., Brooks, A. G. C Sun, P. D. 2000. Crystal structure of an NK cell immunoglobulin-like receptor in complex with its class I MHC ligand. Nature, 405, 537–543.

Brondijk, T. H., De Ruiter, T., Ballering, J., Wienk, H., Lebbink, R. J., VAN Ingen, H., Boelens, R., Farndale, R. W., Meyaard, L. C Huizinga, E. G. 2010. Crystal structure and collagen-binding site of immune inhibitory receptor LAIR-1: unexpected implications for collagen binding by platelet receptor GPVI. Blood, 115, 1364–73.

Chapman, T. L., Heikema, A. P., West, A. P., Jr. C Bjorkman, P. J. 2000. Crystal structure and ligand binding properties of the D1D2 region of the inhibitory receptor LIR-1 (ILT2). Immunity, 13, 727–36.

Chapman, T. L., Heikeman, A. P. C Bjorkman, P. J. 1999. The inhibitory receptor LIR-1 uses a common binding interaction to recognize class I MHC molecules and the viral homolog UL18. Immunity, 11, 603–13.

Chen, H., Chen, Y., Deng, M., John, S., Gui, X., Kansagra, A., Chen, W., Kim, J., Lewis, C., Wu, G., Xie, J., Zhang, L., Huang, R., Liu, X., Arase, H., Huang, Y., Yu, H., Luo, W., Xia, N., Zhang, N., An, Z. C Zhang, C. C. 2020. Antagonistic anti-LILRB1 monoclonal antibody regulates antitumor functions of natural killer cells. Journal for ImmunoTherapy of Cancer, 8, e000515.

Chen, Y., Xu, K., Piccoli, L., Foglierini, M., Tan, J., Jin, W., Gorman, J., Tsybovsky, Y., Zhang, B., Traore, B., Silacci-Fregni, C., Daubenberger, C., Crompton, P. D., Geiger, R., Sallusto, F., Kwong, P. D. C Lanzavecchia, A. 2021. Structural basis of malaria RIFIN binding by LILRB1-containing antibodies. Nature, 592, 639–643.

Davidson, C. L., Li, N. L. C Burshtyn, D. N. 2010. LILRB1 polymorphism and surface phenotypes of natural killer cells. Human Immunology, 71, 942–949.

Davis, I. W., Leaver-Fay, A., Chen, V. B., Block, J. N., Kapral, G. J., Wang, X., Murray, L. W., Arendall, W. B., 3rd, Snoeyink, J., Richardson, J. S. C Richardson, D. C. 2007. MolProbity: all-atom contacts and structure validation for proteins and nucleic acids. Nucleic Acids Res, 35, W375–83.

Dédier, S., Reinelt, S., Reitinger, T., Folkers, G. C Rognan, D. 2000. Thermodynamic Stability of HLA-B*2705C#xb7;Peptide Complexes: EFFECT OF PEPTIDE AND MAJOR HISTOCOMPATIBILITY COMPLEX PROTEIN MUTATIONS *. Journal of Biological Chemistry, 275, 27055–27061.

Delano, W. L. 2002. Pymol: An open-source molecular graphics tool. CCP4 Newsl. protein crystallogr, 40, 82–92.

Deng, M., Chen, H., Liu, X., Huang, R., He, Y., Yoo, B., Xie, J., John, S., Zhang, N., An, Z. C Zhang, C. C. 2021. Leukocyte immunoglobulin-like receptor subfamily B: therapeutic targets in cancer. Antib Ther, 4, 16–33.

Dietrich, J., Cella, M. C Colonna, M. 2001. Ig-Like Transcript 2 (ILT2)/Leukocyte Ig-Like Receptor 1 (LIR1) Inhibits TCR Signaling and Actin Cytoskeleton Reorganization1. The Journal of Immunology, 166, 2514–2521.

Duey, D. Y. M. E., Allen James JR.; Kaplan, Daniel David; Lam, Chia-Ying Kao; Mondal, Kalyani; Stone, Geoffrey William; Wang, YAN. 2021. Ilt-binding agents and methods of use thereof. WO patent application.

Dulberger, C. L., Mcmurtrey, C. P., Hölzemer, A., Neu, K. E., Liu, V., Steinbach, A. M., Garcia-Beltran, W. F., Sulak, M., Jabri, B., Lynch, V. J., Altfeld, M., Hildebrand, W. H. C Adams, E. J. 2017. Human Leukocyte Antigen F Presents Peptides and Regulates Immunity through Interactions with NK Cell Receptors. Immunity, 46, 1018–1029.e7.

Ellis, T. M. 2013. Interpretation of HLA single antigen bead assays. Transplantation Reviews, 27, 108–111.

Emsley, P. C Cowtan, K. 2004. Coot: model-building tools for molecular graphics. Acta Crystallographica Section D, 60, 2126–2132.

Evans, P. 2006. Scaling and assessment of data quality. Acta Crystallographica Section D, 62, 72–82.

Fukazawa, T., Hermann, E., Edidin, M., Wen, J., Huang, F., Kellner, H., Floege, J., Farahmandian, D., Williams, K. M. C Yu, D. T. 1994. The effect of mutant beta 2-microglobulins on the conformation of HLA-B27 detected by antibody and by CTL. J Immunol, 153, 3543–50.

Gras, S., Chen, Z., Miles, J. J., Liu, Y. C., Bell, M. J., Sullivan, L. C., Kjer-Nielsen, L., Brennan, R. M., Burrows, J. M., Neller, M. A., Khanna, R., Purcell, A. W., Brooks, A. G., Mccluskey, J., Rossjohn, J. C Burrows, S. R. 2010. Allelic polymorphism in the T cell receptor and its impact on immune responses. Journal of Experimental Medicine, 207, 1555–1567.

Harrison, T. E., Mørch, A. M., Felce, J. H., Sakoguchi, A., Reid, A. J., Arase, H., Dustin, M. L. C Higgins, M. K. 2020. Structural basis for RIFIN-mediated activation of LILRB1 in malaria. Nature, 587, 309–312.

Hu, Z., Zhang, Q., He, Z., Jia, X., Zhang, W. C Cao, X. 2024. MHC1/LILRB1 axis as an innate immune checkpoint for cancer therapy. *Frontiers in Immunology*, Volume 15–2024.

Huang, J., Rauscher, S., Nawrocki, G., Ran, T., Feig, M., De Groot, B. L., Grubmüller, H. C Mackerell Jr, A. D. 2017. CHARMM36m: an improved force field for folded and intrinsically disordered proteins. Nature methods, 14, 71–73.

Jappe, E. C., Garde, C., Ramarathinam, S. H., Passantino, E., Illing, P. T., Mifsud, N. A., Trolle, T., Kringelum, J. V., Croft, N. P. C Purcell, A. W. 2020. Thermostability profiling of MHC-bound peptides: a new dimension in immunopeptidomics and aid for immunotherapy design. Nat Commun, 11, 6305.

Jo, S., Kim, T., Iyer, V. G. C Im, W. 2008. CHARMM-GUI: A web-based graphical user interface for CHARMM. Journal of Computational Chemistry, 29, 1859–1865.

Jones, D. C., Kosmoliaptsis, V., Apps, R., Lapaǫue, N., Smith, I., Kono, A., Chang, C., Boyle, L. H., Taylor, C. J., Trowsdale, J. C Allen, R. L. 2011. HLA Class I Allelic Sequence and Conformation Regulate Leukocyte Ig-Like Receptor Binding. The Journal of Immunology, 186, 2990–2997.

Jumper, J., Evans, R., Pritzel, A., Green, T., Figurnov, M., Ronneberger, O., Tunyasuvunakool, K., Bates, R., Žídek, A. C Potapenko, A. 2021. Highly accurate protein structure prediction with AlphaFold. nature, 596, 583–589.

Kabsch, W. 2010. XDS. Acta Crystallogr D Biol Crystallogr, 66, 125–32.

Kuroki, K., Matsubara, H., Kanda, R., Miyashita, N., Shiroishi, M., Fukunaga, Y., Kamishikiryo, J., Fukunaga, A., Fukuhara, H., Hirose, K., Hunt, J. S., Sugita, Y., Kita, S., Ose, T. C Maenaka, K. 2019. Structural and Functional Basis for LILRB Immune Checkpoint Receptor Recognition of HLA-G Isoforms. The Journal of Immunology, 203, 3386–3394.

Kuroki, K., Tsuchiya, N., Shiroishi, M., Rasubala, L., Yamashita, Y., Matsuta, K., Fukazawa, T., Kusaoi, M., Murakami, Y., Takiguchi, M., Juji, T., Hashimoto, H., Kohda, D., Maenaka, K. C Tokunaga, K. 2005. Extensive polymorphisms of LILRB1 (ILT2, LIR1) and their association with HLA-DRB1 shared epitope negative rheumatoid arthritis. Human Molecular Genetics, 14, 2469–2480.

Leijonhufvud, C., Reger, R., Segerberg, F., Theorell, J., Schlums, H., Bryceson, Y. T., Childs, R. W. C Carlsten, M. 2021. LIR-1 educates expanded human NK cells and defines a unique antitumor NK cell subset with potent antibody-dependent cellular cytotoxicity. Clin Transl Immunology, 10, e1346.

Liu, F., Cocker, A. T. H., Pugh, J. L., Djaoud, Z., Parham, P. C Guethlein, L. A. 2022. Natural LILRB1 D1-D2 variants show frequency differences in populations and bind to HLA class I with various avidities. Immunogenetics, 74, 513–525.

Mccutcheon, J. A., Gumperz, J., Smith, K. D., Lutz, C. T. C Parham, P. 1995. Low HLA-C expression at cell surfaces correlates with increased turnover of heavy chain mRNA. Journal of Experimental Medicine, 181, 2085–2095.

Mohammed, F., Stones, D. H., C Willcox, B. E. 2019. Application of the immunoregulatory receptor LILRB1 as a crystallisation chaperone for human class I MHC complexes. J Immunol Methods, 464, 47–56.

Mohammed, F., Stones, D. H., Zarling, A. L., Willcox, C. R., Shabanowitz, J., Cummings, K. L., Hunt, D. F., Cobbold, M., Engelhard, V. H., C Willcox, B. E. 2017. The antigenic identity of human class I MHC phosphopeptides is critically dependent upon phosphorylation status. Oncotarget, 8, 54160–54172.

Naji, A., Menier, C., Morandi, F., Agaugué, S., Maki, G., Ferretti, E., Bruel, S., Pistoia, V., Carosella, E. D., C Rouas-Freiss, N. 2014. Binding of HLA-G to ITIM-bearing Ig-like transcript 2 receptor suppresses B cell responses. J Immunol, 192, 1536–46.

Nam, G., Shi, Y., Ryu, M., Wang, Q., Song, H., Liu, J., Yan, J., Qi, J., C Gao, G. F. 2013. Crystal structures of the two membrane-proximal Ig-like domains (D3D4) of LILRB1/B2: alternative models for their involvement in peptide-HLA binding. Protein Cell, 4, 761–70.

Nguyen, A. T., Lau, H. M. P., Sloane, H., Jayasinghe, D., Mifsud, N. A., Chatzileontiadou, D. S., Grant, E. J., Szeto, C., C Gras, S. 2022. Homologous peptides derived from influenza A, B and C viruses induce variable CD8(+) T cell responses with cross-reactive potential. Clin Transl Immunology, 11, e1422.

Probst-Kepper, M., Hecht, H.-J. R., Herrmann, H., Janke, V., Ocklenburg, F., Klempnauer, J. R., van den Eynde, B. J., C Weiss, S. 2004. Conformational Restraints and Flexibility of 14-Meric Peptides in Complex with HLA-B*35011. The Journal of Immunology, 173, 5610–5616.

Pymm, P., Saunders, P. M., Anand, S., Maclachlan, B. J., Faoro, C., Hitchen, C., Rossjohn, J., Brooks, A. G., C Vivian, J. P. 2024. The Structural Basis for Recognition of Human Leukocyte Antigen Class I Molecules by the Pan-HLA Antibody W6/32. J Immunol, 213, 876–885.

Robinson, J., Barker, D. J., Georgiou, X., Cooper, M. A., Flicek, P., C Marsh, S. G. E. 2020. IPD-IMGT/HLA Database. Nucleic Acids Res, 48, D948–d955.

Shiroishi, M., Kuroki, K., Tsumoto, K., Yokota, A., Sasaki, T., Amano, K., Shimojima, T., Shirakihara, Y., Rasubala, L., van der Merwe, P. A., Kumagai, I., Kohda, D., C Maenaka, K. 2006. Entropically Driven MHC Class I Recognition by Human Inhibitory Receptor Leukocyte Ig-like Receptor B1 (LILRB1/ILT2/CD85j). Journal of Molecular Biology, 355, 237–248.

Shiroishi, M., Tsumoto, K., Amano, K., Shirakihara, Y., Colonna, M., Braud, V. M., Allan, D. S., Makadzange, A., Rowland-Jones, S., Willcox, B., Jones, E. Y., van der Merwe, P. A., Kumagai, I., C Maenaka, K. 2003. Human inhibitory receptors Ig-like transcript 2 (ILT2) and ILT4 compete with CD8 for MHC class I binding and bind preferentially to HLA-G. Proc Natl Acad Sci U S A, 100, 8856–61.

Swift, M. L. 1997. GraphPad prism, data analysis, and scientific graphing. Journal of chemical information and computer sciences, 37, 411–412.

Wang, Q., Song, H., Cheng, H., Qi, J., Nam, G., Tan, S., Wang, J., Fang, M., Shi, Y., Tian, Z., Cao, X., An, Z., Yan, J., C Gao, G. F. 2020. Structures of the four Ig-like domain LILRB2 and the four-domain LILRB1 and HLA-G1 complex. Cellular & Molecular Immunology, 17, 966–975.

Wicher, K. B., Haneklaus, M., Poindron, A., Lopez-Yrigoyen, M., Seoane, C. R., Fransen, M., Payton, C., Guillame, S., Cassetta, L., Myatt, S., C Ries, C. 2023. 518 Discovery of MACO-355, a novel, first in mechanism ligand-blocking independent anti-LILRB1/2 monoclonal antibody for cancer therapy. Journal for ImmunoTherapy of Cancer, 11.

Willcox, B. E., Thomas, L. M., C Bjorkman, P. J. 2003. Crystal structure of HLA-A2 bound to LIR-1, a host and viral major histocompatibility complex receptor. Nat Immunol, 4, 913–9.

Wright, K. M., Dinapoli, S. R., Miller, M. S., Aitana Azurmendi, P., Zhao, X., Yu, Z., Chakrabarti, M., Shi, W., Douglass, J., Hwang, M. S., Hsiue, E. H.-C., Mog, B. J., Pearlman, A. H., Paul, S., Konig, M. F., Pardoll, D. M., Bettegowda, C., Papadopoulos, N., Kinzler, K. W., Vogelstein, B., Zhou, S., C Gabelli, S. B. 2023. Hydrophobic interactions dominate the recognition of a KRAS G12V neoantigen. Nature Communications, 14, 5063.

Yang, Z., C Bjorkman, P. J. 2008. Structure of UL18, a peptide-binding viral MHC mimic, bound to a host inhibitory receptor. Proceedings of the National Academy of Sciences, 105, 10095–10100.

Zeller, T., Münnich, I. A., Windisch, R., Hilger, P., Schewe, D. M., Humpe, A., C Kellner, C. 2023. Perspectives of targeting LILRB1 in innate and adaptive immune checkpoint therapy of cancer. Front Immunol, 14, 1240275.

Zemmour, J., C Parham, P. 1992. Distinctive polymorphism at the HLA-C locus: implications for the expression of HLA-C. Journal of Experimental Medicine, 176, 937–950.

Zhang, Y., Xu, Y., Wu, Q., Fu, X., Li, Y., C Li, A. 2025. Inhibitory leukocyte immunoglobulin-like receptors, subfamily B (LILRBs) in human diseases: structure, roles, mechanisms, and clinical applications. Theranostics, 15, 8222–8258.

Zhao, J., Zhong, S., Niu, X., Jiang, J., Zhang, R., C Li, Q. 2019. The MHC class I-LILRB1 signalling axis as a promising target in cancer therapy. Scandinavian Journal of Immunology, 90, e12804.

